# Systemic CYP3A inhibition by ritonavir enables selective targeting of hypoxic tumour cells by prodrugs of DNA-PK inhibitors

**DOI:** 10.64898/2026.02.12.705643

**Authors:** Cho Rong Hong, Benjamin D. Dickson, Lydia P. Liew, Way Wua Wong, Jagdish K. Jaiswal, Stephen M.F. Jamieson, Jacqueline M. Ross, Longjin Zhong, David M. Shackleford, William R. Wilson, Michael P. Hay

## Abstract

Hypoxic tumour cells are resistant to many forms of cancer therapy, particularly radiotherapy. Hypoxia-activated prodrugs (HAPs) can potentially address this problem through selective release of drugs (‘effectors’) in oxygen-deficient microenvironments, via metabolic reduction of a nitro(hetero)aromatic ‘trigger’ moiety. While many such HAPs show marked selectivity for hypoxia in cell culture, none have yet been approved for clinical use. Here, we report HAPs that release a novel inhibitor of the DNA repair enzyme DNA-dependent protein kinase (DNA-PK) which, like hypoxia, is a major contributor to radioresistance. These ether-linked HAPs provide hypoxia-dependent radiosensitisation in cell culture, but in mice systemic generation of the DNA-PK inhibitor is observed. Using *in vitro* hepatic metabolism models we demonstrate hypoxia-independent metabolic activation of HAP **4** via oxidation of its linker, which is mediated exclusively by CYP3A. We extend this finding to HAPs with other triggers, linkers and effectors. The clinically used CYP3A-specific inhibitor ritonavir suppressed hepatic metabolism of **4** under oxia without interfering with its hypoxia-dependent activation. In mice, ritonavir markedly enhanced oral bioavailability of the HAP, suppressed systemic formation of the DNA-PK inhibitor, and selectively radiosensitised HCT116 tumours but not the gastrointestinal tract in the radiation field. This combination offers the prospect of increasing the therapeutic ratio of DNA-PK inhibitor-mediated radiosensitisation in patients.

---

Hypoxia is a distinctive feature of the tumour microenvironment in a high proportion of cancers as a consequence of an imbalance between oxygen demand and its supply via spatially disorganised and temporally chaotic microvascular networks^1^. This lack of oxygen is a driver of genomic instability^2^, immune suppression^3-5^, metastasis^6^ and resistance to multiple therapeutic modalities^7^. These effects are mediated, in part, through genetic programmes triggered by oxygen sensors such as the 2-oxyglutarate-dependent dioxygenases that stabilise HIF transcription factors or act as epigenetic modifiers^7,8^. However, in the context of radiation therapy, hypoxia confers radioresistance primarily because oxygen reacts with radiation-induced DNA radicals to generate cytotoxic DNA double strand breaks (DSB) ^9-11^. Increasing adoption of highly hypofractionated radiotherapy protocols of short duration for cancer treatment^12^ reduces reoxygenation during treatment, increasing the need to address hypoxia-mediated radioresistance^13^.

One widely-studied approach uses hypoxia-activated prodrugs (HAPs) with nitro(hetero)aromatic ‘triggers’ (typically 2-nitroimidazoles) that are reduced by ubiquitous reductases under hypoxia, resulting in fragmentation of a linker to release an ‘effector’ drug selectively in tumours^14^. Many examples of nitro(hetero)aromatic HAPs with ether, carbonate or carbamate linkers have been shown to release their effectors selectively under hypoxia in cell culture^15-29^ but demonstration of significant *in vivo* efficacy and tumour selectivity in preclinical models is lacking, limiting clinical translation.

While hypoxia underpins environmental radioresistance, the major intrinsic resistance mechanism is DSB repair, particularly by the non-homologous end-joining (NHEJ) pathway which repairs ∼ 80% of radiation-induced DSBs in mammalian cells^30^. NHEJ is controlled by DNA-dependent protein kinase (DNA-PK) which mediates synapsis between DSB ends and recruits the NHEJ repair machinery^31-33^. As a consequence, genetic deletion of DNA-PK subunits (Ku70, Ku80 and catalytic subunit DNA-PKcs) ^34, 35^ and inhibitors of its kinase activity (DNA-PKi) ^36^ provide marked radiosensitisation, with a magnitude of effect similar to oxygen. However, DNA-PK mutation and/or DNA-PKi also radiosensitise critical normal tissues in the radiation field both in preclinical models^37, 38^ and in humans^39-41^. To address this limitation, we identified new HAPs that release DNA-PKi when reduced in hypoxic cells. This approach offers the potential for selective radiosensitisation of the most radioresistant cells in tumours whilst avoiding radiosensitisation of well-oxygenated normal tissues.

Our starting point is a recently described^42^ class of DNA-PKi exemplified by compound **1** (see Table S1 for compound structures). The analogous phenol **2** enables preparation of ether-linked HAPs **3**-**5** where the nitroimidazole trigger units mask DNA-PK inhibition in the prodrugs. In addition, conjugative metabolism of the phenolic OH (e.g. glucuronidation and sulfation) potentially acts as a ‘safety catch’ if the DNA-PKi effector is released in normal cells with high capacity for Phase II metabolism, such as enterocytes and hepatocytes. The canonical hypoxia-dependent bioreductive pathway for release of **2** from the preferred ether-linked HAP in this study, **4**, is illustrated in Fig. 1.

**Fig. 1.**
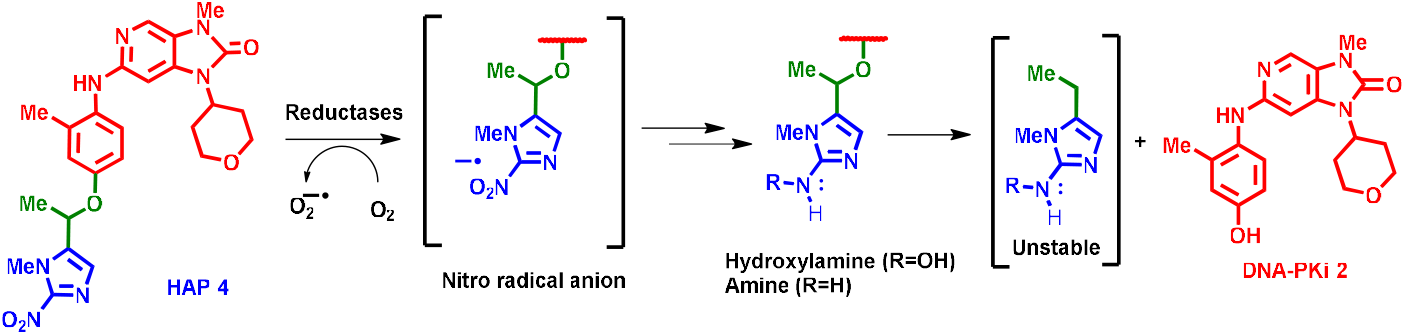
Proposed mechanism of bioreductive activation of hypoxia-activated prodrug 4. Reduction of the 2-nitroimidazole ‘trigger’ unit (blue) by ubiquitous one-electron reductases generates a nitro radical anion which is reoxidised by O_2_ in well-oxygenated cells, but under hypoxia is reduced further to the hydroxylamine or amine in which the linker (green) spontaneously fragments to release the DNA-PKi effector **2** (red).

However, we demonstrate a hypoxia-independent (and O_2_-dependent) pathway for activation of **4** and related HAPs in *in vitro* liver models and in mice. We show that this pathway is mediated by CYP3A and can be blocked by the specific CYP3A inhibitor ritonavir, which is widely used clinically as a component of Paxlovid™ for treatment of COVID-19^43,44^. In mice, dosing with ritonavir prior to HAP **4** provides enhanced oral bioavailability of HAP **4** and suppresses systemic release of the DNA-PKi, culminating in tumour radiosensitisation without enhancing radiation toxicity to the gastrointestinal tract, unlike DNA-PKi **2** which radiosensitises both tumour and normal tissues within the radiation field.

## Results

### Compound 2 is a potent DNA-PKi in cells with low glucuronidation activity

Compound **2**, similarly to the initial lead compound in the 6-anilino-imidazo[4,5-*c*]pyridin-2-one class^42^ **1**, is a potent DNA-PKi in biochemical assays (IC_50_ 13.1 nM) with high selectivity (170-3600-fold) relative to related PI3K kinases and mTOR (Table S1). It is also a DNA-PK-dependent radiosensitiser in cell culture regrowth assays, with radiosensitisation of wild-type HAP-1 cells but not its isogenic *PRKDC-*null derivative (Fig. 2a). In clonogenic survival assays, **2** showed concentration-dependent radiosensitisation in oxic human colorectal carcinoma HCT116 cultures (Fig. 2b) with potency similar to **1** (Fig. 2c). Given that our objective was to release **2** in hypoxic regions from a HAP, we then confirmed, in a panel of human tumour cell lines, that **2** at 1 μM (which was not cytotoxic alone; Table S2) radiosensitises under hypoxia as well as under oxia (Fig. 2d) with a similar magnitude under both conditions (Fig. 2e). Notably, sensitiser enhancement ratios at 10% survival (SER_10_) were similar in magnitude to the oxygen enhancement ratio (OER_10_) in most of these cell lines (Fig. 2e) demonstrating the importance of DNA-PK kinase activity in radiation resistance.

**Fig. 2.**
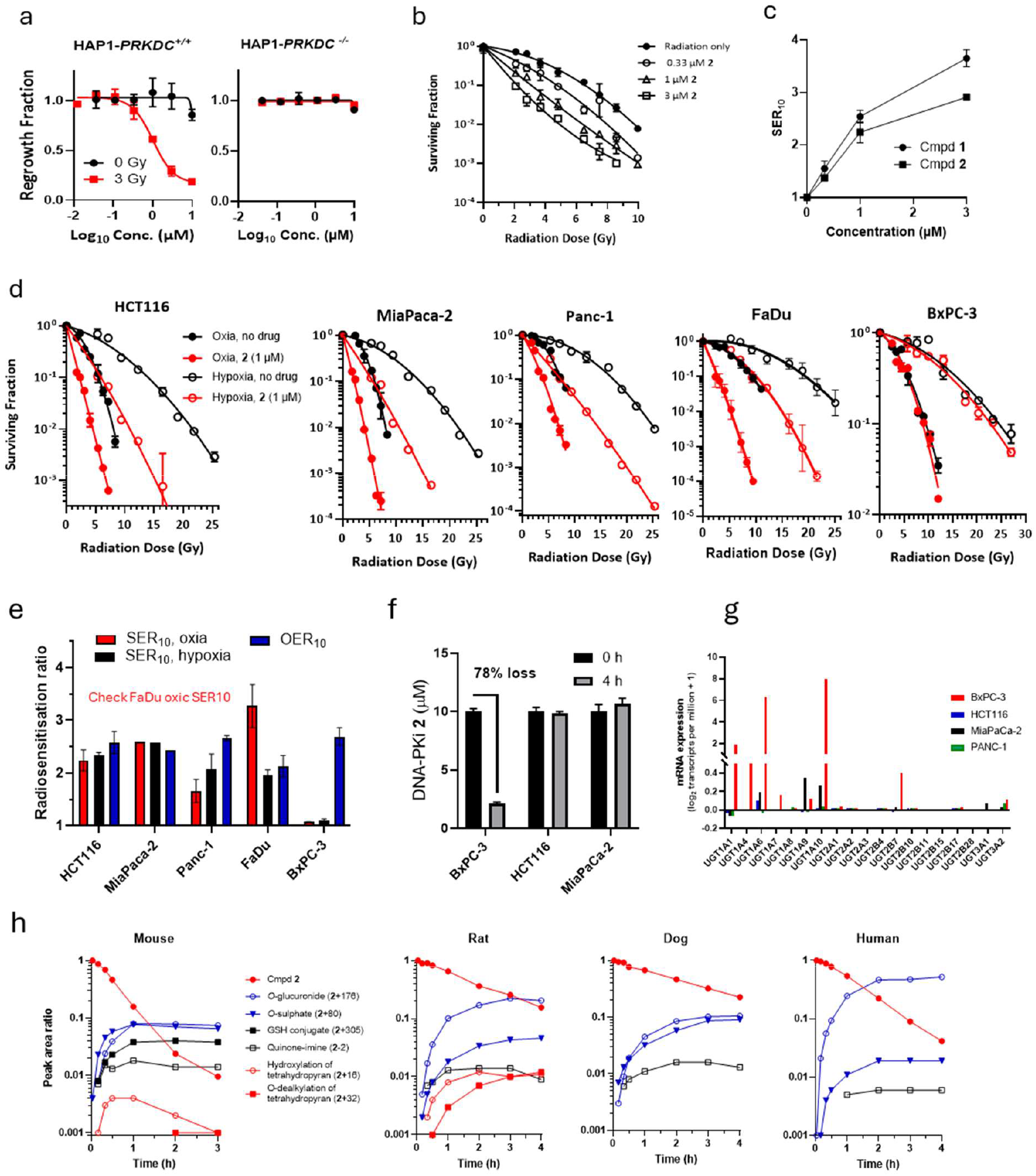
Radiosensitisation by phenolic DNA-PKi 2 in oxic and hypoxic cell cultures, and protection against radiosensitisation by glucuronidation. **a** Growth inhibition of *PRKDC*^*+/+*^ and *PRKDC*^*-/-*^ HAP-1 cell lines by **2** under oxia, with and without 3 Gy irradiation (IR). Mean and SEM from 3 experiments. **b** Radiosensitisation of oxic HCT116 cells by clonogenic assay with 18 h exposure to **2** post IR. Values are for 3 replicate cultures in a single experiment. **c** Sensitiser enhancement ratios at 10% survival (SER_10_) for 18 h exposure to **1** and **2** after IR in oxic HCT116 cultures. **d** Clonogenic survival curves for radiosensitisation of human tumour cell lines by **2** under oxia and hypoxia. Means are for 2-3 replicates in a representative experiment (all experiments in Table S2). **e** Comparison of radiosensitisation (SER_10_) by 1 μM **2** and 20% O_2_ (oxygen enhancement ratio, OER_10_) for the same cell lines. **f** Concentration of **2** in oxic cell cultures (10^6^ cells/mL). **g** mRNA expression of UGT isoforms in the cell lines retrieved from the DepMap 25Q3 dataset. **h** Metabolic loss of **2** (0.5 μM) in hepatocytes (2x10_6_ cells/mL) and formation of metabolites identified by LC-MS/MS. Quantitation is based on the peak area ratio relative to internal standard. Values in brackets refer to m/z changes of the metabolites relative to **2**.

An exception to the above was the pancreatic ductal adenocarcinoma BxPC-3, which was radiosensitised effectively by oxygen (OER_10_ = 2.69 ± 0.17), and by DNA-PKi **1**^42^, but not by **2** under either oxia or hypoxia (Fig. 2d,e). Therefore, we compared the metabolic stability of **2** in BxPC-3, HCT116 and MiaPaCa-2 cultures, demonstrating that it is rapidly metabolised only by the BxPC-3 line (Fig. 2f). Loss of **2** was accompanied by formation of a polar species, identified as the *O-*glucuronide by LC-MS/MS, only in BxPC-3 cultures (Fig. S1). Consistent with this, BxPC-3, but not cell lines that are radiosensitised by **2**, highly express UDP-glucuronosyltransferases UGT1A1, -1A6 and -1A10 (Fig. 2g), all of which are known to efficiently conjugate phenols^45^. Thus, cells with high activity of phenol-conjugating UGTs can metabolise **2** rapidly enough to become resistant to this radiosensitiser.

While few human tumours show high UGT expression^46-48^, this detoxification pathway may have potential to protect normal tissues from radiosensitisation by **2**, particularly via metabolic clearance in the liver, should the DNA-PKi efflux from tumours or be generated in non-tumour tissues. Thus, we evaluated clearance of **2**, and characterised its metabolites, in hepatocyte suspension cultures by LC-MS/MS (Fig. 2h). Intrinsic clearance (units μL/min/10_6_ cells) was highest in mouse (16.3; 95% CI 15.1-17.5) followed by human (6.9; 6.6-7.2), rat (3.9; 3.7-4.2) and dog (3.1; 2.9-3.3). Metabolites, identified by collision-induced dissociation (Fig. S2) and high mass resolution m/z values, provided the inferred metabolic scheme in Fig. S3 which has three main branches: conjugation of the phenol to the *O-*glucuronide (and to a lesser extent the *O*-sulfate), which were the prominent metabolites (based on ion counts) in all species (Fig. 2h); oxidation of the tetrahydropyran ring in rodents; and oxidation to a quinone-imine intermediate (subsequently shown to be CYP-mediated; Fig. S5) that adds glutathione in rodents. Notably, the *O*-glucuronide was a much more prominent metabolite in human hepatocytes than in the other species (Fig. 2h).

### Hypoxia-selective metabolic activation and radiosensitisation by HAPs *in vitro*

Metabolism of the three ether-linked HAPs (**3**-**5**) was assessed in hypoxic HCT116 cultures. The prodrugs with 2-nitroimidazole triggers (**3,4**) were depleted within 4 h, with formation of the DNA-PKi effector **2** (Fig. 3a), consistent with the proposed bioreductive activation mechanism (Fig. 1). In contrast, prodrug **5**, used as a negative control because its lower reduction potential 5-nitroimidazole trigger^15^, is less readily reduced by hypoxia-dependent mammalian reductases^49^, was metabolically stable with no formation of **2** (Fig. 3a). The kinetics of metabolism of **3** in hypoxic HCT116 cultures confirmed loss of prodrug and formation of **2**, with a higher rate in an isogenic HCT116 line overexpressing cytochrome P450 oxidoreductase (POR) (Fig. 3b). This supports the proposed mechanism (Fig. 1) given that POR is known to catalyse hypoxia-dependent one-electron activation of 2-nitroimidazole-linked HAPs^50, 51^.

**Fig. 3.**
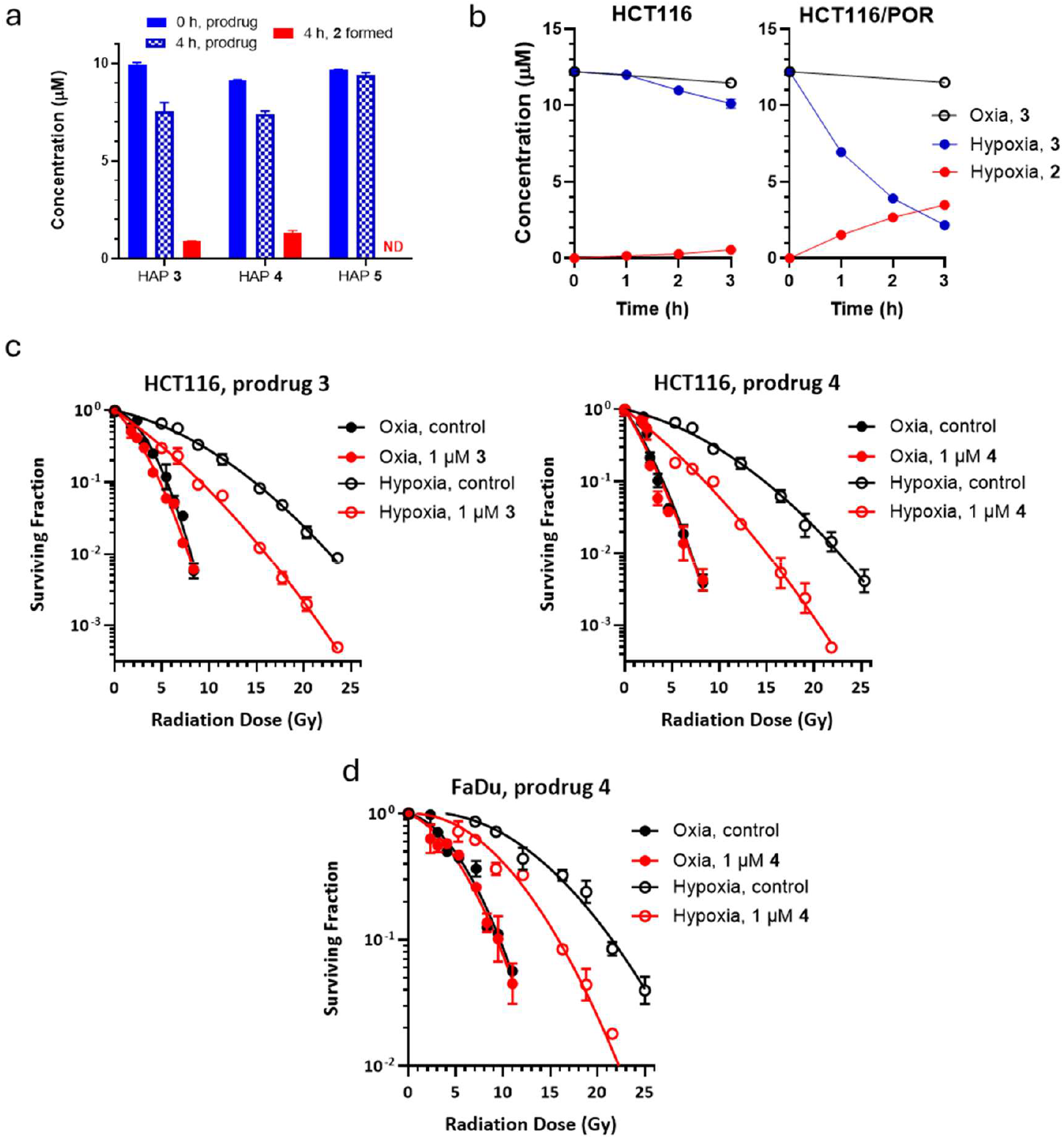
Hypoxia-dependent metabolic activation and radiosensitisation by HAPs in cell culture. **a** Metabolism of prodrugs **3-5** and formation of DNA-PKi **2** after 4 h hypoxia in HCT116 cultures (10_6_ cells/mL). **b** Time course of metabolism of HAP **3** and formation of DNA-PKi **2** in an isogenic HCT116 and POR-overexpressing (HCT116/POR) cell line pair (10^6^ cells/mL) under oxia and hypoxia. Means of 3 replicates (SEMs smaller than the plotted points). DNA-PKi **2** was not detected under oxia in either line. **c** Clonogenic survival curves for radiosensitisation of HCT116 by **3** and **4** under oxia and hypoxia. Representative experiments each with 2-4 replicates (see also Table S3). **d** Radiosensitisation of FaDu cells by HAP **4** under oxia and anoxia, as for c.

Consistent with this hypoxic activation, and the lack of DNA-PK inhibition by the prodrugs under oxia (Table S1), HAPs **3** and **4** at 1 μM radiosensitised hypoxic HCT116 cells while neither prodrug had any effect in oxic cultures (Fig. 3c). This selective radiosensitisation under hypoxia was confirmed for prodrug **4** in FaDu hypopharyngeal squamous carcinoma cells (Fig. 3d). SER_10_ values for replicate experiments were 1.51±0.07 for **3** and 1.36±0.14 for **4** in HCT116 cultures, and 1.35±0.02 for **4** in FaDu, with SER_10_ of only 1.04-1.05 for these compounds under oxia (Table S3). As expected, given its lack of bioreductive activation, prodrug **5** provided little radiosensitisation of HCT116 under hypoxia (Table S3).

### Tumour xenograft radiosensitisation by the HAPs is at least partially mediated by systemic generation of DNA-PKi 2

We tested the ability of **2** and its prodrugs **3-5** to radiosensitise tumours, comparing the compounds at equimolar doses and measuring the number of surviving clonogens 18 h after treatment (Fig. 4a). In the well-characterised^52, 53^ UT-SCC-74B tongue squamous cell carcinoma xenograft model, the DNA-PKi **2** and its 2-nitroimidazole HAPs **3** and **4** had no effect as single agents but provided highly significant radiosensitisation (Fig. 4b). Compounds **2**-**4** were also effective in radiosensitising HCT116 tumours but, surprisingly, so was the negative control 5-nitroimidazole **5** (Fig. 4c). This cast doubt on the exclusive role of bioreduction, leading us to evaluate biodistribution of the prodrugs (Fig. 4d), and **2** as a metabolite (Fig. 4e), in mice with HCT116 tumours. After a single i.p. dose at 90 μmol/kg, levels of HAPs **3-5** and metabolite **2** decreased rapidly in plasma and tumour, while prodrug levels fell more slowly in normal tissue (liver, lung, spleen) resulting in very low tumour/normal tissue ratios (0.01-0.001; (Fig. 4d). Importantly, DNA-PKi **2** was a prominent metabolite in all tissues; tumour/normal tissue ratios (range 0.25-5.6) were much higher than for the prodrugs, suggesting possible intra-tumour activation, but the presence of **2** as a metabolite in plasma and all normal tissues strongly suggested significant systemic exposure to the DNA-PKi as a result of extra-tumoural metabolism of the prodrugs. Subsequent experiments demonstrated that **2** is also formed at high levels from **4** in non-tumour bearing mice (e.g. Fig. 8a), supporting this conclusion.

**Fig. 4.**
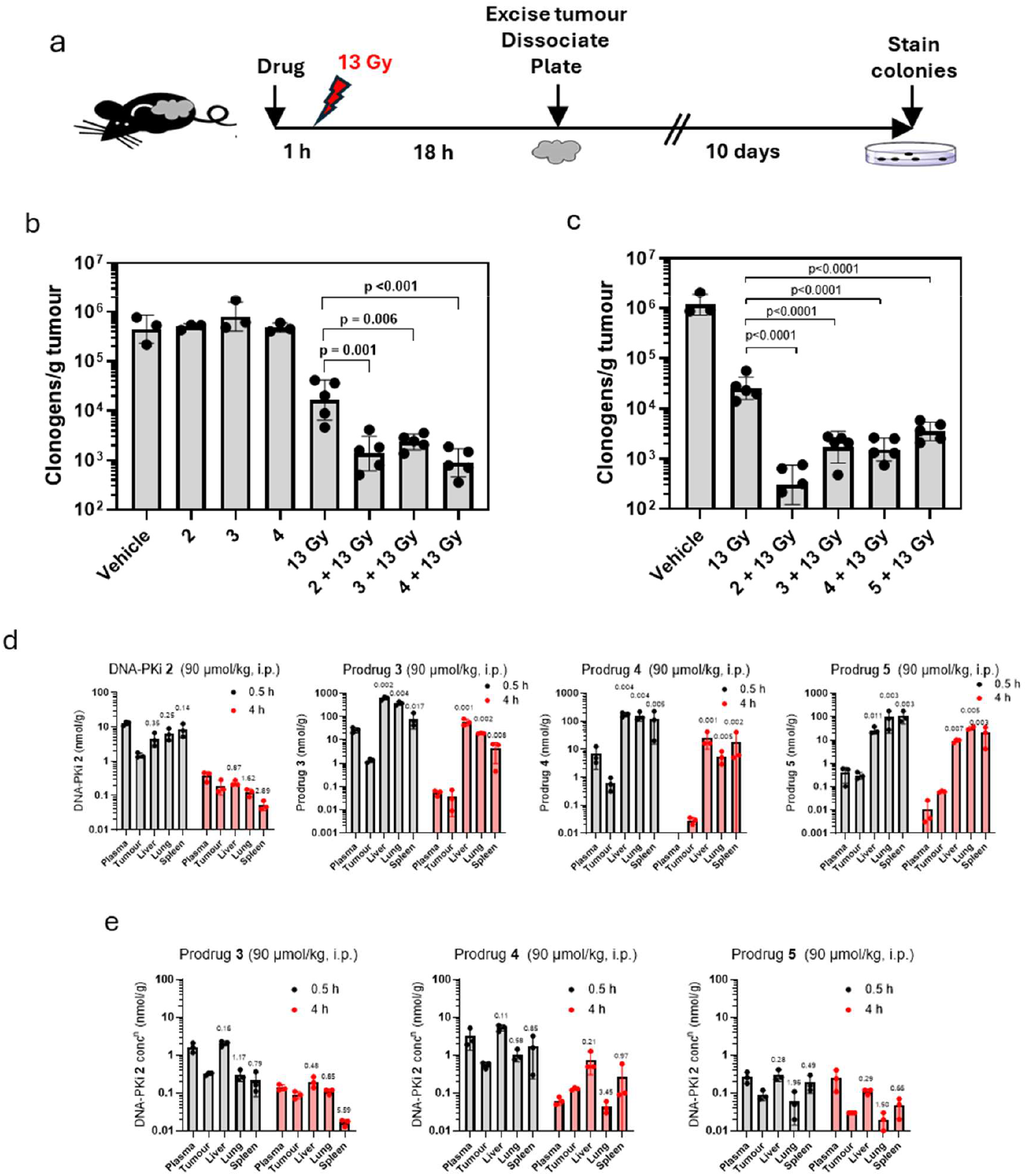
Radiosensitisation of human tumour xenografts by DNA-PKi 2 and its HAPs, and their biodistribution in tumour and normal tissues. **a** Schematic of the *ex vivo* assay for surviving tumour clonogens after treatment of subcutaneous xenografts in NIH-III mice. **b** Surviving tumour clonogens 18 h after treatment of UT-SCC-74B squamous cell carcinoma xenografts with DNA-PKi **2** or HAPs (2x90 μmol/kg, i.p.) 0.25 h before and 2.75 h after IR (13 Gy) or sham irradiation. **c** Clonogenic survivors after treatment of HCT116 xenografts as for b. No colonies recovered from 1 of 5 tumours. **d** Plasma and tissue concentrations of DNA-PKi **2** and prodrugs **3**-**5**, 0.5 and 4 h after single i.p. doses (90 μmol/kg; 3 animals/group). **e** Plasma and tissue concentrations of **2** as a metabolite from prodrugs **3**-**5**, in the same animals as for d. Numbers above the bars in d and e are average tumour/normal tissue concentration ratios.

### The HAPs are metabolically activated by CYP-mediated linker oxidation in oxic liver S9 and hepatocytes

The metabolic instability of the prodrugs in mice led us to test their metabolism in oxic mouse liver S9 preparations supplemented with 1 mM NADPH. DNA-PKi **2** was metabolically stable while all three prodrugs showed rapid NADPH-dependent metabolism (rates **5** > **4** > **3**) with formation of **2** as a metabolite (Fig. S4); the high rate for **5** confirmed that this oxic metabolism is not bioreductive.

We inferred that the likely mechanism is CYP-mediated oxidation of the linker and hypothesised that this would generate the unreduced 2-nitroimidazole ketone metabolite **6** from **4** (Fig. 5a). To test this, we synthesised **6** and confirmed its formation from **4** in oxic mouse liver S9 by mass spectrometry (multiple reaction monitoring) and retention time (Fig. 5b). The cofactor dependence (NADPH > NADH) was also consistent with CYP-mediated metabolism, and CYP-dependence was confirmed by the quantitative inhibition of loss of **4** and formation of **2** by a pan-CYP inhibitor (CYPi) cocktail comprising 1-aminobenzotriazole and tienilic acid (Fig. 5c). The kinetics of formation of **6** was similar, but not identical, to that of **2** (Fig. 5c), likely because the ketone metabolite undergoes further NADPH-dependent metabolism at a similar rate to **4** (data not shown). However, estimation of **6** by electrospray ionisation for LC-MS/MS was only semi-quantitative as the detection sensitivity (ion counts per pmol on column) was only 0.4% of that for **2**.

**Fig. 5.**
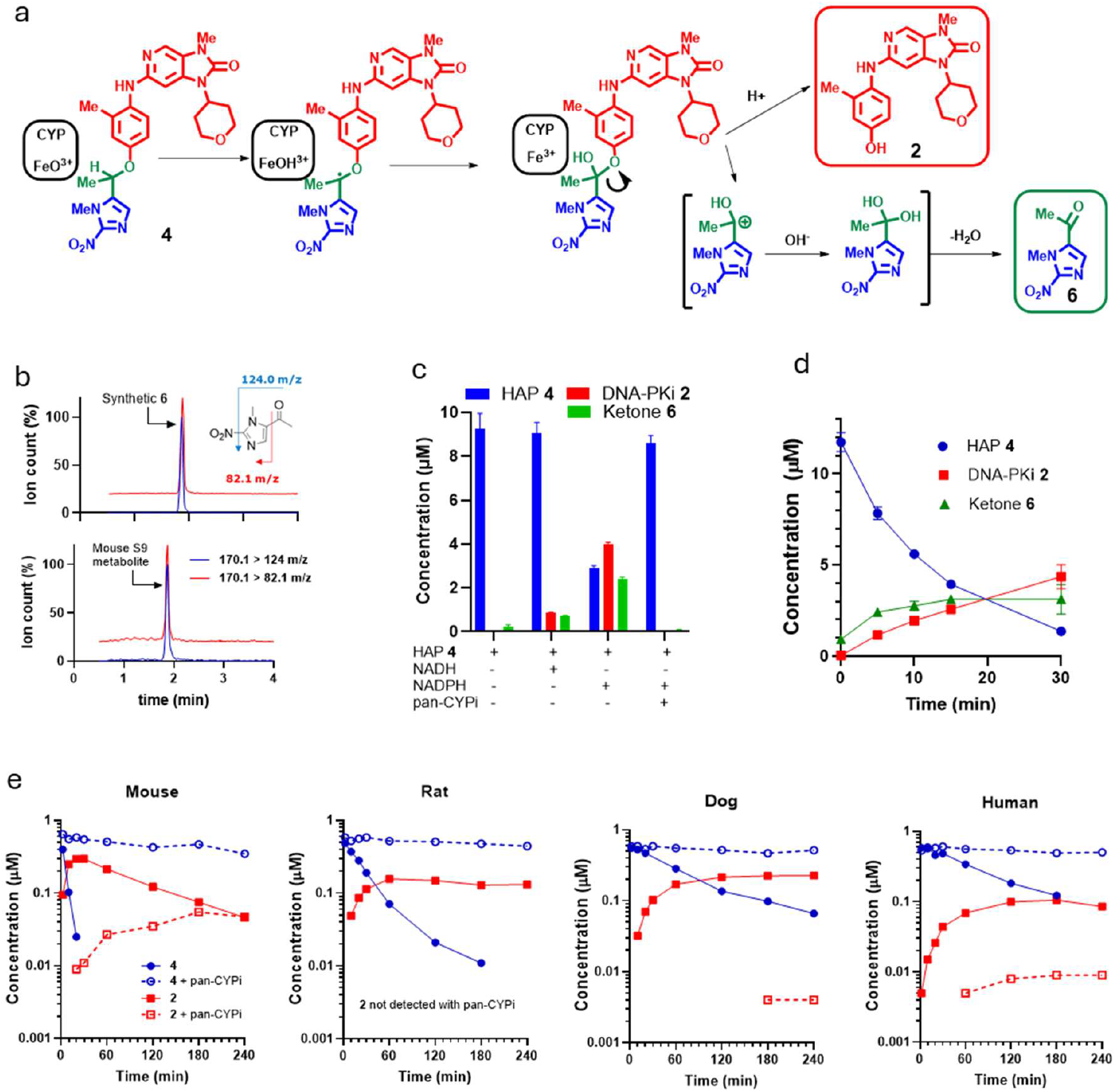
HAP 4 undergoes CYP-mediated fragmentation to release DNA-PKi 2 in oxic liver S9 and hepatocytes. Identification and quantitation by LC-MS/MS. **a** Proposed scheme for CYP-mediated oxidation of the linker in HAP **4**, generating **2** and the unreduced 2-nitroimidazole ketone **6. b** Identification of **6** as a metabolite of HAP **4** in oxic mouse liver S9 (1 mg protein/mL, 1 mM NADPH, 30 min incubation) **c** Metabolism of **4** in oxic mouse liver S9 (1 mg protein/mL, 30 min) with 1 mM NADH or NADPH, and a pan-CYP inhibitor (pan-CYPi) cocktail (1 mM 1-aminobenzotriazole + 15 μM tienilic acid). Means and SEM for 4 replicates. **d** Kinetics of metabolism of **4** under the same conditions as in c. Means and SEM for 3 replicates. **e** Metabolic loss of **4** (initial concentration 0.5 μM) and formation of **2** in oxic hepatocytes (5x10^5^ viable cells/mL) with and without pan-CYPi.

CYP dependence of metabolism of **4** in oxic hepatocyte cultures was further evaluated across four species (mouse, rat, dog and human; Fig. 5e). In all species **4** was consumed rapidly with intrinsic clearances mouse > rat > dog > human (quantified in Table S4). DNA-PKi **2** was the major metabolite in all species, with yields 32-84% of prodrug loss (Table S4), while pan-CYP inhibition markedly reduced both loss of **4** and formation of **2** (Fig. 5e, Table S4). The full profile of metabolites in hepatocytes (Fig. S5 and Fig. S6) includes the 2-nitroimidazole ketone **6**, a minus-2Da metabolite of **4 (4**-2) and downstream metabolites from **2**.

### Oxic metabolism of HAP 4 is mediated by CYP3A and inhibited by ritonavir

To identify CYP isoforms responsible for release of **2** from HAP **4**, the metabolism of **2** and **4** was evaluated in oxic human liver microsomes with NADPH in the absence and presence of isoform-specific inhibitors^54^. When incubated in the absence of CYP inhibitors DNA-PKi **2** was stable (data not shown), while **4** was rapidly metabolised to **2** (Fig. 6a) with a first order rate constant of 0.121±0.007 min^-1^. The CYP3A-specific inhibitor ketoconazole reduced that rate constant to 0.005±0.007 min (fraction of metabolism by CYP3A = 0.96) with formation of **2** below the level of detection (Fig. 6a). None of the other inhibitors in this reaction phenotyping study inhibited loss of **4** or formation of **2** (Fig. S7).

**Fig. 6.**
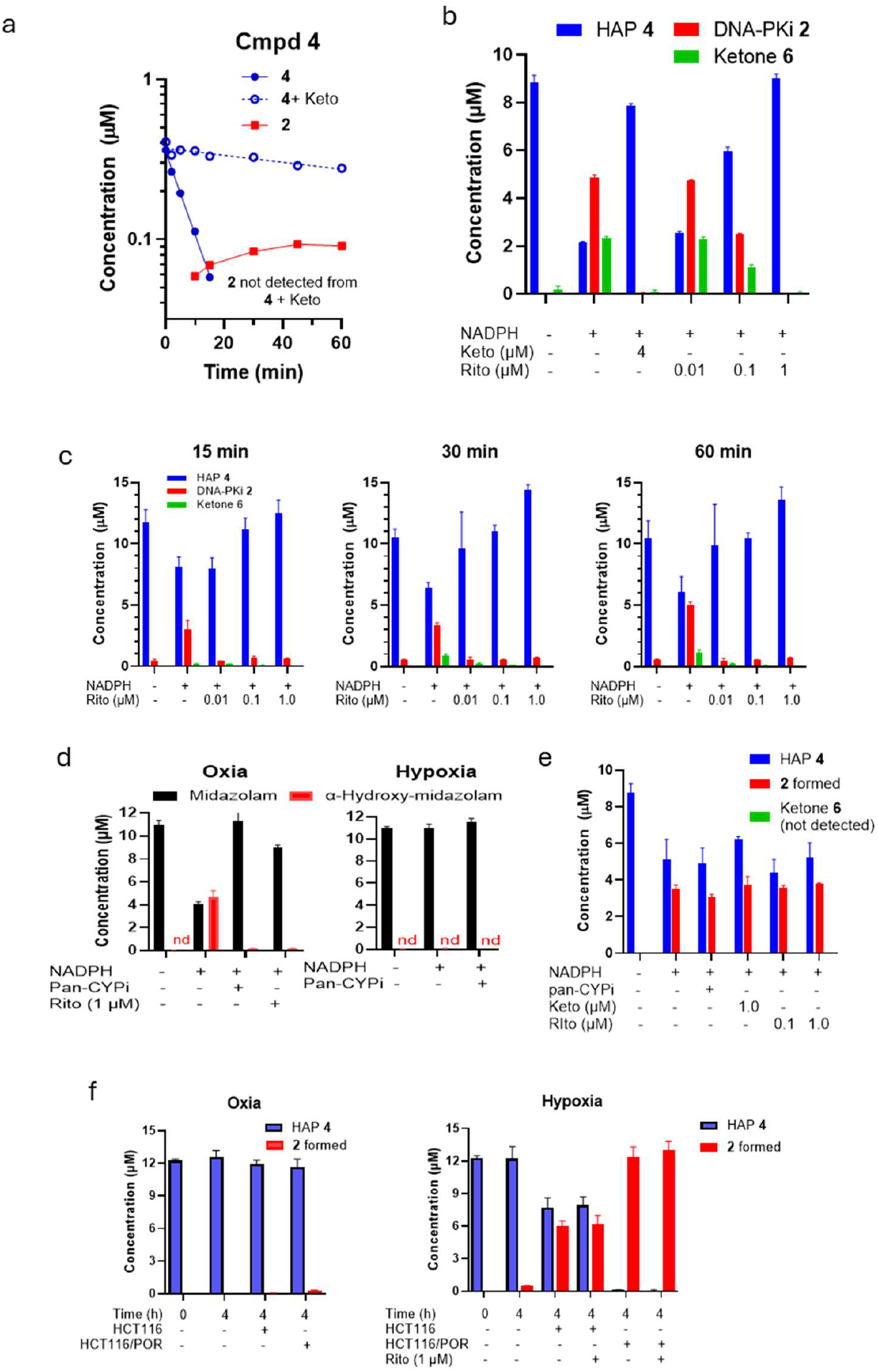
Oxic metabolism of HAP 4 in human liver microsomes and mouse liver S9 is dependent on CYP3A. **a** Metabolism of **4** in human liver microsomes (1 mg protein/mL) with 1.3 mM NADPH ± CYP3A-specific inhibitor ketoconazole (Keto; 1.5 μM). **b** Loss of **4** and formation of metabolites **2** and **6** in S9 (1 mg protein/mL, 30 min) with 1 mM NADPH ± Keto or ± Rito (ritonavir) (added 5 min prior to **4**). Mean and SEM for 4 replicates. **c** As for b, but addition of Rito 20 min prior to **4**. Mean and SEM for 3 replicates. **d** Metabolism of midazolam in mouse liver S9 (1 mg protein/ml, 15 min) with 1 mM NADPH ± Rito (20 min preincubation; left) or hypoxic (right) conditions. Mean and SEM for 4-9 replicates pooled from two experiments. **e** Lack of effect of pan-CYP inhibition or CYP3A inhibition (Rito, 20 min preincubation) on metabolism of **4** in hypoxic S9 (1 mg protein/mL, 30 min). Mean and SEM for 3 replicates. **f** Metabolism of **4** in oxic and hypoxic cultures of HCT116 and HCT116/POR (10^6^ cells/mL) with Rito (1 μM) added 20 min before the prodrug. Mean and SEM for 3-4 cultures.

Ritonavir, based on its use as a co-drug in Paxlovid™ for treatment of COVID-19^43,44^, is a well-validated CYP3A inhibitor for clinical use^55^. We therefore tested inhibition of metabolism of **4** in oxic mouse S9 by ketoconazole (4 μM) and ritonavir (1 μM), both of which fully inhibited loss of **4** and formation of **2**, with ritonavir providing partial inhibition at 0.1 μM when added 5 min prior to **4** (Fig. 6b). Inhibition was more effective when ritonavir was added 20 min prior (Fig. 6c), consistent with its known time-dependent irreversible inhibition of CYP3A^56^.

We then tested whether ritonavir would interfere with the desired hypoxia-dependent activation mechanism. NADPH-dependent oxidation of the CYP3A probe midazolam^57^ to α-hydroxy-midazolam in oxic mouse liver S9, was blocked by pan-CYP inhibition and by ritonavir as expected, but no metabolism was observed under hypoxia (Fig. 6d) demonstrating that the concentration of the O_2_ co-substrate is sufficiently low in our hypoxic S9 model to preclude CYP3A activity. Under hypoxia, the rate of metabolism of **4** to **2** was similar to that under oxia, notably without formation of the ketone metabolite **6**, and was not inhibited by either pan-CYPi or ritonavir (Fig. 6e) demonstrating that the oxic and hypoxic activation pathways are distinct. Similarly, we confirmed that the hypoxia-dependent bioreductive metabolism of **4** in HCT116 and HCT116/POR cells is not inhibited by ritonavir (Fig. 6f).

### CYP3A also mediates oxic activation of other ether- and carbamate-linked HAPs

To assess the generality of the CYP3A dependence of metabolic activation, we evaluated the metabolism of other trigger-linker-effector HAPs in oxic mouse liver S9 (Fig. 7). As for **4**, activation of prodrugs of **2** with a methylene linker (**3**; Fig. 7a), a 5-nitroimidazole trigger (**5**; Fig. 7b) or a 2-nitrothiophene trigger (**7**; Fig, 7c) was completely inhibited by pan-CYPi, ketoconazole and ritonavir. Similar CYP3A dependence was demonstrated for a carbamate-linked HAP of DNA-PKi **1** (HAP **8**; Fig. 7d), as judged by loss of prodrug, although its DNA-PKi metabolite was metabolised faster than **8** so formation of **1** was less informative. The 2-nitroimidazole ether-linked HAP with an unrelated effector (an inhibitor of protein kinase R-like endoplasmic reticulum kinase; PERKi), **9**^21^, was also metabolised to release the PERKi **10** in oxic mouse liver S9, and this was again inhibited by the CYP inhibitors (although significant further metabolism of **10** again complicated quantitation of effector release). We conclude that CYP3A is involved in off-target activation of a wide range of nitroheterocyclic HAPs with differing triggers, linkers and effectors.

**Fig. 7.**
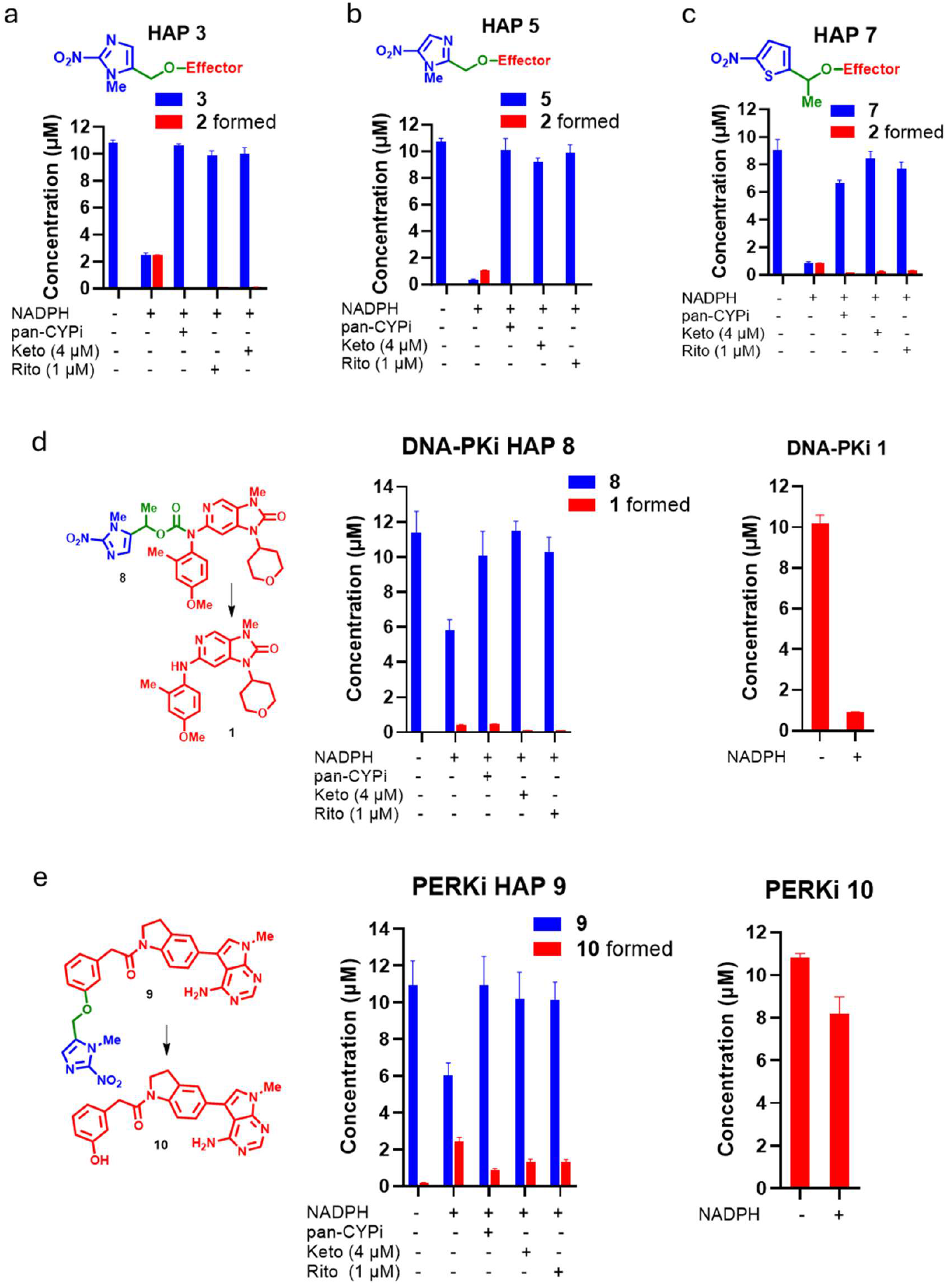
CYP3A-mediated metabolism of other ether-linked and carbamate-linked HAPs in oxic mouse liver S9. 1 mM NADPH, with preincubation with Rito (when used) for 20 min. **a** Metabolism of the 2-nitroimidazole HAP **3** (1 mg protein/mL, 30 min incubation). **b** 5-Nitroimidazole HAP **5** (1 mg protein/ml, 15 min). **c** 2-Nitrothiophene HAP **7** (1 mg protein/mL, 30 min). **d** Carbamate-linked HAP **8** (left) and its DNA-PKi effector **1** (right) (2 mg protein/mL, 60 min). **e** 2-Nitroimidazole HAP **9** (left) and its PERKi effector **10** (right) (1 mg protein/mL, 60 min).

### Ritonavir enhances oral bioavailability of HAP 4 in mice and suppresses systemic release of 2

The finding that CYP3A mediates oxic activation of HAPs led us to test whether ritonavir can suppress systemic exposure to **2** following HAP **4** administration in mice. Oral ritonavir (25 mg/kg, 1 h before i.p. **4** at 90 μmol/kg (46 mg/kg)) strongly enhanced levels of **4** and suppressed the ratio of **2**/**4** at 0.5 and 4 h in plasma and in the three normal tissues (liver, lung, spleen), particularly by 4 h. Much less suppression of **2** by ritonavir was seen in HCT116 tumours by 4 h (Fig. 8a) suggesting that intra-tumour activation of **4** to **2** becomes significant when systemic metabolism is suppressed by ritonavir.

**Fig. 8.**
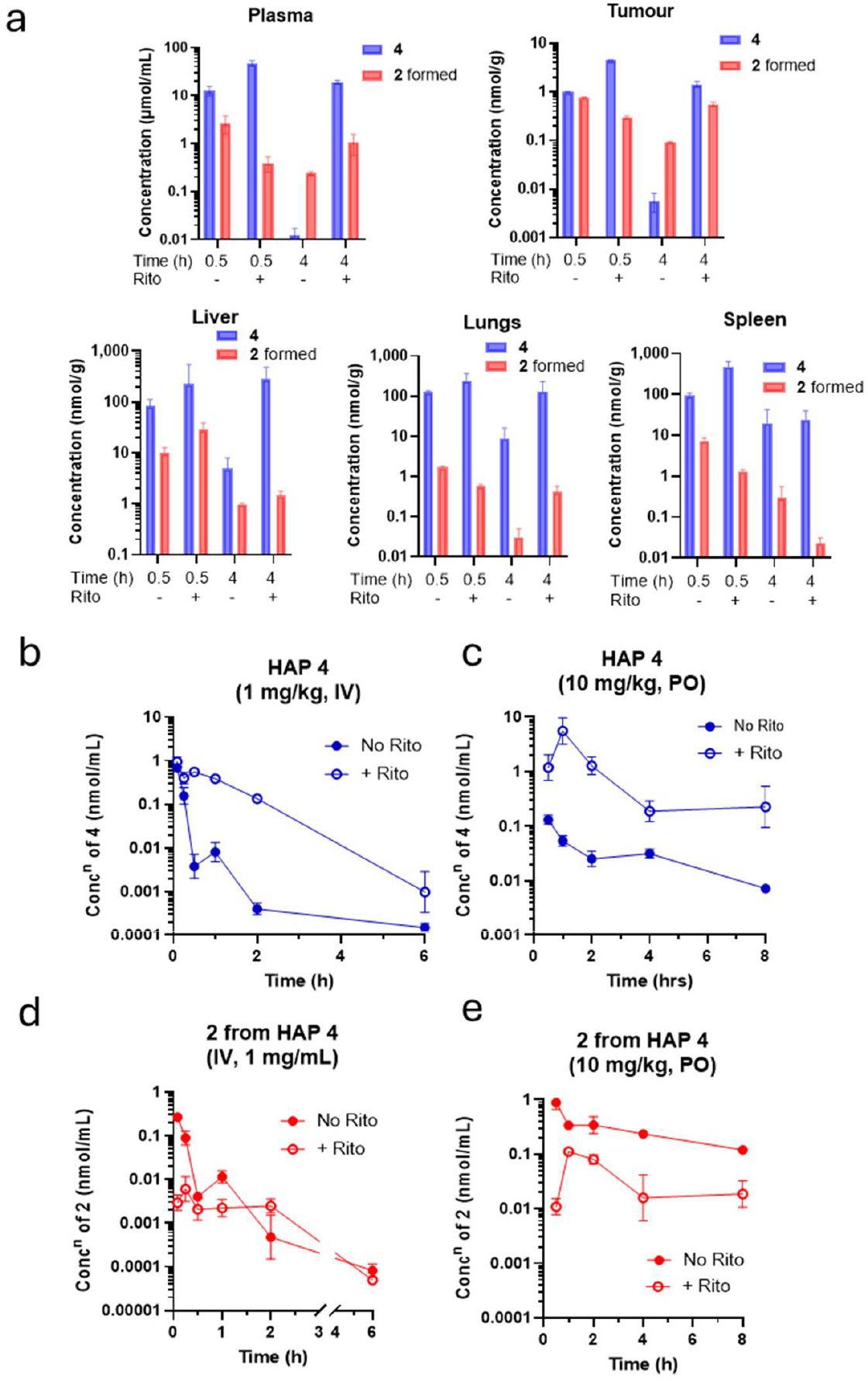
Effect of ritonavir (Rito) on tissue distribution, plasma pharmacokinetics (PK) and oral bioavailability of HAP 4 and its DNA-PKi metabolite 2. Rito was dosed orally by gavage at 25 mg/kg body weight, 1 h prior to **4**. Values are geometric means and SEM for 3 animals/group. **a** NIH-III mice with HCT116 tumours were dosed i.p. with **4** at 90 μmol/kg (46 mg/kg) with or without Rito. **b**-**e** Plasma PK in non-tumour-bearing CD1 Swiss mice, with or without Rito. **b** HAP **4** dosed i.v. at 1 mg/kg. **c** HAP **4** dosed orally by gavage at 10 mg/kg. **d** Metabolite **2** in the mice dosed i.v. with **4** at 1 mg/kg in b. **e** Metabolite **2** in the mice dosed p.o. with **4** at 10 mg/kg in c.

The dramatic reduction in systemic **2** after ritonavir led us to quantify effects of the CYP3A inhibitor on plasma pharmacokinetics (Fig. 8b-e). Exposure to **4**, based on the area under the plasma concentration-time curve for 0-6 h (AUC_0-6h_), was greatly increased by ritonavir from 0.136 to 1.08 μmol.h/L after i.v. dosing at 1 mg/kg, and from 0.265 to 10.1 μmol.h/L after oral dosing at 10 mg/kg. Consequently, oral availability of **4** was increased by ritonavir from 19% to 94%, while plasma concentrations of **2** from oral **4** were strongly suppressed (Fig. 8e) consistent with Fig. 8a.

Given the high oral bioavailability of **4** with ritonavir (**4**+Rito), we tested tolerability of the combination at higher doses of **4**. No body weight loss or clinical signs of toxicity were observed with oral **4**+Rito up to the highest dose tested (417 mg/kg of **4**; Fig. S8). Intraperitoneal dosing with **2** was also well tolerated but was limited to 100 mg/kg because of its solubility in the formulation (Fig. S8).

### HAP 4 with ritonavir provides more selective tumour radiosensitisation than administration of DNA-PKi 2

The above pharmacokinetic analysis is based on whole tissue averages and may substantially underestimate concentrations of **2** in radioresistant hypoxic zones. We therefore turned to pharmacodynamic endpoints to assess whether **4**+Rito provides a therapeutic advantage over **2** for tumour radiosensitisation. The DNA-PKi **2** radiosensitised HCT116 tumours significantly across the range 10-100 mg/kg, with no activity as a single agent at the highest dose (Fig. 9a). Oral **4**+Rito alone also had no antitumour activity, but significantly radiosensitised the tumours at 100 and 400 mg/kg of **4** (Fig. 9b).

**Fig. 9.**
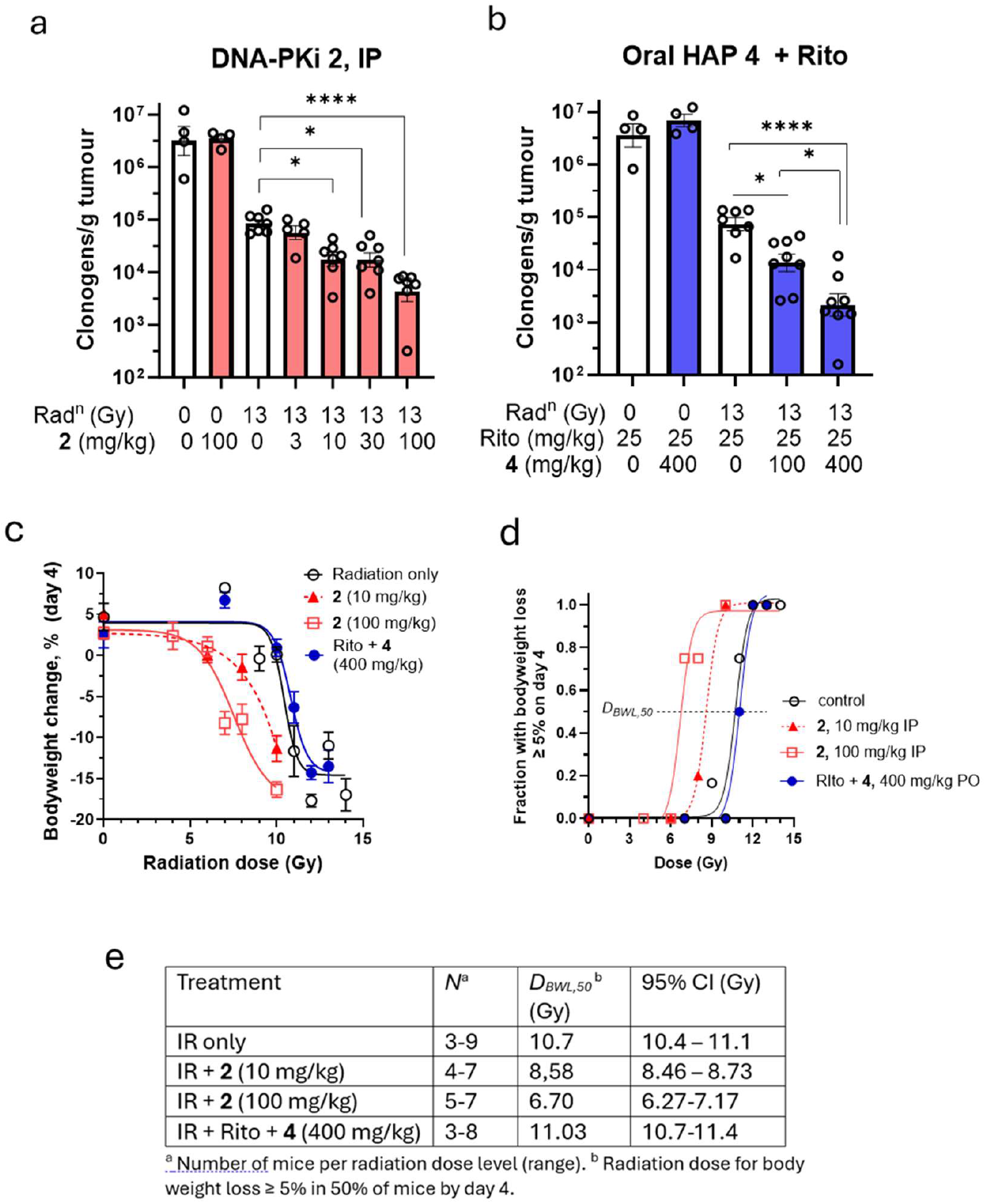
Radiosensitisation of HCT116 tumour xenografts and bodyweight loss by i.p. DNA-PKi 2 or oral 4+Rito in mice. **a,b** Surviving tumour clonogens 18 h after treatment. Geometric mean and SEM for 4 animals/group without radiation and 5-8 animals/group with 13 Gy irradiation. **a** DNA-PKi **2** i.p. 15 min before radiation. **b** HAP **4** p.o. 1 h after Rito (25 mg/kg, p.o.) and 0.5 h before radiation. **c** Radiation dose dependence of bodyweight change on day 4 after whole-body irradiation relative to pre-treatment, as a surrogate of gastrointestinal tract radiation toxicity, measured at 8-9 am to minimise diurnal variation, in female CD1 Swiss mice. Mice were dosed i.p. with **2** and orally with **4**+Rito. **d** Radiation dose dependence of the fraction of animals with body weight loss ≥ 5% relative to pre-treatment. Lines are 3 parameter logistic regressions. **e** Numbers of animals per group, mean *D*_*BWL,50*_ values and 95% confidence intervals for the bodyweight change study.

We then compared the two approaches with respect to radiosensitisation of the gastrointestinal tract, for which early bodyweight loss is a well-validated surrogate in both mice and humans^58^, and is acutely dose-limiting in many radiotherapy settings^59, 60^. The DNA-PKi **2** was non-toxic alone, as expected, but caused a marked dose-dependent reduction in bodyweight by day 4 after whole body irradiation (Fig. 9c). In contrast, oral **4** at 400 mg/kg, a dose providing equivalent tumour radiosensitisation to **2** at 100 mg/kg, did not reduce bodyweight in mice pre-treated with ritonavir. Logistic regression (Fig. 9d) confirmed the lack of change in the radiation dose for ≥5% bodyweight loss in 50% of mice (*D*_*BWL,50*_) for **4**+Rito while *D*_*BWL,50*_ was significantly depressed by **2** at both 10 and 100 mg/kg (Fig. 9e).

To more directly assess radiation toxicity in the gastrointestinal tract we employed labelling of regenerating (S-phase) stem cells, with 5-ethynyl-2’-deoxyuridine (EdU), in the small intestine (ileum) and oral mucosa (tongue) on day 4 after irradiation as previously^37^. Representative images show a rebound in EdU+ cells in the ileum after 10 Gy, with proliferating ‘transit amplifying’ cells extending up the villus. This response was markedly suppressed by 100 mg/kg **2** but not by **4**+Rito at the HAP dose (400 mg/kg) providing equivalent tumour radiosensitisation (Fig. 10a). Similarly, S-phase cells in the oral mucosa were strongly suppressed by 100 mg/kg **2** but not by 400 mg/kg **4**+Rito (Fig.10b). Quantitation of EdU+ nuclei in the images from all mice, including drug-only controls, confirmed these findings with highly significant suppression by 10 Gy +100 mg/kg **2** relative to 10 Gy alone (Fig. 10c). The increase in the S-phase population after 10 mg/kg **2** and 400 mg/kg **4** with 10 Gy was not statistically significant but this regenerative response could be detected more sensitively by measuring the mean thickness of the band of EdU+ve nuclei extending up the villi (Fig. 10d) and suggests that both 10 mg/kg **2** and 400 mg/kg **4**+Rito cause some increase in DNA damage by 10 Gy IR. In the oral mucosa this repopulation was not seen after 10 Gy, but again the number of S-phase cells was suppressed by 100 mg/kg **2** but not HAP **4**+Rito (Fig. 10e).

**Fig. 10.**
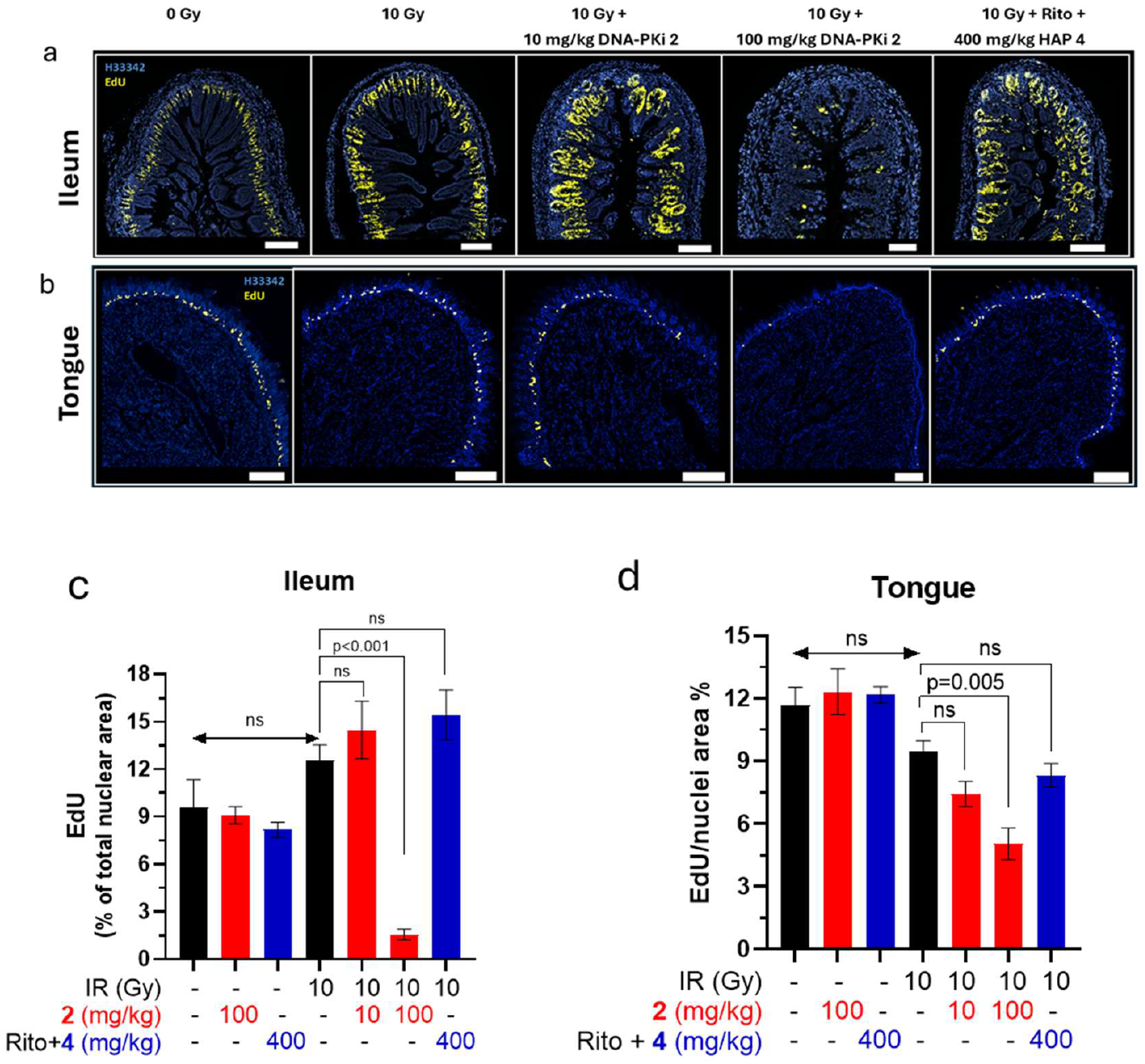
Effect of DNA-PKi 2 and HAP 4 + Rito on regenerating stem cells in the gastrointestinal tract after 10 Gy whole body irradiation. S-phase cells were labelled by dosing mice i.p. with EdU 2 h before euthanasia, and sections were stained for EdU and total nuclei with H33342. **a, b**: representative images for transverse sections of ileum (a) and longitudinal sections of tongue (b). Scale bars 200 μm. **c, d** Quantitation of EdU+ nuclei as the area ratio EdU/H33342 on thresholded binary images for ileum (c) and tongue (d) for all images (Fig. S9 and Fig. S10, N=4 mice/group), using one ileum section and 2 tongue sections from each mouse.

## Discussion

Targeting treatment-resistant hypoxic cells in tumours has been a major objective in radiation oncology since the 1970s^61, 62^. However, selective elimination of hypoxic cells has proven elusive, and no hypoxia-targeted agent has yet achieved regulatory approval for use in cancer treatment. Multiple challenges with use of HAPs to deliver drugs to hypoxic regions in tumours have been identified, including extravascular transport into hypoxic zones^63-65^, activation under mild (physiological) hypoxia in normal tissue^66, 67^ and metabolism by two-electron reductases such as NAD(P)H quinone dehydrogenase 1 (NQO-1) that effect reductive activation but are not inhibited by oxygen^68, 69^.

The present study identifies linker fragmentation via oxidation by CYP3A as a new mechanism of off-target hypoxia-independent/O_2_-dependent activation. This oxidative pathway may have broad implications for the trigger-linker-effector class of HAPs based on our initial survey of HAP structures (Figs. 6,7). Notably, the clinically validated CYP3A-specific inhibitor ritonavir potently inhibits this oxic activation pathway in *in vitro* liver models, while markedly increasing the exposure to **4** and reducing levels of **2** in plasma and normal tissues in mice treated with the HAP (Fig. 8). Modification of the linker to mask sites of CYP-mediated oxidation warrants investigation, although substitution of ether-linked HAPs of combretastatin A-4 with gem-dimethyl linkers is both technically challenging^22^ and been shown to compromise inhibition of reductive activation by O_2_^16^.

The greater suppression of **2** by ritonavir in plasma and normal tissues than in HCT116 tumours, after dosing with HAP **4**, suggests significant intra-tumour activation of the prodrug. However, average levels of **2** in tissues are unlikely to report exposure in critically important sub-compartments such as hypoxic zones in tumours or stem cell niches in normal tissue. For that reason, we turned to pharmacodynamic endpoints to assess therapeutic utility of the co-drug combination of **4** with ritonavir. Notably, doses of **2** providing equivalent tumour clonogen radiosensitisation to **4** + ritonavir (Fig. 9a,b) caused marked bodyweight loss which was not observed with the HAP (Fig. 9c,d). The superior tumour selectivity of the HAP with ritonavir was confirmed by the lack of enhancement of radiation damage to stem cells in the gastrointestinal tract (Fig. 10). These findings argue for the potential of HAP **4** with ritonavir as a co-drug combination for selective tumour radiosensitisation in patients. The need for targeted delivery of DNA-PK inhibitors is underscored by marked enhancement of radiation toxicities in normal tissue by DNA-PKi in both preclinical^37, 38^ and clinical^39-41^ studies. In relation to achieving tumour targeting in patients, it is encouraging that conjugation (and presumed detoxification) of the DNA-PKi metabolite **2** is more extensive in human hepatocytes than in the other species tested (Fig. 2h) so protection due to the phenolic OH ‘safety catch’ can be expected to be even more effective in humans than in preclinical models.

The discovery that HAP **4** is activated almost exclusively by CYP3A in human liver microsomes suggests potential application for cancers with high expression of the enzyme, such as T cell lymphomas resistant to doxorubicin and etoposide because of copy number amplification of *CYP3A4*^70^. However, initial clinical development of HAP **4** + ritonavir would preferably be with ultrahypofractionated radiation (stereotactic body radiotherapy/stereotactic ablative radiotherapy protocols) in settings where hypoxia limits outcomes^13^. It will also be important to evaluate use with conventionally fractioned radiation to assess whether hypoxia can be exploited to generate **2** that will diffuse locally into oxic zones to provide additional radiosensitisation, even in settings where hypoxia is not a limiting factor in local control by radiotherapy.

## Methods

### Cell lines

Sources of cell lines, and culture media, are specified in Table S6. All cell lines were passaged for up to 3 months, without antibiotics, from frozen stocks authenticated by short tandem repeat profiling and confirmed negative for mycoplasma (PlasmoTest, InvivoGen).

### Compounds

DNA-PK inhibitors **1, 2**, HAPs **3**-**5**,**7-9**, internal standard **11** and metabolite **6** were synthesised as detailed in the Supporting Information. Purities of the compounds are documented in Table S1 and sources of purchased compounds in Table S7. Stock solutions in DMSO were stored at -20 °C.

### In vitro radiosensitisation under oxia by regrowth inhibition

HAP1 (*PRKDC*^*+/+*^) or HAP1/*PRKDC*^*-/-*^) cells were seeded in 96-well plates at 200 and 700 cells/well respectively for unirradiated cultures, and at 600 and 7000 cells/well for 3 Gy IR. After attachment for 3 h, drugs were added and plates were irradiated or sham-irradiated 1 h later. After 24 h drugs were removed by washing with fresh medium and plates were incubated for a further 3 days. Cells were stained with sulforhodamine B, and absorbance was calculated as a fraction of the no-drug controls.

### In vitro radiosensitisation under oxia and hypoxia by clonogenic assay

10^5^ cells/well were plated in 96-well plates under oxia (20% O_2_/5% CO_2_), or hypoxia (0% O_2_/5% CO_2_/5% H_2_ in a Bactron anaerobic chamber; Sheldon Manufacturing). 3 h after drug addition, plates were sealed in metal boxes under oxia or inside the anaerobic chamber and irradiated at a range of radiation doses (Eldorado 78 Cobalt-60 teletherapy unit, 0.07-0.32 Gy/min) using a lead wedge to achieve a dose range across each plate^71^. Cultures were then incubated for 18 h under oxia, drugs were removed, cells were trypsinised, enumerated with a particle counter (Beckman Coulter model Z2) and re-plated in 6-well plates. After 10-14 days (Table S6) colonies were stained with methylene blue (2 g/L in 50% EtOH). Colonies with ≥50 cells were counted to determine plating efficiency (PE) and surviving fraction (PE treated/PE untreated).

### Cellular drug metabolism assays

10^6^ cells/0.5 mL were incubated in 24-well plates under oxic or hypoxic conditions as above. Drugs were added after 2 h, with ritonavir (when used) added 20 min prior to **4**. At various times extracellular medium was collected then the cell monolayer was extracted with ice-cold methanol, pooled with the corresponding extracellular sample, and supernatants were mixed with an equal volume of 0.1% formic acid in water and stored at -20 °C for analysis by LC-MS/MS.

### Drug metabolism in post-mitochondrial (S9) liver fractions

Murine (CD1 male) and pooled human liver S9 preparations (Sekisui XenoTech) were incubated in 96 well plates with NADH/NADPH cofactors (1 mM), prodrugs/drugs (10 μM) and inhibitors in 100 mM potassium phosphate buffer containing 5 mM MgCl_2_ and 1 mM disodium EDTA (pH 7.4) buffer at 1-3 mg protein/mL. Plates were equilibrated by shaking at 300 rpm on a Thermomixer (Eppendorf) at 37 °C under oxia (room air), with S9 plus inhibitors and cofactors preincubated for 5 or 20 min prior to adding the test drugs to initiate reaction. At various times, samples (50 μL) were quenched with 200 µL acetonitrile + 0.1% formic acid (typically containing 1 µM **11**) as deuterated internal standard) on ice, and supernatants were stored at -80 °C for LC-MS/MS analysis.

### Drug metabolism in hepatocytes

Human (Lot 2310092), dog (Lot 2010202), rat (Lot 2310093) and mouse (Lot 2310051) cryopreserved hepatocytes were sourced from XenoTech (USA) and suspended in protein-free Krebs-Henseleit buffer (KHB; pH 7.4) at 0.5 x 10^6^ or 2 x 10^6^ viable (Trypan blue excluding) cells/mL. Cell suspensions were equilibrated for 30 min, with or without a pan-CYP inhibitor (pan-CYPi) cocktail (1 mM 1-aminobenzotriazole + 15 μM tienilic acid) on a plate shaker (900 rpm) in a 37 °C humidified incubator (7.5% CO_2_, 19.4% O_2_). After addition of test compounds, or quality control compounds (Table S7), samples were withdrawn at up to 4 h, quenched with acetonitrile (containing 0.15 μg/mL metolazone as internal standard) on ice for 15 min and supernatants were analysed by LC-MS/MS.

### Reaction phenotyping in human liver microsomes (HLM)

HLM (XenoTech, lot 1910096) at 1 mg protein/mL in pH 7.4 0.1 M phosphate buffer containing 3.3 mM MgCl_2_ were incubated at 37 °C with 1.3 mM NADPH for 15 min prior to addition of **2** or **4** (0.5 µM), with and without isoform-specific inhibitors based on published conditions^54^. At various timepoints samples were quenched with ice-cold acetonitrile containing metolazone internal standard and assayed by LC-MS/MS. Lack of interference by the inhibitors with analyte quantitation was confirmed.

### Mouse studies

Animal studies were approved under University of Auckland Animal Ethics Committee Approval 2256. Outbred CD1 Swiss and immunodeficient NIH-III (NIH-Lyst^bg-J^ Foxn1^nu^-Btk^xid^) mice were supplied by the Vernon Jansen Unit, University of Auckland. Mice were monitored every 2-3 days after s.c. inoculation of tumour cells and twice daily after drug or radiation treatment. Compound **2** was formulated for dosing in 5% DMSO/20% PEG400/15% cyclodextrin/60% water and **4** was formulated in 0.5% hydroxypropyl methylcellulose + tween80 in water. Ritonavir stock solution (12.5 mg/mL in 50% EtOH/water) was stored at -20 °C and diluted to 5 mg/mL in water immediately before dosing by oral gavage. Animals held for > 24 h after whole body radiation received wet mash to minimise loss of feeding secondary to oral mucositis. No animals in this study required euthanasia as a result of humane endpoints being reached (body weight loss > 20% or >10% within 24 h, measured twice daily, or clinical signs of toxicity or tumours with largest diameter > 20 mm).

### Ex vivo clonogenic assay

5 x 10^6^ HCT116 or UT-SCC-74B cells were injected subcutaneously on the right flank of female NIH-III mice. When tumours reached approximately 500 mm^3^ (range 350-700 mm^3^) mice were randomised to treatment groups. Unanaesthetised, freely moving mice received whole-body radiation (Eldorado 78 Cobalt-60 unit, 0.6-0.85 Gy/min) following drug treatment as detailed in the figure legends. 18 h after irradiation, tumours were enzymatically digested to single cells, counted and plated in p60 dishes for determination of surviving clonogens as previously described^37^.

### Pharmacokinetic studies

Male CD1 Swiss mice, or female NIH-III mice bearing HCT116 tumours, were treated with various doses of **2** (i.p., 10 mL/kg), or with **4** (p.o., 10 mL/kg). When used, ritonavir (p.o., 5 mL/kg) was dosed 60 min before **4**. Blood was collected by cardiac puncture in EDTA tubes for preparation of plasma. Tissue samples were diluted 1:3 w/v with 4.5 mM formate buffer (pH 4.5) and homogenised using a TissueLyser (Qiagen) at 30 Hz for 3 min with stainless steel beads. Plasma and tissue homogenates were deproteinised with 4 vol of ice-cold acetonitrile, centrifuged (13,000 *g* for 10 min at 4 °C) and supernatants were mixed with an equal volume of 0.1% formic acid in water for LC-MS/MS analysis. Non-compartmental pharmacokinetic analysis was performed using PKSolver^72^.

### LC-MS/MS analyses

Cell culture, S9, plasma and tissue samples were analysed with an Agilent model 6460 LC-MS/MS using electrospray ionisation (ESI) in positive mode with external standard calibration curves or with 1 µM **11** internal standard. Hepatocyte and human liver microsome samples were analysed with a Waters Acquity UPLC interfaced with a Waters Xevo G2-S QToF in both positive and negative mode ESI using external standard calibration for **2** and **4**, and relative quantitation of metabolites using the metabolite-to-internal standard peak area ratio. Details of the bioanalytical methods are provided in Supplementary Information.

### EdU labelling of S-phase cells in the gastrointestinal tract

Female CD-1 mice were dosed i.p. with EdU (Abcam, 50 mg/kg) 2 h prior to tissue collection. Tissues were processed for microscopy and image analysis of EdU+ve and H33342-stained nuclei as previously^37^ with minor variations as detailed in supplementary methods.

### Statistical analysis

Values are arithmetic means and SEM unless otherwise indicated. One-way ANOVA analysis with Tukey’s multiple comparisons test was performed using Graphpad Prism v10. *: p<0.05, **: p<0.01, ***: p<0.001, ****: p<0.0001.

## Supporting information

NMR spectra

## Acknowledgements

The authors thank Dr Peter Davis and Dr Andrew Macann for helpful discussions, Sree Sreebhavan for assistance with LC-MS/MS method development, Karin Tan for semipreparative HPLC, Sisira Kumara for HPLC analyses, and Gary Cheng, Rebecca Airey, Sophia Gortner-O’Brien, Sarah McManaway and Allyza Nerida for assistance with cell culture experiments. This research was supported by funding from the Health Research Council of New Zealand (19/213, 22/444), Cancer Research Trust New Zealand (1641-RPG), Cancer Society New Zealand (17.13), The Maurice Wilkins Centre (MWC4604), Auckland Medical Research Foundation (1117020), Simplicity Foundation and the Peter Neilson Fund for Pancreatic Cancer Research, the Cancer Society Auckland/Northland (Senior Fellowships to M.P.H., S.M.F.J.), and DDRx Pharmaceuticals Ltd. The Centre for Drug Candidate Optimisation (L.Z. and D.M.S.) is partially supported by the Monash University Technology Research Platform network and Therapeutic Innovation Australia (TIA) through the Australian Government National Collaborative Research Infrastructure Strategy (NCRIS) program.

## Author contributions

M.P. H. and W.R.W. designed the research. B.D.D., L.P.L. and M.P.H. synthesised the compounds. C.R.H., W.W.W., J.K.J., J.M.R., L.Z and D.M.S. carried out the experiments and analysed the data. W.R.W., L.P.L., C.R.H., S.M.F.J., D.M.S, and M.P.H. wrote the manuscript. All authors have read and approved the manuscript.

## Supplementary Information

Supplementary methods, Tables and Figures are available in a single PDF file, and NMR spectra in a second PDF.

## References

1 Dhani, N., Fyles, A., Hedley, D. & Milosevic, M. The clinical significance of hypoxia in human cancers. Semin. Nucl. Med. 45, 110–121 (2015).

2 Suvac, A., Ashton, J. & Bristow, R. G. Tumour hypoxia in driving genomic instability and tumour evolution. Nature Rev. Cancer 25, 167–188 (2025).

3 Yotnda, P., Wu, D. & Swanson, A. M. Hypoxic tumours and their effect on immune cells and cancer therapy. Methods Mol. Biol. 651, 1–29 (2010).

4 Labiano, S., Palazon, A. & Melero, I. Immune response regulation in the tumor microenvironment by hypoxia. Semin. Oncol. 42, 378–386 (2015).

5 Chouaib, S., Noman, M. Z., Kosmatopoulos, K. & Curran, M. A. Hypoxic stress: obstacles and opportunities for innovative immunotherapy of cancer. Oncogene 36, 439–445 (2017).

6 Rankin, E. B. & Giaccia, A. J. Hypoxic control of metastasis. Science 352, 175–180 (2016).

7 Singleton, D. C., Macann, A. & Wilson, W. R. Therapeutic targeting of the hypoxic tumour microenvironment. Nature Rev. Clin. Oncol. (2021).

8 Losman, J.-A., Koivunen, P. & Kaelin, W. G. 2-Oxoglutarate-dependent dioxygenases in cancer. Nature Rev. Cancer 20, 710–726 (2020).

9 Radford, I. R. The level of induced DNA double-strand breakage correlates with cell killing after X-irradiation. Int. J. Radiat. Biol. Relat. Stud. Phys. Chem. Med. 48, 45–54 (1985).

10 Grimes, D. R. & Partridge, M. A mechanistic investigation of the oxygen fixation hypothesis and oxygen enhancement ratio. Biomed. Phys. Eng. Express 1, 045209 (2015).

11 Wardman, P. Time as a variable in radiation biology: The oxygen effect. Radiat. Res. 185, 1–3 (2016).

12 Timmerman, R. D., Herman, J. & Cho, L. C. Emergence of stereotactic body radiation therapy and its impact on current and future clinical practice. J. Clin. Oncol. 32, 2847–2854 (2014).

13 Carlson, D. J., Keall, P. J., Loo, B. W., Jr., Chen, Z. J. & Brown, J. M. Hypofractionation results in reduced tumor cell kill compared to conventional fractionation for tumors with regions of hypoxia. Int. J. Radiat. Onc. Biol. Phys. 79, 1188–1195 (2011).

14 Wilson, W. R. & Hay, M. P. Targeting hypoxia in cancer therapy. Nature Rev. Cancer 11, 393–410 (2011).

15 Hay, M. P., Anderson, R. F., Ferry, D. M., Wilson, W. R. & Denny, W. A. Synthesis and evaluation of nitroheterocyclic carbamate prodrugs for use with nitroreductase-mediated gene-directed enzyme prodrug therapy. J. Med. Chem. 46, 5533–5545 (2003).

16 Thomson, P. et al. Synthesis and biological properties of bioreductively targeted nitrothienyl prodrugs of combretastatin A-4. Mol. Cancer Ther. 5, 2886–2894 (2006).

17 Winn, B. A. et al. Bioreductively activatable prodrug conjugates of phenstatin designed to target tumor hypoxia. Bioorg. Med. Chem. Lett. 27, 636–641 (2017).

18 Jin, C., Zhang, Q. & Lu, W. Synthesis and biological evaluation of hypoxia-activated prodrugs of SN-38. Eur. J. Med. Chem. 132, 135–141 (2017).

19 O’Connor, L. J. et al. Design, synthesis and evaluation of molecularly targeted hypoxia-activated prodrugs. Nat. Protoc. 11, 781–794 (2016).

20 Wong, W. W. et al. Hypoxia-selective radiosensitisation by SN38023, a bioreductive prodrug of DNA-dependent protein kinase inhibitor IC87361. Biochem. Pharmacol. 169, 113641 (2019).

21 Liew, L. P. et al. Hypoxia-activated prodrugs of PERK inhibitors. Chem. Asian J. 14, 1238–1248 (2019).

22 Winn, B. A. et al. Bioreductively activatable prodrug conjugates of combretastatin A-1 and combretastatin A-4 as anticancer agents targeted toward tumor-associated hypoxia. J. Nat. Prod. 83, 937–954 (2020).

23 Yan, V. C. et al. Bioreducible phosphonoamidate pro-drug inhibitor of enolase: proof of concept study. ACS Med. Chem. Lett. 11, 1484–1489 (2020).

24 Kim, J. H. et al. A small molecule strategy for targeting cancer stem cells in hypoxic microenvironments and preventing tumorigenesis. J. Am. Chem. Soc. 143, 14115–14124 (2021).

25 Skwarska, A. et al. Development and pre-clinical testing of a novel hypoxia-activated KDAC inhibitor. Cell Chem. Biol. 28, 1258-1270.e1213 (2021).

26 Wang, S. et al. Discovery of a highly efficient nitroaryl group for detection of nitroreductase and imaging of hypoxic tumor cells. Org. Biomol. Chem. 19, 3469–3478 (2021).

27 Tosun, Ç. et al. Antibody-based imaging of bioreductive prodrug release in hypoxia. JACS Au 3, 3237–3246 (2023).

28 Shi, Z. et al. Targeting tumor-associated hypoxia with bioreductively activatable prodrug conjugates derived from dihydronaphthalene, benzosuberene, and indole-based inhibitors of tubulin polymerization. RSC Med. Chem. 16, 5472–5487 (2025).

29 Wong, W. W. et al. Hypoxia-activated prodrugs of phenolic olaparib analogues for tumour-selective chemosensitisation. RSC Med. Chem. 14, 1309–1330 (2023).

30 Mahaney, B. L., Meek, K. & Lees-Miller, S. P. Repair of ionizing radiation-induced DNA double-strand breaks by non-homologous end-joining. Biochem. J. 417, 639–650 (2009).

31 Ghosh, D. & Raghavan, S. C. Nonhomologous end joining: new accessory factors fine tune the machinery. Trends Genet. 37, 582–599 (2021).

32 Chaplin, A. K. et al. Dimers of DNA-PK create a stage for DNA double-strand break repair. Nature Struct. Mol. Biol. 28, 13–19 (2021).

33 Vogt, A., He, Y. & Lees-Miller, S. P. How to fix DNA breaks: new insights into the mechanism of non-homologous end joining. Biochem. Soc. Trans. 51, 1789–1800 (2023).

34 Fulop, G. M. & Phillips, R. A. The scid mutation in mice causes a general defect in DNA repair. Nature 347, 479–482 (1990).

35 Biedermann, K. A., Sun, J. R., Giaccia, A. J., Tosto, L. M. & Brown, J. M. Scid mutation in mice confers hypersensitivity to ionizing radiation and a deficiency in DNA double-strand break repair. Proc. Natl Acad. Sci. U.S.A. 88, 1394–1397 (1991).

36 Lee, T. W. et al. Radiosensitization of head and neck squamous cell carcinoma lines by DNA-PK inhibitors is more effective than PARP-1 inhibition and is enhanced by SLFN11 and hypoxia. Int. J. Radiat. Biol. 95, 1597–1612 (2019).

37 Hong, C. R. et al. Radiosensitisation of SCCVII tumours and normal tissues in mice by the DNA-dependent protein kinase inhibitor AZD7648. Radiother. Oncol. 166, 162–170 (2022).

38 Baker, J. H. E. et al. Radiation and chemo-sensitizing effects of DNA-PK inhibitors are proportional in tumors and normal tissues. Mol. Cancer Ther. 23, 1230–1240 (2024).

39 van der Burg, M. et al. A DNA-PKcs mutation in a radiosensitive T-B-SCID patient inhibits Artemis activation and nonhomologous end-joining. J. Clin. Invest. 119, 91–98 (2009).

40 Samuels, M. et al. A phase 1 study of the DNA-PK inhibitor peposertib in combination with radiation therapy with or without cisplatin in patients with advanced head and neck tumors. Int. J. Radiat. Oncol. Biol. Phys. 118, 743–756 (2024).

41 Romesser, P. B. et al. A phase Ib study of the DNA-PK Inhibitor peposertib combined with neoadjuvant chemoradiation in patients with locally advanced rectal cancer. Clin. Cancer Res. 30, 695–702 (2024).

42 Hong, C. R. et al. Identification of 6-anilino imidazo [4,5-c]pyridin-2-ones as selective DNA-dependent protein kinase inhibitors and their application as radiosensitizers. J. Med. Chem. 67, 12366–12385 (2024).

43 Lamb, Y. N. Nirmatrelvir plus ritonavir: First approval. Drugs 82, 585–591 (2022).

44 Gerhart, J. et al. A comprehensive review of the clinical pharmacokinetics, pharmacodynamics, and drug interactions of nirmatrelvir/ritonavir. Clin. Pharmacokinet. 63, 27–42 (2024).

45 Luukkanen, L. et al. Kinetic characterization of the 1A subfamily of recombinant human UDP-glucuronosyltransferases. Drug Metab. Dispos. 33, 1017–1026 (2005).

46 Lu, L. et al. Drug-metabolizing activity, protein and gene expression of UDP-glucuronosyltransferases are significantly altered in hepatocellular carcinoma patients. PLOS ONE 10, e0127524 (2015).

47 Margaillan, G. et al. Quantitative profiling of human renal UDP-glucuronosyltransferases and glucuronidation activity: a comparison of normal and tumoral kidney tissues. Drug Metab. Dispos. 43, 611–619 (2015).

48 Hu, D. G. et al. The expression profiles and deregulation of UDP-glycosyltransferase (UGT) genes in human cancers and their association with clinical outcomes. Cancers 13 (2021).

49 Wardman, P. Electron transfer and oxidative stress as key factors in the design of drugs selectively active in hypoxia. Curr. Med. Chem. 8, 739–761 (2001).

50 Su, J. et al. Zinc finger nuclease knockout of NADPH: cytochrome P450 oxidoreductase (POR) in human tumour cell lines demonstrates that hypoxia-activated prodrugs differ in POR dependence. J. Biol. Chem. 288, 37138–37153 (2013).

51 Hunter, F. W. et al. Identification of P450 oxidoreductase as a major determinant of sensitivity to hypoxia-activated prodrugs. Cancer Res. 75, 4211–4223 (2015).

52 Lee, T. W. et al. Clonal dynamics limits detection of selection in tumour xenograft CRISPR/Cas9 screens. Cancer Gene Ther. 30, 1610–1623 (2023).

53 Lee, T. W. et al. Clinical relevance and therapeutic predictive ability of hypoxia biomarkers in head and neck cancer tumour models. Mol. Oncol. 18, 1885–1903 (2024).

54 Nirogi, R. et al. Chemical inhibitors of CYP450 enzymes in liver microsomes: combining selectivity and unbound fractions to guide selection of appropriate concentration in phenotyping assays. Xenobiotica 45, 95–106 (2015).

55 Loos, N. H. C., Beijnen, J. H. & Schinkel, A. H. The Mechanism-Based Inactivation of CYP3A4 by Ritonavir: What Mechanism? Int. J. Mol. Sci. 23, 9866 (2022).

56 von Moltke, L. L., Durol, A. L. B., Duan, S. X. & Greenblatt, D. J. Potent mechanism-based inhibition of human CYP3A in vitro by amprenavir and ritonavir: comparison with ketoconazole. Eur. J. Clin. Pharmacol. 56, 259–261 (2000).

57 Keubler, A., Weiss, J., Haefeli, W. E., Mikus, G. & Burhenne, J. Drug interaction of efavirenz and midazolam: Efavirenz activates the CYP3A-mediated midazolam 1’-hydroxylation in vitro. Drug Metab. Disp. 40, 1178–1182 (2012).

58 Garcia, D. A. et al. Modeling the acute mucosal toxicity of fractionated radiotherapy combined with the ATM inhibitor WSD0628. Mol. Cancer Ther. 24, 299–309 (2025).

59 Langius, J. A. E. et al. Critical weight loss is a major prognostic indicator for disease-specific survival in patients with head and neck cancer receiving radiotherapy. Brit. J. Cancer 109, 1093–1099 (2013).

60 Ang, K. K. et al. Randomized phase III trial of concurrent accelerated radiation plus cisplatin with or without cetuximab for stage III to IV head and neck carcinoma: RTOG 0522. J. Clin. Oncol. 32, 2940–2950 (2014).

61 Adams, G. E., Dische, S., Fowler, J. F. & Thomlinson, R. H. Hypoxic cell sensitisers in radiotherapy. Lancet 1, 186–188 (1976).

62 Brown, J. M. Clinical perspectives for the use of new hypoxic cell sensitizers. Int. J. Radiat. Oncol. Biol. Phys. 8, 1491–1497 (1982).

63 Hicks, K. O. et al. Use of three-dimensional tissue cultures to model extravascular transport and predict in vivo activity of hypoxia-targeted anticancer drugs. J. Natl Cancer Inst. 98, 1118–1128 (2006).

64 Minchinton, A. I. & Tannock, I. F. Drug penetration in solid tumours. Nature Rev. Cancer 6, 583–592 (2006).

65 Dewhirst, M. W. & Secomb, T. W. Transport of drugs from blood vessels to tumour tissue. Nature Rev. Cancer 17, 738–750 (2017).

66 Wilson, W. R. et al. Bystander effects of bioreductive drugs: potential for exploiting pathological tumor hypoxia with dinitrobenzamide mustards. Radiat. Res. 167, 625–636 (2007).

67 Gu, Y. et al. Reductive metabolism influences the toxicity and pharmacokinetics of the hypoxia-targeted benzotriazine di-oxide anticancer agent SN30000 in mice. Front. Pharmacol. 8, 531 (2017).

68 Bailey, S. M. et al. Involvement of NADPH: cytochrome P450 reductase in the activation of indoloquinone EO9 to free radical and DNA damaging species. Biochem. Pharmacol. 62, 461–468 (2001).

69 Yan, C., Kepa, J. K., Siegel, D., Stratford, I. J. & Ross, D. Dissecting the role of multiple reductases in bioactivation and cytotoxicity of the antitumor agent 2,5-diaziridinyl-3-(hydroxymethyl)-6-methyl-1,4-benzoquinone (RH1). Mol. Pharmacol. 74, 1657–1665 (2008).

70 Rodríguez-Antona, C. et al. Expression of CYP3A4 as a predictor of response to chemotherapy in peripheral T-cell lymphomas. Blood 110, 3345–3351 (2007).

71 Cross, P. et al. Proliferative assays for the assessment of radiosensitivity of tumor cell lines using 96-well microcultures. Radiat. Oncol. Invest. 1, 261–269 (1994).

72 Zhang, Y., Huo, M., Zhou, J. & Xie, S. PKSolver: An add-in program for pharmacokinetic and pharmacodynamic data analysis in Microsoft Excel. Comp. Meth. Prog. Biomed. 99, 306–314 (2010).

