## Supplemental Information for "Systemic CYP3A inhibition by ritonavir enables selective targeting of hypoxic tumour cells by prodrugs of DNA-PK inhibitors"

### Table of Contents


| | | Pg |
|---|---|---|
| Supplementary Methods | General synthetic methods and materials | 2 |
| | Synthesis and characterisation of compounds | 2 |
| | Bioanalytical methods | 10 |
| | Microscopy and image analysis | 12 |
| Supplementary Tables | Table S1. DNA-PKi and HAPs: structures, kinase inhibition, purity and radiosensitisation of HAP1 cells by regrowth assay | 13 |
| | Table S2. Radiosensitisation by DNA-PK inhibitors: clonogenic assay data | 14 |
| | Table S3. Radiosensitisation by HAPs of DNA-PK inhibitors: clonogenic assay data | 15 |
| | Table S4. Intrinsic clearance of HAP **4,** yields of DNA-PKi **2**, and effect of pan-CYPi in oxic hepatocytes | 16 |
| | Table S5. Metabolites of HAP **4** in oxic hepatocytes | 17 |
| | Table S6. Cell lines and culture media | 18 |
| | Table S7. Sources of reference substrates, inhibitors and quality control compounds | 19 |
| Supplementary Figures | Fig. S1. Formation of an *O*-glucuronide from DNA-PKi **2** in BxPC-3 cells | 20 |
| | Fig. S2. Collision-induced dissociation of metabolites of DNA-PKi **2** | 21 |
| | Fig. S3. Inferred metabolic pathway of metabolism of DNA-PKi **2** | 23 |
| | Fig. S4. Metabolism of ether-linked DNA-PKi HAPs in oxic mouse liver S9 | 24 |
| | Fig. S5: Metabolite profile of HAP **4** with and without pan-CYPi, in oxic mouse, rat, dog and human hepatocytes | 25 |
| | Fig. S6. Provisional identification of a minus 2 Da metabolite of **4** | 26 |
| | Fig. S7**.** Metabolism of **4** in oxic human liver microsomes is not sensitive to other isoform-specific CYP inhibitors | 27 |

| | | |
|---|---|---|
| | Fig. S8. Tolerability of IP **2** and oral **4** + Rito in CD-1 mice | 28 |
| | Fig. S9. Images of EdU+ cells in mouse ileum sections (individual mice) | 29 |
| | Fig, S10. Images of EdU+ cells in mouse tongue sections (individual mice) | 32 |
| Supplementary references | | 35 |


### Supplementary Methods

#### *General synthetic methods and materials*

Analyses were carried out in the Microchemical Laboratory, University of Otago, Dunedin, NZ. Melting points were determined on an Electrothermal 2300 Melting Point Apparatus. NMR spectra were obtained on a Bruker Avance 400 spectrometer at 400 MHz for $^1$H and 100 MHz for $^{13}$C spectra. Spectra were obtained in $CDCl_3$, unless otherwise specified, and were referenced to $Me_4Si$. Spectra in $(CD_3)_2SO$ are referenced to the residual solvent peak. Chemical shifts and coupling constants were recorded in units of ppm and Hz, respectively. Assignments were determined using COSY, HSQC, and HMBC two-dimensional experiments. Low resolution mass spectra were gathered by direct injection of methanolic solutions into a Agilent Technologies 6120 Quadrupole LCMS using an atmospheric pressure chemical ionization (APCI) mode with a corona voltage of 50 V and a source temperature of 400 °C. High resolution mass spectra (HRMS) were measured on an Agilent Technologies 6530 Accurate-Mass Quadrupole Time of Flight LCMS interfaced with an Agilent Jet Stream Electrospray Ionization (ESI) source allowing positive or negative ions detection. Solutions in organic solvents were dried with anhydrous $MgSO_4$. Solvents were evaporated under reduced pressure on a rotary evaporator. Thin-layer chromatography was carried out on aluminium-backed silica gel plates (Merck 60 $F_{254}$) with visualization of components by UV light (254 nm) or exposure to $I_2$. Column chromatography was carried out on silica gel (Merck 230-400 mesh). All final products were analysed by RP-HPLC (Altima C18 5 µm column, 150 mm × 3.2 mm; Alltech Associated, Inc., Deerfield, IL) using an Agilent HP1260 LC equipped with a diode-array detector. Mobile phases were gradients of 80% MeCN/20% $H_2O$ (v/v) in 45 mM ammonium formate at pH 3.5 and 0.5 mL/min. Purity was determined by monitoring at 330 (± 100 nm) and was >95%.

#### *Synthesis and characterisation of compounds*

**6-((4-Methoxy-2-methylphenyl)amino)-3-methyl-1-(tetrahydro-2*H*-pyran-4-yl)-1,3-dihydro-2*H*-imidazo[4,5-*c*]pyridin-2-one (1).** Compound **1** was prepared as described previously[1]. HPLC purity 99.6%.

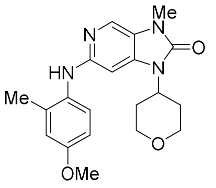


**1**

**6-((4-Hydroxy-2-methylphenyl)amino)-3-methyl-1-(tetrahydro-2*H*-pyran-4-yl)-1,3-dihydro-2*H*-imidazo[4,5-*c*]pyridin-2-one (2).**

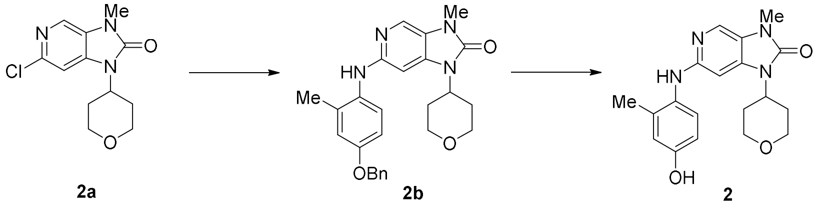


**2a** **2b** **2**

**6-((4-(Benzyloxy)-2-methylphenyl)amino)-3-methyl-1-(tetrahydro-2*H*-pyran-4-yl)-1,3-dihydro-2*H*-imidazo[4,5-c]pyridin-2-one (2b).** A degassed mixture of 6-chloro-3-methyl-1-(tetrahydro-2*H*-pyran-4-yl)-1,3-dihydro-2*H*-imidazo[4,5-*c*]pyridin-2-one[1] **2a** (415 mg, 1.55 mmol), 4-(benzyloxy)-2-methylaniline (397 mg, 1.86 mmol), $Pd_2dba_3$ (71 mg, 78 µmol), XPhos (148 mg, 310 µmol) and $Cs_2CO_3$ (1.10 g, 3.41 mmol) in MeCN (8 mL) was stirred in a sealed tube at 120 °C for 16 h. The mixture was cooled and the mixture diluted with EtOAc (30 mL) and filtered through diatomaceous earth and the filtrate was evaporated. The residue was partitioned between EtOAc (50 ml) and water (50 mL). The organic fraction was washed with water (30 mL), washed with brine (30 mL), dried, filtered and the solvent evaporated. The residue was purified by chromatography, eluting with EtOAc, to give imidazopyridinone **2b** (538 mg, 78%) as white needles: mp (EtOAc/pet ether) 179–181 °C; $^1$H NMR δ 7.78 (s, 1 H, H-4), 7.47 (br d, *J* = 7.0 Hz, 2 H, H-2‴, H-6‴), 7.40 (br dd, *J* = 7.5, 7.1 Hz, 2 H, H-3‴, H-5‴), 7.34 (br t, *J* = 7.2 Hz, 1 H, H-4‴), 7.26 (d, *J* = 8.6 Hz, 1 H, H-6″), 6.93 (d, *J* = 2.9 Hz, 1 H, H-3″), 6.84 (dd, *J* = 8.6, 2.9 Hz, 1 H, H-5″), 6.26 (d, *J* = 0.5 Hz, 1 H, H-7), 5.94 (s, 1 H, 6-NH), 5.08 (s, 2 H $CH_2O$), 4.41 (tt, *J* = 12.4, 4.2 Hz, 1 H, H-4′), 4.07 (dd, *J* = 11.6, 4.4 Hz, 2 H, H-2′, H-6′), 3.50 (dt, *J* = 11.9, 1.6 Hz, 2 H, H-2′, H-6′), 3.33 (s, 3 H, 3-$CH_3$), 2.20–2.35 (m, 5 H, 2″-$CH_3$, H-3′, H-5′), 1.69 (dd, *J* = 12.4, 2.5 Hz, 2 H, H-3′, H-5′); MS *m/z* 445.2 ($MH^+$, 100%); HRMS calcd for $C_{26}H_{29}N_4O_3$ ($MH^+$) *m/z* 445.2234, found 445.2250 (-3.6 ppm). HPLC purity 98.9%.

**6-((4-Hydroxy-2-methylphenyl)amino)-3-methyl-1-(tetrahydro-2*H*-pyran-4-yl)-1,3-dihydro-2*H*-imidazo[4,5-c]pyridin-2-one (2).** A mixture of benzyl ether **2b** (172 mg, 0.40 mmol) and Pd/C (20 mg) in a mixture of EtOAc (25 mL) and EtOH (25 mL) was stirred under $H_2$ (50 psi) at 20 °C for 6 h. The mixture was filtered through diatomaceous earth and the filtrate was evaporated. The residue was purified by chromatography, eluting with a gradient (50–100%) of EtOAc/pet. ether, to give imidazopyridinone **2** (44 mg, 32%) as a cream powder: mp (EtOAc/pet ether) 258–261 °C; $^1$H NMR δ 9.00 (s, 1 H, 4″-OH), 7.76 (s, 1 H, H-4), 7.56 (s, 1 H, 6-NH), 7.22 (d, *J* = 8.5 Hz, 1 H, H-6″), 6.16 (d, *J* = 2.7 Hz, 1 H, H-3″), 6.54 (dd, *J* = 8.5, 2.8 Hz, 1 H, H-5″), 6.51 (s, 1 H, H-7), 4.33 (tt, *J* = 12.3, 4.2 Hz, 1 H, H-4′), 3.96 (br dd, *J* = 11.4, 4.2 Hz, 2 H, H-2′, H-6′), 3.44 (br t, *J* = 11.3 Hz, 2 H, H-2′, H-6′), 3.25 (s, 3 H, 3-$CH_3$), 2.25 (dq, *J* = 12.4, 4.5 Hz, 2 H, H-3′, H-5′), 2.20 (s, 3 H, 2″-$CH_3$), 1.63 (dd, *J* = 12.4, 2.9 Hz, 2 H, H-3′, H-5′); $^{13}$C NMR δ 153.4, 153.3, 152.9, 136.3, 133.0, 131.4, 125.4, 125.3, 120.8, 116.9, 112.7, 88.1, 66.4 (2), 49.3, 29.4 (2), 27.0, 18.2; MS *m/z* 355.2 ($MH^+$, 100%); HRMS calcd for $C_{19}H_{23}N_4O_3$ ($MH^+$) *m/z* 355.1765, found 355.1782 (-4.9 ppm). HPLC purity 99.1%.

**3-Methyl-6-((2-methyl-4-((1-methyl-2-nitro-1*H*-imidazol-5-yl)methoxy)phenyl)amino)-1-(tetrahydro-2*H*-pyran-4-yl)-1,3-dihydro-2*H*-imidazo[4,5-*c*]pyridin-2-one (3).**

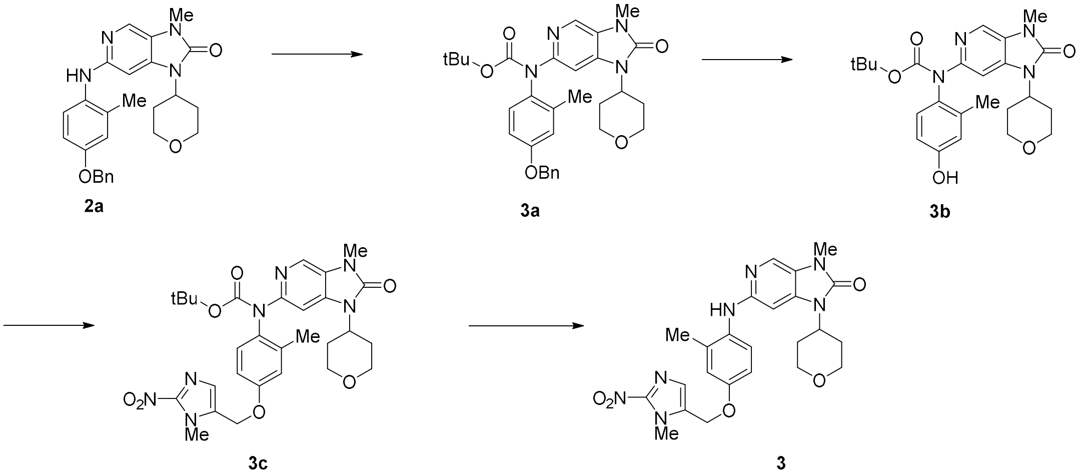


***tert*-Butyl (4-(benzyloxy)-2-methylphenyl)(3-methyl-2-oxo-1-(tetrahydro-2*H*-pyran-4-yl)-2,3-dihydro-1*H*-imidazo[4,5-c]pyridin-6-yl)carbamate (3a).** A mixture of aniline **2a** (416 mg, 0.89 mmol), $iPr_2NEt$ (0.23 mL, 1.34 mmol), DMAP (11 mg, 0.09 mmol) and $BOC_2O$ (388 mg, 1.78 mmol) in dry THF (25 ml) was stirred at 70 °C for 16 h. The mixture was cooled to 20 °C, diluted with EtOAc (100 mL) and washed with water (2 × 25 mL), brine (30 mL), dried, and the solvent evaporated. The residue was purified by chromatography, eluting with a gradient (75–100%) of EtOAc/pet. ether, to give carbamate **3a** (511 mg, 100%) as a white foam: $^1$H NMR δ 7.93 (s, 1 H, H-4′), 7.50 (s, 1 H, H-7′), 7.43 (br d, *J* = 8.4 Hz, 2 H, H-2′′′′, H-6′′′′), 7.38 (ddd, *J* = 8.2, 7.1, 1.5 Hz, 2 H, H-3′′′′, H-5′′′′), 7.32 (tt, *J* = 7.1, 1.5 Hz, 1 H, H-4′′′′), 7.09 (d, *J* = 8.6 Hz, 1 H, H-6′′′), 6.88 (d, *J* = 2.9 Hz, 1 H, H-3′′′), 6.80 (dd, *J* = 8.6, 2.9 Hz, 1 H, H-5′′′), 5.04 (s, 2 H, $CH_2O$), 4.54 (tt, *J* = 12.4, 4.2 Hz, 1 H, 1′-CH), 4.16 (br dd, *J* = 11.6, 4.3 Hz, 2 H, H-2′′, H-6′′), 3.57 (br dt, *J* = 11.9, 1.5 Hz, 2 H, H-2′′, H-6′′), 3.38 (s, 3 H, 3′-$CH_3$), 2.47 (dq, *J* =12.6, 4.6 Hz, 2 H, H-3′′, H-5′′), 2.23 (s, 3 H, 2′′′-$CH_3$), 1.78 (br dd, *J* = 12.4, 2.4 Hz, 2 H, H-3′, H-5′), 1.44 (s, 9 H, $CO_2$tBu); MS *m/z* 545.2 ($MH^+$, 100%); HRMS calcd for $C_{31}H_{37}N_4O_5$ ($MH^+$) *m/z* 545.2758, found 545.2780 (-4.0 ppm).

***tert*-Butyl (4-hydroxy-2-methylphenyl)(3-methyl-2-oxo-1-(tetrahydro-2*H*-pyran-4-yl)-2,3-dihydro-1*H*-imidazo[4,5-c]pyridin-6-yl)carbamate (3b).** A mixture of benzyl ether **3a** (498 mg, 0.91 mmol) and Pd/C (50 mg) in EtOH (50 mL) was stirred under $H_2$ (50 psi) at 20 °C for 16 h. The mixture was filtered through diatomaceous earth and the filtrate was evaporated. The residue was purified by chromatography, eluting with a gradient (50–100%) of EtOAc/pet. ether, to give phenol **3b** (402 mg, 97%) as a white powder: mp 223 °C (dec.); $^1$H NMR δ 7.93 (s, 1 H, H-4′), 7.52 (s, 1 H, H-7′), 6.97 (dd, *J* = 6.4, 2.7 Hz, 1 H, H-5′′′), 6.53–6.57 (m, 2 H, H-3′′′, H-6′′′), 6.07 (s, 1 H, 4′′′-OH), 4.54 (tt, *J* = 12.4, 4.3 Hz, 1 H, 1′-CH), 4.16 (br dd, *J* = 11.5, 4.4 Hz, 2 H, H-2′′, H-6′′), 3.58 (br dt, *J* = 12.0, 1.5 Hz, 2 H, H-2′′, H-6′′), 3.39 (s, 3 H, 3′-$CH_3$), 2.47 (dq, *J* =12.5, 4.6 Hz, 2 H, H-3′′, H-5′′), 2.17 (s, 3 H, 2′′′-$CH_3$), 1.80 (br dd, *J* = 12.4, 2.4 Hz, 2 H, H-3′, H-5′), 1.44 (s, 9 H, $CO_2$tBu); MS *m/z* 455.2 ($MH^+$, 100%); HRMS calcd for $C_{24}H_{31}N_4O_5$ ($MH^+$) *m/z* 455.2302, found 455.2314 (-2.5 ppm).

***tert*-Butyl (3-methyl-2-oxo-1-(tetrahydro-2*H*-pyran-4-yl)-2,3-dihydro-1*H*-imidazo[4,5-c]pyridin-6-yl)(2-methyl-4-((1-methyl-2-nitro-1*H*-imidazol-5-yl)methoxy)phenyl)carbamate (3c).** A mixture of phenol **3b** (395 mg, 0.87 mmol), 5-(chloromethyl)-1-methyl-2-nitro-1*H*-imidazole[2] (168 mg, 0.96 mmol) and $Cs_2CO_3$ (369 mg, 1.13 mmol) in dry DMF (20 mL) was stirred at 60 °C for 3 h. The mixture was cooled and diluted with EtOAc (100 mL). The organic fraction was washed with water (3 × 50 mL), washed with brine (30 mL), dried, filtered and the solvent evaporated. The residue was purified by chromatography, eluting with a gradient (80–100%) of EtOAc/pet. ether, to give carbamate **3c** (185 mg, 36%) as a clear oil: $^1$H NMR δ 7.91 (s, 1 H, H-4′′′), 7.54 (s, 1 H, H-4′′), 7.21 (s, 1 H, H-7′′′), 7.15 (d, *J* = 8.6 Hz, 1 H, H-6′), 6.87 (d, *J* = 2.9 Hz, 1 H, H-3′), 6.80 (dd, *J* = 8.6, 2.9 Hz, 1 H, H-5′), 5.04 (s, 2 H, $CH_2O$), 4.54 (tt, *J* = 12.4, 4.2 Hz, 1 H, 1′′′-CH), 4.18 (br dd, *J* = 11.6, 4.2 Hz, 2 H, H-2′′′′, H-6′′′′), 4.05 (s, 3 H, 1′′′-$CH_3$), 3.58 (br t, *J* = 11.9 Hz, 2 H, H-2′′′′, H-6′′′′), 3.38 (s, 3 H, 3′′′-$CH_3$), 2.48 (dq, *J* =12.6, 4.6 Hz, 2 H, H-3′′′, H-5′′′), 2.17 (s, 3 H, 2′′′-$CH_3$), 1.80 (br dd, *J* = 12.6, 2.6 Hz, 2 H, H-3′′′′, H-5′′′′), 1.43 (s, 9 H, $CO_2$tBu); MS *m/z* 594.2 ($MH^+$, 100%).

**3-Methyl-6-((2-methyl-4-((1-methyl-2-nitro-1*H*-imidazol-5-yl)methoxy)phenyl)amino)-1-(tetrahydro-2*H*-pyran-4-yl)-1,3-dihydro-2*H*-imidazo[4,5-c]pyridin-2-one (3).** TFA (0.5 mL, 6.2 mmol) was added to a stirred solution of carbamate **3c** (185 mg, 0.31 mmol) in DCM (10 mL) and the mixture was stirred at 20 °C for 24 h. The solvent was evaporated and the residue was partitioned between EtOAc (50 ml) and aqueous $NaHCO_3$ (50 mL). The organic fraction was washed with water (30 mL), washed with brine (30 mL), dried, filtered and the solvent evaporated. The residue was purified by chromatography, eluting with a gradient (0–5%) of MeOH/DCM, to give imidazopyridinone **3** (65 mg, 42%) as a clear glass: $^1$H NMR δ 7.78 (s, 1 H, H-4), 7.33 (d, *J* = 8.7 Hz, 1 H, H-6′′), 7.24 (s, 1 H, H-4′′′), 6.90 (d, *J* = 2.9 Hz, 1 H, H-3′′), 6.82 (dd, *J* = 8.7, 2.8 Hz, 1 H, H-5′′), 6.33 (s, 1 H, H-7), 6.14 (br s, 1 H, 6-NH), 5.07 (s, 2 H, $CH_2O$), 4.49 (tt, *J* = 12.4, 4.2 Hz, 1 H, 1-CH), 4.03–4.15 (m, 5 H, H-2′, H-6′, 1′′′-$CH_3$), 3.51 (br t, *J* = 11.9 Hz, 2 H, H-2′, H-6′), 3.40 (s, 3 H, 3-$CH_3$), 2.20–2.32 (m, 5 H, H-3′, H-5′, 2′′′-$CH_3$), 1.72 (br dd, *J* = 12.4, 2.5 Hz, 2 H, H-3′, H-5′); $^{13}$C NMR δ 154.5, 153.9, 153.1, 143.5, 137.3, 134.2, 133.8, 132.9, 129.2, 126.0, 124.8, 122.2, 117.8, 113.2, 88.4, 67.6, 60.1 (2), 50.2, 34.8, 30.2 (2), 27.6, 18.6; MS *m/z* 494.2 ($MH^+$, 100%); HRMS calcd for $C_{24}H_{28}N_7O_5$ ($MH^+$) *m/z* 494.2416, found 494.2151 (-0.9 ppm). HPLC purity 97.1%.

**3-Methyl-6-((2-methyl-4-(1-(1-methyl-2-nitro-1*H*-imidazol-5-yl)ethoxy)phenyl)amino)-1-(tetrahydro-2*H*-pyran-4-yl)-1,3-dihydro-2*H*-imidazo[4,5-c]pyridin-2-one (4).**

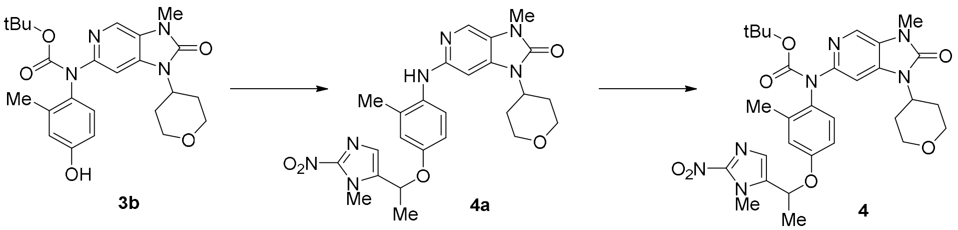


***tert*-Butyl (3-methyl-2-oxo-1-(tetrahydro-2*H*-pyran-4-yl)-2,3-dihydro-1*H*-imidazo[4,5-*c*]pyridin-6-yl)(2-methyl-4-(1-(1-methyl-2-nitro-1*H*-imidazol-5-yl)ethoxy)phenyl)carbamate (237).** A solution of 5-(1-chloroethyl)-1-methyl-2-nitro-1*H*-imidazole[3] (83 mg, 0.44 mmol) in DMF (1 mL) and $Cs_2CO_3$ (170 mg, 0.52 mmol) was added to phenol **3b** (180 mg, 0.40 mmol) in DMF (2 mL) and the mixture was stirred 18 h. The mixture was cooled to 0 °C and diluted with water (5 mL). The aqueous mixture was extracted with EtOAc (3 × 5 mL), the combined organic fractions were washed with water (2 × 10 mL), dried, and solvent was removed *in vacuo*. The resulting residue was purified by chromatography, eluting with EtOAc, to give carbamate **4a** (240 mg, quant.): $^1$H NMR δ 7.91 (s, 1 H, H-4), 7.52 (s, 1 H, H-7), 7.18 (s, 1 H, H-4‴), 7.11 (d, *J* = 8.6 Hz, 1 H, H-6″), 6.82 (d, *J* = 2.8 Hz, 1 H, H-3″), 6.76 (dd, *J* = 8.8, 2.9 Hz, 1 H, H-5″), 5.43 (q, *J* = 6.5 Hz, 1 H, 5‴-CC*H*($CH_3$)), 4.56 (tt, *J* = 12.5, 4.2 Hz, 1 H, H-4′), 4.17 (dd, *J* = 11.6, 4.2 Hz, 2 H, H-2′, H-6′), 4.01 (1‴-$NCH_3$), 3.58 (td, *J* = 12.0, 1.5 Hz, 2 H, H-2′, H-6′), 3.39 (s, 3 H, 3-$NCH_3$), 2.46 (app. qd, *J* = 12.6, 4.6 Hz, H-3′, H-5′), 2.23 (s, 3 H, 2″-$CCH_3$), 1.84–1.74 (m, 5 H, 5‴-CCH(C*H*$_3$), H-3′, H-5′), 1.44 (s, 9 H, C($CH_3$)$_3$).

**3-Methyl-6-((2-methyl-4-(1-(1-methyl-2-nitro-1*H*-imidazol-5-yl)ethoxy)phenyl)amino)-1-(tetrahydro-2*H*-pyran-4-yl)-1,3-dihydro-2*H*-imidazo[4,5-c]pyridin-2-one (4).** TFA (0.51 mL, 6.6 mmol) was added to a solution of carbamate **4a** (200 mg, 0.33 mmol) in DCM (10 mL) and the resulting mixture was stirred for 24 h. The solvent removed *in vacuo* and the residue purified by chromatography, eluting with 100% EtOAc, to give ether **4** (100 mg, 59%) as a yellow solid: mp 229–231 °C; $^1$H NMR [$(CD_3)_2SO$] δ 7.78 (s, 1 H, H-4), 7.68 (s, 1 H, NH), 7.55 (d, *J* = 8.8 Hz, 1 H, H-6″), 7.27 (s, 1 H, H-4‴), 6.90 (d, *J* = 2.8 Hz, 1 H, H-3″), 6.82 (dd, *J* = 8.8, 2.9 Hz, 1 H, H-5″), 6.75 (s, 1 H, H-7), 5.68 (q, *J* = 6.4 Hz, 1 H, 5‴-CCH), 4.37 (tt, *J* = 12.3, 4.2 Hz, 1 H, H-4′), 3.99 (dd, *J* = 11.4, 4.1, 2 H, H-2′, H-6′), 3.95 (s, 3 H, 1‴-$NCH_3$), 3.46 (t, *J* = 11.4 Hz, 2 H, H-2′, H-6′), 3.27 (s, 3 H, 3-$NCH_3$), 2.31–2.18 (m, 2 H, H-3′, H-5′), 2.19 (s, 3 H, 2″-$CCH_3$), 1.70–1.61 (m, 2 H, H-3′, H-5′), 1.64 (d, *J* = 6.4 Hz, 3 H, 5‴-CCH(C*H*$_3$)); $^{13}$C NMR [$(CD_3)_2SO$] δ 153.4, 152.8, 152.1, 146.5, 138.9, 136.7, 134.9, 131.9, 126.7, 125.6, 123.7, 121.8, 118.6, 114.0, 90.0, 67.2, 67.0 (2), 49.7, 34.9, 29.9 (2), 27.5, 19.3, 18.8; HRMS calcd for $C_{25}H_{30}N_7O_5$ ($MH^+$) *m/z* 508.2303, 508.2314 (+2.15 ppm). HPLC purity 99.1%.

**3-Methyl-6-((2-methyl-4-((1-methyl-5-nitro-1*H*-imidazol-2-yl)methoxy)phenyl)amino)-1-(tetrahydro-2*H*-pyran-4-yl)-1,3-dihydro-2*H*-imidazo[4,5-c]pyridin-2-one (5).**

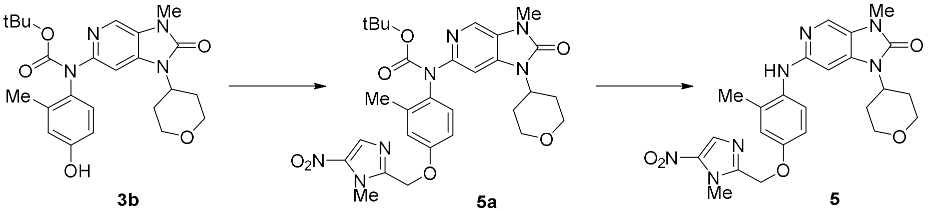


***tert*-Butyl (3-methyl-2-oxo-1-(tetrahydro-2*H*-pyran-4-yl)-2,3-dihydro-1*H*-imidazo[4,5-*c*]pyridin-6-yl)(2-methyl-4-((1-methyl-5-nitro-1*H*-imidazol-2-yl)methoxy)phenyl)carbamate (5a).** 5-(Chloromethyl)-1-methyl-2-nitro-1*H*-imidazole[2] (102 mg, 0.57 mmol) and $Cs_2CO_3$ (190 mg, 0.57 mmol) were added to a solution of phenol **3b** (200 mg, 0.44 mmol) in DMF (3 mL) and the mixture was stirred for 24 h. The mixture was cooled to 0 °C and diluted with water (5 mL). The aqueous mixture was extracted with EtOAc (3 × 5 mL), the combined organic fractions were washed with water (2 × 10 mL), dried, and solvent was removed *in vacuo*. The resulting residue was purified by chromatography, eluting with a gradient of (50–100%) EtOAc/pet. ether, to give carbamate **5a** (190 mg, 73%): $^1$H NMR δ 7.97 (s, 1 H, H-4‴), 7.91 (s, 1 H, H-4), 7.52 (s, 1 H, H-7), 7.11 (d, *J* = 8.6 Hz, 1 H, H-6″), 6.89 (d, *J* = 2.9, 1 H, H-3″), 6.84 (dd, *J* = 8.6, 3.0 Hz, 1 H, H-5″), 5.19 (s, 2 H, 2‴-$CCH_2$), 4.55 (tt, *J* = 12.4, 4.3 Hz, 1 H, H-4′), 4.16 (dd, *J* =

11.6, 4.3 Hz, 2 H, H-2′, H-6′), 4.06 (s, 3 H, 1‴-$NCH_3$), 3.57 (td, $J$ = 11.9, 1.5 Hz, 2 H, H-2′, H-6′), 3.38 (s, 3 H, 3-$NCH_3$), 2.47 (app. qd, $J$ = 12.6, 4.5 Hz, 2 H, H-3′, H-5′), 2.24 (s, 3 H, 2″-$CCH_3$), 1.79 (dd, $J$ = 12.4, 2.6 Hz, 2 H, H-3′, H-5′), 1.44 (s, 9 H, $C(CH_3)_3$).

**3-Methyl-6-((2-methyl-4-((1-methyl-5-nitro-1*H*-imidazol-2-yl)methoxy)phenyl)amino)-1-(tetrahydro-2*H*-pyran-4-yl)-1,3-dihydro-2*H*-imidazo[4,5-c]pyridin-2-one (5).** TFA (0.38 mL, 5.0 mmol) was added to a solution of carbamate **5a** (150 mg, 0.25 mmol) in DCM (8 mL) and the resulting mixture was stirred for 24 h. The solvent was removed *in vacuo* and the residue taken up in EtOAc (10 mL), washed with saturated $NaHCO_3$ (10 mL), water (10 mL), saturated NaCl (10 mL) and dried. Solvent was removed *in vacuo* and the residue was purified by chromatography, eluting with 100% EtOAc, to give ether **5** (110 mg, 92%) as a yellow solid: mp 196–198 °C; $^1H$ NMR [$(CD_3)_2SO$] δ 8.08 (s, 1 H, H-4‴), 7.79 (s, 1 H, H-4), 7.70 (s, 1 H, NH), 7.55 (d, $J$ = 8.8 Hz, 1 H, H-6″), 6.93 (d, $J$ = 2.9 Hz, 1 H, H-3″), 6.85 (dd, $J$ = 8.8, 3.0 Hz, 1 H, H-5″), 6.74 (s, 1 H, H-7), 5.23 (s, 2 H, 2‴-$CCH_2$), 4.37 (tt, $J$ = 12.3, 4.2 Hz, 1 H, H-4′), 4.03–3.94 (m, 2 H, H-2′, H-6′), 3.96 (s, 3 H, 1‴-$NCH_3$), 3.46 (t, $J$ = 11.3 Hz, H-2′, H-6′), 3.27 (s, 3 H, 3-$NCH_3$), 2.32–2.16 (m, 2 H, H-3′, H-5′), 2.19 (s, 3 H, 2″-$CCH_3$), 1.66 (dd, $J$ = 12.2, 2.2 Hz, H-3′, H-5′); $^{13}C$ NMR [$(CD_3)_2SO$] δ 153.4, 153.3, 152.9, 148.7, 140.1, 136.7, 134.8, 132.0, 131.97, 125.6, 123.8, 121.7, 117.4, 112.9, 89.8, 66.9 (2), 62.9, 49.8, 34.1, 29.9 (2), 27.5, 18.8; HRMS calcd for $C_{24}H_{28}N_7O_5$ ($MH^+$) *m/z* 494.2146, found 494.2155 (+1.67 ppm). HPLC purity 99.9%

**1-(1-Methyl-2-nitro-1*H*-imidazol-5-yl)ethan-1-one (6).** Compound **6** was prepared as described previously[3]. HPLC purity 100.0%.

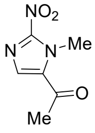


**6**

***N*3-Methyl-6-((2-methyl-4-(1-(5-nitrothiophen-2-yl)ethoxy)phenyl)amino)-1-(tetrahydro-2*H*-pyran-4-yl)-1,3-dihydro-2*H*-imidazo[4,5-*c*]pyridin-2-one (7).**

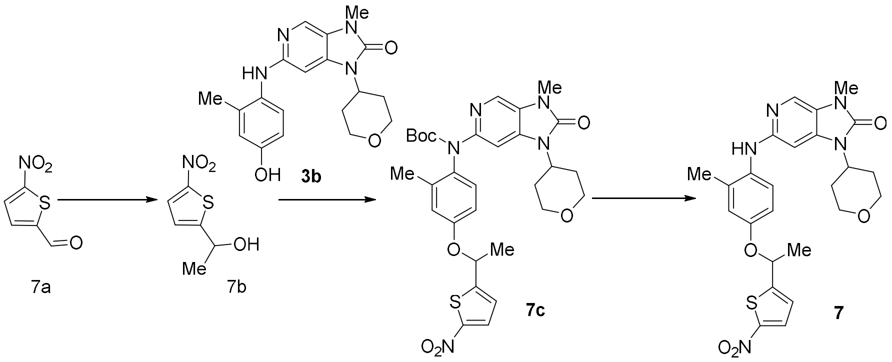


**1-(5-Nitrothiophen-2-yl)ethan-1-ol (7b).** $TiCl_4$ (1.74 mL, 15.9 mmol) was added dropwise to a solution of DCM (50 mL) at -78 °C. This was followed by slow addition of 3.0 M MeMgBr in diethyl ether (5.17 mL, 15.5 mmol). A solution of 5-nitrothiophene-2-carbaldehyde[4] **7a** (1.25 g, 7.95 mmol) in DCM (10 mL) was added to the mixture dropwise. The resulting mixture was allowed to stir at -78 °C for 2 hours. The reaction was quenched by slow addition of sat. $NH_4Cl$ at -78 °C and slowly warmed to 20 °C with continuous stirring. Sat. $NH_4Cl$ (50 mL) was added to the mixture, and the product was extracted with DCM (2 x 50 mL). The combined organic layers were washed with brine (50 mL), dried and the solvent evaporated *in vacuo*. The crude mixture was purified by column chromatography, eluting with DCM, to provide alcohol **7b** (1.26 g, 92%) as a brown oil: $^1H$ NMR δ 7.81 (d, $J$ = 4.2 Hz, 1 H, H-4), 6.89 (dd, $J$ = 4.2, 0.9 Hz, 1 H, H-3), 5.09–5.16 (m, 1 H, CH), 2.23 (d, $J$ = 4.9 Hz, 1 H, OH), 1.62 (d, $J$ = 6.5 Hz, 3 H, $CH_3$); HRMS calcd for $C_6H_8NO_3S$ ($MH^+$) *m/z* 174.0219, found 174.0220 (0.35 ppm).

***tert*-Butyl (3-methyl-2-oxo-1-(tetrahydro-2*H*-pyran-4-yl)-2,3-dihydro-1*H*-imidazo[4,5-*c*]pyridin-6-yl)(2-methyl-4-(1-(5-nitrothiophen-2-yl)ethoxy)phenyl)carbamate (7c).** A solution of DIAD (0.27

mL, 1.39 mmol) in THF (1 mL) was added to a mixture of phenol (417 mg, 0.92 mmol), alcohol **7b** (159 mg, 0.92 mmol) and triphenylphosphine (361 mg, 1.38 mmol) in THF (4 mL). The resulting mixture was allowed to stir at 20 °C for 21 h. The solvent was evaporated *in vacuo*. The crude mixture was purified by column chromatography, eluting with 65% EtOAc/pet. ether, to give a crude product **6c** containing triphenylphosphine oxide which was used without further purification in the next step: $^1$H NMR δ 7.91 (s, 1 H, H-4), 7.79 (s, *J* = 4.2 Hz, 1 H, H-4'''), 7.44-7.50 (m obscured by triphenylphosphine oxide, 1 H, H-7), 7.07 (d, *J* = 8.6 Hz, 1 H, H-6''), 6.94 (dd, *J* = 4.2, 0.7 Hz, 1 H, H-3'''), 6.82 (d, *J* = 2.9 Hz, 1 H, H-3''), 6.73 (dd, *J* = 8.6, 2.9 Hz, 1 H, H-5''), 5.52 (q, *J* = 6.4 Hz, 1 H, CH), 4.50–4.58 (m, 1 H, H-4'), 4.16 (dd, *J* = 12.0, 4.3 Hz, 2 H, H-2', H-6'), 3.57 (ddd, *J* = 12.0, 12.0, 1.4 Hz, 2 H, H-2', H-6'), 3.38 (s, 3 H, 3-$NCH_3$), 2.46 (tdd, *J* = 12.5, 12.5, 4.3 Hz, 2 H, H-3', H-5'), 2.21 (s, 3 H, 2''-Me), 1.79 (dd, *J* = 12.5, 2.7 Hz, 2 H, H-3', H-5'), 1.73 (d, *J* = 6.4 Hz, 3 H, $CH_3$), 1.42 (s, 9 H, $C(CH_3)_3$).

**3-Methyl-6-((2-methyl-4-(1-(5-nitrothiophen-2-yl)ethoxy)phenyl)amino)-1-(tetrahydro-2*H*-pyran-4-yl)-1,3-dihydro-2*H*-imidazo[4,5-*c*]pyridin-2-one (7).** TFA (0.53 mL, 6.93) was added to the crude carbamate **7c** (423 mg, 0.69 mmol) in DCM (10 mL) and stirred at 20 °C for 21 h. The resulting mixture was diluted with DCM (10 mL) and washed with sat. $NaHCO_3$ (20 mL). The organic layer was washed with brine (20 mL), dried and the solvent evaporated. The crude residue was purified by column chromatography, eluting with 1% MeOH/DCM, to give ether **7** (110 mg, 31%) as an orange solid: mp 175–176 °C; $^1$H NMR δ 7.81 (d, *J* = 4.2 Hz, 1 H, H-4'''), 7.78 (d, *J* = 0.5 Hz, 1 H, H-7), 7.29 (d, *J* = 8.7 Hz, 1 H, H-6''), 6.96 (dd, *J* = 4.2, 0.7 Hz, 1 H, H-3'''), 6.86 (d, *J* = 2.9 Hz, 1 H, H-3''), 6.78 (dd, *J* = 8.7, 2.9 Hz, 1 H, H-5''), 6.32 (d, *J* = 0.6 Hz, 1 H, H-7), 5.96 (br s, 1H, NH), 5.53 (q, *J* = 6.4 Hz, 1 H, CH), 4.41–4.49 (m, 1 H, H-4'), 4.05–4.09 (m, 2 H, H-2', H-6'), 3.50 (ddd, 12.0, 12.0, 1.7 Hz, 2 H, H-2', H-6'), 3.39 (s, 3 H, 3-$NCH_3$), 2.20–2.31 (m, 5 H, 2''-Me, H-3', H-5'), 1.76 (d, *J* = 6.4 Hz, 3 H, $CH_3$), 1.69–1.72 (m, 2 H, H-3', H-5'); $^{13}$C NMR δ 155.8, 153.9, 153.8, 153.0, 151.0, 137.2, 133.8, 133.6, 128.6, 126.2, 124.5, 123.1, 122.1, 119.2, 114.5, 88.3, 72.8, 67.5 (2), 50.1, 30.1 (2), 27.5, 24.1, 18.5; MS *m/z* 510.2 ($MH^+$, 100%); HRMS calcd for $C_{25}H_{28}N_5O_5S$ ($MH^+$) *m/z* 510.1806, found 510.1819 (2.62 ppm). HPLC purity 95.1%.

**1-(1-Methyl-2-nitro-1*H*-imidazol-5-yl)ethyl (4-methoxy-2-methylphenyl)(3-methyl-2-oxo-1-(tetrahydro-2*H*-pyran-4-yl)-2,3-dihydro-1*H*-imidazo[4,5-*c*]pyridin-6-yl)carbamate (8).**

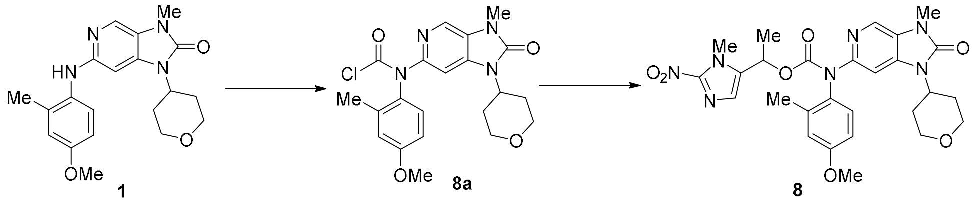


**(4-Methoxy-2-methylphenyl)(3-methyl-2-oxo-1-(tetrahydro-2*H*-pyran-4-yl)-2,3-dihydro-1*H*-imidazo[4,5-*c*]pyridin-6-yl)carbamic chloride (8a).** Triphosgene (320 mg, 1.09 mmol) was added to a suspension of imidazopyridinone **1** (400 mg, 1.09 mmol) and $NaHCO_3$ (180 mg, 2.18 mmol) in dry THF (20 mL) and the resulting mixture was stirred at 20 °C for 24 h. Dry $N_2$ was bubbled through the reaction mixture for 10 minutes, water (0.2 mL) was added and the mixture filtered through diatomaceous earth and solvent evaporated. The product was purified by chromatography, eluting with a gradient (50–100%) of EtOAc/pet. ether, to give carbamic chloride **8a** (250 mg, 53%) as a pale yellow foam which was used directly without further characterisation: $^1$H NMR [$(CD_3)_2SO$] δ 8.17 (s, 1 H, H-4), 7.86 (s, 1 H, H-7), 7.44 (d, *J* = 8.7 Hz, 1 H, H-6″), 6.89 (d, *J* = 2.8 Hz, 1 H, H-3″), 6.81 (dd, *J* = 8.7, 2.9 Hz, 1 H, H-5″), 4.49 (tt, *J* = 12.3, 4.1 Hz, 1 H, H-4′), 4.01 (dd, *J* = 12.1, 4.4 Hz, 2 H, H-2′, H-6′), 3.74 (s, 3 H, $OCH_3$), 3.49 (t, *J* = 11.2 Hz, 2 H, H-2′, H-6′), 3.37 (s, 3 H, 3-$NCH_3$), 2.44–2.31 (5H, m, 2″-$CCH_3$, H-3′, H-5′), 1.68 (dd, *J* = 12.0, 2.3 Hz, H-3′, H-5′).

**1-(1-Methyl-2-nitro-1*H*-imidazol-5-yl)ethyl (4-methoxy-2-methylphenyl)(3-methyl-2-oxo-1-(tetrahydro-2*H*-pyran-4-yl)-2,3-dihydro-1*H*-imidazo[4,5-c]pyridin-6-yl)carbamate (8).** 1-(1-Methyl-2-nitro-1*H*-imidazol-5-yl)ethan-1-ol[3] (47 mg, 0.28 mmol) and $Cs_2CO_3$ (90 mg, 0.28 mmol) was added to a solution of carbamic chloride **8a** (100 mg, 0.23 mmol) in DMF (2 mL) at 0 °C and the resulting mixture was stirred at room temperature for 24 h. The mixture was diluted with water (10 mL), and the resulting residue was extracted with EtOAc (3 × 10 mL), then the combined organic fractions were washed with water (2 × 10 mL), dried, and the solvent evaporated. The residue was purified by chromatography, eluting with 1% MeOH/DCM, to give carbamate **8** (24 mg, 18%) as a pale yellow solid: mp 172–175 °C; $^1$H NMR [$(CD_3)_2SO$] δ 8.06 (s, 1 H, H-4), 7.61 (s, 1 H, H-7), 7.19 (d, *J* = 8.7 Hz, H-6″), 7.14 (s, 1 H, H-4‴), 6.83 (d, *J* = 2.8 Hz, H-3″), 6.74 (dd, *J* = 8.6, 2.9 Hz, H-5″), 6.04 (q, *J* = 6.6 Hz, 5‴-CCH), 4.50 – 4.38 (m, 1 H, H-4′), 4.05–3.93 (m, 2 H, H-3′, H-5′), 3.84 (s, 3 H, 1‴-$NCH_3$), 3.73 (s, 3 H, 4″-$OCH_3$), 3.54–3.41 (m, 3 H, H-3′, H-5′), 3.31 (s, 3 H, 3-$NCH_3$), 2.39–2.25 (m, 2 H, H-2′, H-6′), 2.21 (s, 3 H, 2″-$CCH_3$), 1.70–1.61 (m, 2 H, H-2′, H-6′), 1.58 (d, *J* = 6.6 Hz, 3 H, 5‴-$C(CH_3)$. HRMS calcd for $C_{27}H_{32}N_7O_7$ ($MH^+$) *m/z* 566.2358, found 566.2350 (-1.32 ppm). HPLC purity 97.8%.

**1-(5-(4-Amino-7-methyl-7*H*-pyrrolo[2,3-*d*]pyrimidin-5-yl)indolin-1-yl)-2-(3-((1-methyl-2-nitro-1*H*-imidazol-5-yl)methoxy)phenyl)ethan-1-one (9).** Compound **9** was prepared as described previously[2]. HPLC purity 97.6%.

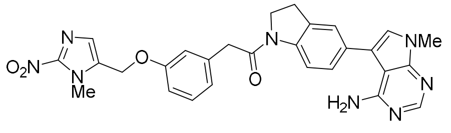


**9**

**1-(5-(4-Amino-7-methyl-7*H*-pyrrolo[2,3-*d*]pyrimidin-5-yl)indolin-1-yl)-2-(3-hydroxyphenyl)-ethan-1-one (10).** Compound **10** was prepared as described previously[2]. HPLC purity 96.6%.

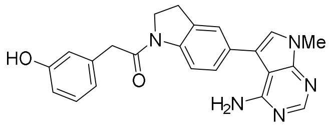


**10**

**6-((4-Hydroxy-2-methylphenyl)amino)-3-(methyl-*d*₃)-1-(tetrahydro-2*H*-pyran-4-yl)-1,3-dihydro-2*H*-imidazo[4,5-*c*]pyridin-2-one (11).**

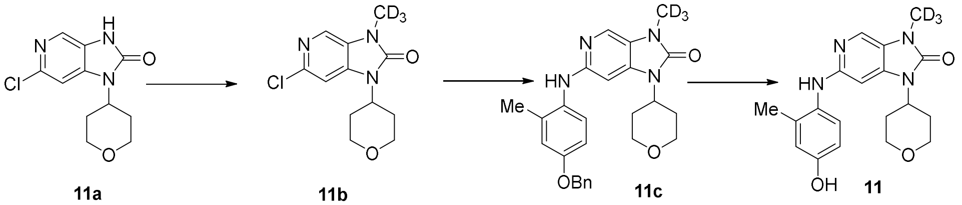


**11a** **11b** **11c** **11**

**6-Chloro-3-(methyl-*d*₃)-1-(tetrahydro-2*H*-pyran-4-yl)-1,3-dihydro-2*H*-imidazo[4,5-*c*]pyridin-2-one (11b).** $K_2CO_3$ (1.11 g, 8.04 mmol) was added to a stirred solution of pyridinone[1] **11a** (927 mg, 3.65 mmol) and $CD_3I$ (0.27 mL, 4.38 mmol) in dry DMF (10 mL) at 20 °C. The mixture was stirred at 20 °C for 16 h. The mixture was partitioned between EtOAc (100 mL) and water (50 ml). The organic fraction was washed with water (2 × 50 mL), washed with brine (50 mL), dried, filtered and the solvent evaporated. The residue was purified by chromatography, eluting with a gradient (50–100%) of EtOAc/pet. ether, to give chloride **11b** (0.77 g, 77%) as a white solid: mp 189 °C; $^1$H NMR δ 7.99 (d, *J* = 0.5 Hz, 1 H, H-4), 7.15 (d, *J* = 0.5 Hz, 1 H, H-7), 4.55 (tt, *J* = 12.5, 4.4 Hz, 1 H, H-4′), 4.15 (dd, *J* = 11.7, 4.7 Hz, 2 H, H-2′, H-6′), 3.55 (dt, *J* = 12.10, 1.9 Hz, 2 H, H-2′, H-6′), 2.38 (dq, *J* = 12.5, 4.7 Hz, 2 H, H-3′, H-5′), 1.77 (ddd, *J* = 12.5, 4.0, 1.5 Hz, 2 H, H-3′, H-5′); MS *m/z* 271.1 ($MH^+$, 100%), 273.1 ($MH^+$, 35%); HRMS calcd for $C_{12}H_{12}D_3{}^{35}ClN_3O_2$ ($MH^+$) *m/z* 271.1036, found 271.1026 (-3.6 ppm).

**6-((4-(Benzyloxy)-2-methylphenyl)amino)-3-(methyl-*d*₃)-1-(tetrahydro-2*H*-pyran-4-yl)-1,3-dihydro-2*H*-imidazo[4,5-*c*]pyridin-2-one (11c).** A degassed mixture of chloride **11b** (0.77 g, 2.84 mmol), 4-(benzyloxy)-2-methylaniline (0.61 g, 2.84 mmol), $Pd_2dba_3$ (0.13 g, 0.14 mmol), XPhos (0.27 g, 0.57 mmol) and $Cs_2CO_3$ (2.04 g, 6.26 mmol) in MeCN (10 mL) was stirred in a sealed tube at 105 °C for 22 h. The mixture was cooled, diluted with EtOAc (30 mL), filtered through diatomaceous earth and the filtrate was evaporated. The residue was purified by chromatography, eluting with EtOAc give imidazopyridinone **11c** (0.71 g, 56%) as a brown residue: $^1$H NMR δ 7.77 (d, *J* = 0.6 Hz, 1 H, H-4),

7.46–7.48 (m, 2 H, H-2''', H-6'''), 7.38–7.42 (m, 2 H, H-3''', H-5'''), 7.32 (m, 1 H, H-4'''), 7.24-7.27 (m, 1 H, H-6''), 6.93 (d, $J$ = 2.9 Hz, 1 H, 3''), 6.85 (dd, $J$ = 8.7, 2.9 Hz, 2 H, H-5''), 6.26 (d, $J$ = 0.6 Hz, 1 H, H-4), 5.93 (br s, 1 H, NH), 5.08 (s, 2 H, $CH_2$), 4.37–4.45 (m, 1 H, H-4'), 4.05–4.09 (m, 2 H, H-2', H-6'), 3.47–3.53 (m, 2 H, H-2', H-6'), 2.23–2.34 (m, 2 H, H-3', H-5'), 1.68–1.71 (m, 2 H, H-3', H-5').

**6-((4-Hydroxy-2-methylphenyl)amino)-3-(methyl-d$_3$)-1-(tetrahydro-2*H*-pyran-4-yl)-1,3-dihydro-2*H*-imidazo[4,5-*c*]pyridin-2-one (11).** A mixture of imidazopyridinone **11c** (101 mg, 0.23 mmol) and 2.0 M HCl (0.56 mL, 1.13 mmol) and 10% Pd/C (10 mg) in EtOH (20 mL) was stirred under $H_2$ (50 psi) for 24 h. Additional 10% Pd/C (10 mg) was added and the reaction mixture was stirred under $H_2$ (50 psi) for a further 24 h. The catalyst was removed by filtration through diatomaceous earth and the filtrate evaporated. The residue was triturated with EtOAc to obtain **11** (88 mg, 99%) as a light brown solid: mp 290 °C (decomp.); $^1$H NMR [$(CD_3)_2SO$] δ 12.86 (br s, 1 H, OH), 9.72 (br s, 2 H, $NH_2$), 9.58 (br s, 2 H, $NH_2$), 7.76 (s, 1 H, H-4), 7.08 (d, $J$ = 8.5 Hz, 1 H, H-6''), 6.79 (d, $J$ = 2.6 Hz, 1 H, H-3''), 6.77 (s, 1 H, H-7), 6.73 (dd, $J$ = 8.5, 2.6 Hz, 1 H, H-5''), 4.40 (m, 1 H, H-4'), 3.96 (dd, $J$ = 12.0, 4.3 Hz, 2 H, H-2', H-6'), 3.41–3.47 (m, 2 H, H-2', H-6'), 2.22 (dtd, $J$ = 12.0, 12.0, 4.3 Hz, 2 H, H-3', H-5'), 1.69 (dd, $J$ = 12.0, 2.7 Hz, 2 H, H-3', H-5'); $^{13}$C NMR [$(CD_3)_2SO$] δ 156.8, 153.3, 150.1, 141.7, 136.2, 128.0, 125.8, 121.9, 117.8, 114.2 (2), 88.1, 66.2 (2), 50.3, 29.1 (2), 27.0, 17.7; MS *m/z* 358.2 ($MH^+$, 100%). HPLC purity 97.9%.

*Bioanalytical methods*

**LC-MS analysis of hepatocyte and liver microsome samples**

Acetonitrile-quenched samples generated during metabolic stability studies in both hepatocytes and liver microsomes were analysed via LC-MS using the LC and MS conditions outlined below.

**LC conditions:**

Chromatography was achieved via a Waters Acquity UPLC system. For hepatocyte samples, an extended LC gradient was used to facilitate enhanced separation of potential polar metabolites.

| **Assay:** | Hepatocyte assay | Microsomal assay |
|---|---|---|
| **Column:** | Waters BEH C18 column (2.1 x 50 mm, 1.7 µm) | Ascentis Express C18 column (2.1 x 50 mm, 2.7 µm) |
| **LC conditions:** | Gradient cycle time: 8 minutes; Injection volume: 3 µL; Flow rate: 0.4 mL/min | Gradient cycle time: 4 minutes; Injection volume: 5 µL; Flow rate: 0.4 mL/min |
| **Mobile phase:** | Acetonitrile-water gradient with 0.05% formic acid | Acetonitrile-water gradient with 0.05% formic acid |

**MS conditions:**

| **Assay:** | Hepatocyte assay | Microsomal assay |
|---|---|---|
| **Instrument:** | Waters Xevo G2-S QToF | Waters Xevo G2 QTOF |
| **MS$^E$ detection:** | Quantitation of HAP **4** and DNA-PKi **2**: positive mode electrospray ionisation<br>Metabolite identification: positive and negative mode electrospray ionisation | Positive mode electrospray ionisation |
| **Instrument parameters:** | Sample cone voltage: 30 V<br>Low collision energy: 4 eV<br>High energy ramp: 4 – 50 eV | Sample cone voltage: 30 V |

**LC-MS/MS analysis of mouse liver S9, cell culture and tissue samples**

Deproteinised samples (10 µL) were injected from a refrigerated autosampler using an Agilent 1260 series HPLC coupled with a model 6460 MS/MS equipped with a jet stream ESI source (Agilent Technologies, USA) in positive ionisation mode. The source parameters were: gas and sheath gas temperature 300°C, gas flow 7 L/min, sheath gas flow 10 L/min, nebuliser pressure 45 psi, capillary voltage 3000V and nozzle voltage 500 V. The C18 column (Zorbax-SB, 2.1×50 mm, 5 µm) was thermostatted at 30 °C at a flow rate of 0.5 mL/min using a gradients of acetonitrile + 0.1% formic acid/water with a typical total run time of 5 min. MRM data acquisition was performed with the following transitions using a cell accelerator voltage of 7 V and analysed using Agilent software Agilent MassHunter Workstation Quantitative Analysis for QQQ (Version 10.2).

| Compound name | SN number | Precursor Ion (m/z) | Product Ion (m/z) | Dwell (ms) | Fragmentor (V) | Collision Energy (V) |
|---|---|---|---|---|---|---|
| Ritonavir | - | 721 | 296.1 | 50 | 188 | 12 |
| | | | 140 [a] | 50 | 188 | 60 |

| | | | | | | |
|---|---|---|---|---|---|---|
| Ketoconazole | - | 531 | 82.1 [a] | 50 | 162 | 52 |
| Midazolam | - | 326.09 | 291.1 [a] | 50 | 202 | 29 |
| | | | 249 | 50 | 202 | 41 |
| α-Hydroxy-midazolam | - | 342.08 | 324 [a] | 50 | 136 | 21 |
| | | | 168 | 50 | 136 | 41 |
| **1** | 39536 | 369.4 | 285 [a] | 50 | 152 | 28 |
| | | | 163 | 50 | 152 | 40 |
| **2** | 39872 | 355 | 271 [a] | 50 | 170 | 28 |
| | | | 163 | 50 | 170 | 40 |
| **3** | 39897 | 494 | 227 [a] | 25 | 160 | 64 |
| | | | 149 | 25 | 160 | 72 |
| **4** | 40458 | 508 | 354.1 [a] | 50 | 190 | 28 |
| | | | 271.1 | 50 | 190 | 40 |
| **4**-2 | - | 506 | 352.1 | 50 | 208 | 28 |
| | | | 268.1 | 50 | 208 | 48 |
| **5** | 40459 | 494 | 354.1 [a] | 50 | 180 | 28 |
| | | | 269 | 50 | 180 | 44 |
| | | | 255 | 50 | 180 | 52 |
| | | | 241 | 50 | 180 | 64 |
| **6** | 41321 | 170.1 | 124 | 50 | 101 | 16 |
| | | | 82.1 [a] | 50 | 101 | 24 |
| **7** | 41522 | 510.2 | 354.1 | 50 | 202 | 29 |
| | | | 241 [a] | 50 | 202 | 60 |
| **8** | 40425 | 566 | 413.1[a] | 50 | 157 | 16 |
| | | | 395.1 | 50 | 157 | 16 |
| | | | 285.1 | 50 | 157 | 44 |
| **9** | 38718 | 539.2 | 399.1 | 50 | 213 | 28 |
| | | | 265.1 [a] | 50 | 213 | 44 |
| **10** | 38302 | 400.2 | 266.1 [a] | 50 | 213 | 36 |
| | | | 264.1 | 50 | 213 | 48 |
| **11** | 40784 | 538.2 | 274.1 [a] | 50 | 136 | 29 |
| | | | 166.1 | 50 | 136 | 41 |

[a] major product

#### *Microscopy and image analysis of EdU and H33342-stained sections*

Images of sections from mouse ileum and tongue (Fig. S9 and S10) were acquired using an Olympus SLIDEVIEW VS200 slide-scanner equipped with a DAPI filter set for H33342 (excitation 365/20nm, emission 450/40nm), Cy5 filter set for EdU (excitation645/30 nm, emission 700/50 nm) and an UPLXAPO 20x/0.8 NA objective lens. The camera was a Hamamatsu ORCA-Flash4.0 V3 sCMOS camera.

Image analysis of EdU+ and H33342+ nuclei was carried out as previously described[5] with minor changes reflect the different slide-scanner used. In addition, the Triangle algorithm was used for thresholding rather than Otsu and the output montage scaling was reduced to enable easier evaluation of segmentation results.

Ileum and tongue images were exported from the propriety file format into a 2-channel TIFF file format without compression at the highest resolution level using a specific macro for the Olympus VSI files. A duplicate image was created from the EdU channel to serve as a background image for the image border in the same way as the previously-described macro[5]. In addition, the stack order was reversed in order to be consistent with the channel assignments. This resulted in a 3-channel stack that matched the previous data.

**Table S1.** DNA-PKi and HAPs: structures, kinase inhibition, purity and radiosensitisation of HAP1 cells by regrowth assay.

| Cmpd | Structure | Biochemical $IC_{50}$ (nM) | | | | | | | HAP1 radiosensitiation | | Purity | |
|---|---|---|---|---|---|---|---|---|---|---|---|---|
| | | DNA-PK | Prodrug /DNA-PKi Ratio[a] | PI3Ka (p110a/p85a) | PI3Kb (p110b/p85a) | PI3Kg (p110g) | PI3Kd (p110d/p85a) | mTOR/ FRAP | $S_{50}$ (μM) wildtype *PRKDC* [c] | $S_{50}$ (μM) *PRKDC* null [c] | % parent | % effector |
| 1 | 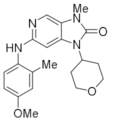 | 7.95 | - | 1840 [231][b] | 8830 [1110] | 10700 [1350] | 6820 [858] | 31300 [3940] | 0.24 ± 0.04 | >10 | 99.6 | na[f] |
| 2 | 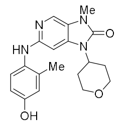 | 13.2 | - | 3217 [244] | 10900 [826] | 2366 [179] | 3589 [272] | 2190 [166] | 1.05 ± 0.15 | >10 | 99.1 | na |
| 3 | 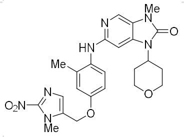 | 780 | 59 | 16175 | nd | nd | | 92915 | >10 | >10 | 97.1 | 0.18 |
| 4 | 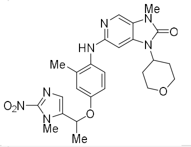 | 448 | 34 | 5854.5 [13] | 30670 [69] | 18,380 [41] | 6050 [14] | >100,000 [>223] | nd[g] | nd | 99.1 | no[h] |
| 5 | 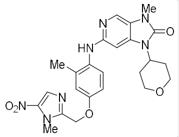 | 207 | 16 | 8653.5 [42] | 5329.5 [26] | 21,235 [103] | 18,620 [90] | 85,840 [415] | nd | nd | 99.9 | no |
| 7 | 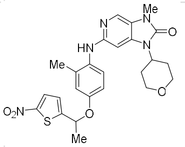 | nd | - | nd | nd | nd | nd | nd | nd | nd | 95.1 | 0.18 |
| 8 | 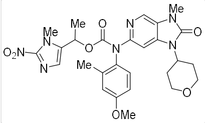 | 350 | 26 | nd | nd | nd | nd | nd | nd | nd | 97.8 | 0.42 |

Footnotes. [a] Ratio of prodrug DNA-PK $IC_{50}$/DNA-PKi $IC_{50}$. [b] Ratio = $IC_{50}$/DNA-PK IC50. [c] $S_{50}$ values are the means of three independent experiments. [d] $IC_{50}$ values are means of two independent experiments. [e] $IC_{50}$ values are means of two replicates in a single experiment. [f] Not applicable. [g] Not determined. [h] Not observed.

**Table S2.** Radiosensitisation by DNA-PK inhibitors: clonogenic assay data.

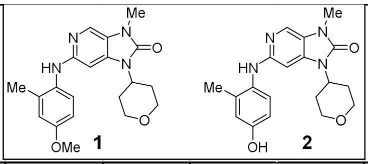


| Cmpd | Cell line | Gas phase | Conc (μM) | N expts | $D_{10}$ [a](Gy) | | $OER_{10}$ [b] | | SF drug only[c] | | $SER_{10}$ [d] | |
|---|---|---|---|---|---|---|---|---|---|---|---|---|
| | | | | | Mean | SEM | Mean | SEM | Mean | SEM | Mean | SEM |
| **1**[e] | HCT116 | Oxia | 0 | 6 | 5.86 | 0.14 | | | 1 | | | |
| | | | 0.33 | 3 | 3.77 | 0.44 | | | | | 1.55 | 0.14 |
| | | | 1 | 6 | 2.46 | 0.12 | | | | | 2.54 | 0.12 |
| | | | 3 | 3 | 1.58 | 0.09 | | | | | 3.65 | 0.16 |
| | | Hypoxia | 0 | 3 | 13.1 | 0.6 | | | 1 | | | |
| | | | 1 | 3 | 5.42 | 0.25 | | | 1 | 0.56 | 2.42 | 0.01 |
| **2** | HCT116 | Oxia | 0 | 5 | 5.47 | 0.39 | 2.58 | 0.21 | 1 | | | |
| | | | 0.33 | 1 | 4.85 | | | | 0.77 | - | 1.37 | - |
| | | | 1 | 5 | 2.52 | 0.29 | | | 0.918 | 0.087 | 2.24 | 0.20 |
| | | | 3 | 1 | 2.28 | | | | 0.63 | - | 2.91 | - |
| | | Hypoxia | 0 | 5 | 14.10 | 0.56 | | | 1 | | | |
| | | | 1 | 5 | 6.06 | 0.19 | | | 0.997 | 0.036 | 2.33 | 0.06 |
| | MiaPaca-2 | Oxia | 0 | 1 | 5.95 | | 2.43 | | 1 | | | |
| | | | 1 | 1 | 2.29 | | | | 1.09 | - | 2.60 | - |
| | | Hypoxia | 0 | 1 | 14.4 | | | | 1 | | | |
| | | | 1 | 1 | 5.6 | | | | 1.04 | - | 2.58 | - |
| | PANC-1 | Oxia | 0 | 2 | 7.11 | 0.74 | 2.66 | 0.49 | 1 | | | |
| | | | 1 | 2 | 4.32 | 0.72 | | | 0.870 | 0.081 | 1.66 | 0.22 |
| | | Hypoxia | 0 | 2 | 18.90 | 2.89 | | | 1 | | | |
| | | | 1 | 2 | 9.42 | 4.05 | | | 1.02 | 0.05 | 2.07 | 0.29 |
| | FaDu | Oxia | 0 | 3 | 9.24 | 0.32 | 2.13 | 0.20 | 1 | | | |
| | | | 1 | 3 | 2.72 | 0.53 | | | 0.96 | 0.14 | 3.61 | 0.55 |
| | | Hypoxia | 0 | 2 | 19.67 | 1.76 | | | 1 | | | |
| | | | 1 | 2 | 10.03 | 0.38 | | | 1.03 | 0.16 | 1.96 | 0.1 |
| | BxPC3 | Oxia | 0 | 2 | 9.27 | 0.51 | 2.69 | 0.17 | 1 | | | |
| | | | 1 | 2 | 8.63 | 0.44 | | | 0.98 | 0.08 | 1.074 | 0.002 |
| | | Hypoxia | 0 | 2 | 24.91 | 0.68 | | | 1 | | | |
| | | | 1 | 2 | 22.56 | 0.64 | | | 1.08 | 0.05 | 1.10 | 0.03 |
| | UT-SCC-74B | Oxia | 0 | 1 | 5.21 | | | | 1 | | | |
| | | | 1 | 1 | 2.23 | | | | 0.77 | - | 2.34 | - |

Footnotes: [a] Dose for 10% survival, interpolated using the linear-quadratic model. [b] Oxygen enhancement ratio = $D_{10}$ hypoxia/$D_{10}$ oxia, radiation alone. [c] Surviving fraction for drug alone = plating efficiency with drug/plating efficiency without drug, [d] Sensitiser enhancement ratio = $D_{10}$ without drug/$D_{10}$ with drug. [e] Oxic data from Reference 1.

**Table S3**. Radiosensitisation by ether-linked HAPs of DNA-PK inhibitors: clonogenic assay data.

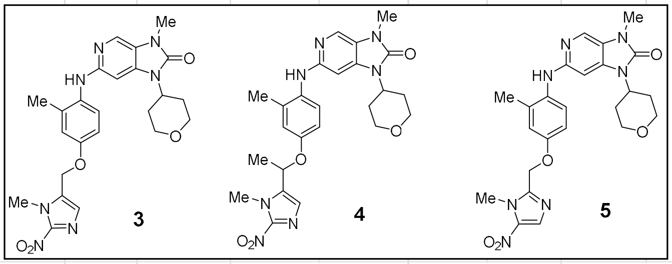

| Cmpd | Cell line | Gas phase | Conc (μM) | N expts | Plating efficiency | | [a]SF (drug only) | | [b]$D_{10}$ (Gy) | | [c]$SER_{10}$ | | p [d] |
|---|---|---|---|---|---|---|---|---|---|---|---|---|---|
| | | | | | Mean | SEM | Mean | SEM | Mean | SEM | Mean | SEM | |
| **3** | HCT116 | Oxia | 0 | 3 | 0.67 | 0.04 | | | 5.49 | 0.15 | | | |
| | | | 1 | 3 | 0.67 | 0.04 | 1.00 | 0.03 | 5.40 | 0.72 | 1.04 | 0.10 | |
| | | Hypoxia | 0 | 3 | 0.57 | 0.05 | | | 14.38 | 0.97 | | | |
| | | | 1 | 3 | 0.61 | 0.06 | 1.08 | 0.01 | 9.47 | 0.25 | 1.52 | 0.07 | 0.003 |
| **4** | HCT116 | Oxia | 0 | 5 | 0.97 | 0.16 | | | 4.36 | 0.31 | | | |
| | | | 1 | 5 | 0.92 | 0.16 | 0.93 | 0.07 | 4.03 | 0.28 | 1.09 | 0.05 | |
| | | Hypoxia | 0 | 4 | 0.88 | 0.18 | | | 12.64 | 0.53 | | | |
| | | | 1 | 4 | 0.95 | 0.20 | 1.08 | 0.08 | 9.54 | 0.69 | 1.36 | 0.15 | 0.063 |
| | FaDu | Oxia | 0 | 3 | 0.71 | 0.16 | | | 9.24 | 0.32 | | | |
| | | | 1 | 3 | 0.97 | 0.11 | 1.52 | 0.41 | 8.88 | 0.53 | 1.04 | 0.03 | |
| | | Hypoxia | 0 | 4 | 0.68 | 0.13 | | | 19.56 | 2.09 | | | |
| | | | 1 | 4 | 0.67 | 0.13 | 1.04 | 0.11 | 14.89 | 1.48 | 1.31 | 0.02 | <0.001 |
| **5** | HCT116 | Hypoxia | 0 | 1 | 0.63 | - | | | 13.4 | - | | | |
| | | | 1 | 1 | 0.56 | - | 0.88 | - | 12.5 | - | 1.07 | - | |

Footnotes: [a] Surviving fraction for drug alone = plating efficiency with drug/plating efficiency without drug, [b] Radiation dose for 10% survival, interpolated using the linear-quadratic model. [c]Sensitiser enhancement ratio = $D_{10}$ without drug/$D_{10}$ with drug.[d] p value for difference in $SER_{10}$ between oxia and hypoxia by one-way ANOVA.

**Table S4.** Intrinsic clearance of HAP **4**, yields of DNA-PKi **2** and effect of pan-CYPi in oxic hepatocytes[a]

| **Species** | **HAP 4, $CL_{int}$ (µL/min/$10^6$ viable cells)** | **Yield of 2 (%)[b]** | **% inhibition by pan-CYPi** | |
|---|---|---|---|---|
| | | | Loss of **4** | Formation of **2** |
| Human | 18.4 (15.9 – 20.9)[c] | 33.9 | 93% | 92% |
| Dog | 24.2 (22.5 – 25.9) | 38.2 | 94% | 99% |
| Rat | 66.6 (62.9 – 70.3) | 34.0 | 97% | 100% |
| Mouse | 307 (107 – 506) | 82.0 | 99% | 76% |

[a] Concentration-time profiles are shown in Fig. 5e. [b] Estimated by linear extrapolation of the log concentration plots to time zero. [c] Mean (95% confidence intervals).

**Table S5**. Metabolites of HAP **4** (0.5 μM) identified in oxic mouse, rat, dog and human hepatocytes ($5x10^5$ cells/mL) by mass spectrometry.

| **Structure/proposed biotransformation** | **Exact mass (Da)** | **Metabolite code** | **Rt (min)** | **Detection in hepatocytes** | |
|---|---|---|---|---|---|
| | | | | **No Inhibitor** | **Pan-CYPi** |
| **4** (parent/prodrug) | 507.223 | | 2.81 | human, dog, rat & mouse | human, dog, rat & mouse |
| **2** (active drug) | 354.1692 | | 2.07 | human, dog, rat & mouse | human, dog, rat & mouse |
| *N*-Glucuronidation of Secondary Amines | 683.2551 | **4**+176 | 2.56, 2.69 | ND[a] | Human |
| Hydroxylation / Mono-oxygenation | 523.2179 | **4**+16 | 2.51, 2.59 | human & dog | ND |
| Desaturation | 505.2073 | **4**-2 | 2.84 | human, dog, rat & mouse | Rat |
| 1-(1-methyl-2-nitro-1H-imidazol-5-yl) ethan-1-ol (**11**) | 171.0644 | **4**-336 | 1.6 | human, dog & rat | ND |
| 1-(1-methyl-2-nitro-1H-imidazol-5-yl) ethan-1-one (**6**) | 169.0487 | **4**-338 | 2.06 | human, dog, rat & mouse | ND |
| Oxidation of amino phenol to quinone-imine | 352.1535 | **2**-2 | 2.17 | human, dog & mouse | ND |
| Glucuronidation of aromatic phenol | 530.2013 | **2**+176 | | human, dog, rat & mouse | ND |
| Sulfation of aromatic alcohol | 434.126 | **2**+80 | | human, dog, rat & mouse | ND |
| Hydroxylation of tetrahydropyran | 370.1641 | **2**+16 | | human, dog, rat & mouse | ND |
| Glutathione S-conjugation | 659.2374 | **2**+305 | | rat & mouse | ND |

[a]Not detected.

**Table S6.** Sources of cell lines, and cell culture media.

| Cell line | Description | Source | Culture medium | Incubation time for clonogenic assays (d) |
|---|---|---|---|---|
| BxPC-3 | Human pancreatic ductal adenocarcinoma | ATCC | αMEM + 5% FBS | 14 |
| FaDu | Human squamous cell carcinoma (hypopharyngeal tumour) | ATCC | αMEM + 5% FBS | 12 |
| HAP1 | C631 subline of the near-haploid line derived from a human chronic myeloid leukaemia[a] | Horizon Discovery | IMDM + 5% FBS | |
| HAP1/*PRKDC*$^{-/-}$ | *PRKDC* knockout of HAP1[a] | Horizon Discovery | IMDM + 5% FBS | |
| HCT116/ATCC | human colorectal carcinoma | ATCC | αMEM + 5% FBS | |
| HCT116 | Subline of HCT116 | This lab [b] | αMEM + 5% FBS | 10 |
| HCT116/POR | Derived from HCT116/ATCC, with forced expression of cytochrome P450 reductase (POR). | This lab [c] | αMEM + 5% FBS + 2 μM puromycin | |
| MiaPaCa-2 | Human pancreatic ductal adenocarcinoma | ATCC | αMEM + 5% FBS | 8 |
| Panc-1 | Human pancreatic ductal adenocarcinoma | ECACC | αMEM + 5% FBS | 12 |
| UT-SCC-74B | Human squamous cell carcinoma (nodal metastasis) | Prof. Bradly G. Wouters, University of Toronto[d] | MEM + 10% FBS + 4.5 g/L D-glucose + 20 mM HEPES | 10 |

Footnotes: [a] The initially haploid HAP1 cell lines were passaged, before use, until they spontaneously reverted to diploidy. [b] Reference 6. [c] Reference 7. [d] Reference 8.

**Table S7**. Sources of reference substrates, metabolites and inhibitors.

| Compound | Use | Source |
|---|---|---|
| 1-Aminobenzotriazole (1-ABT) | Pan-CYP inhibitor | Sigma Aldrich |
| Tienilic acid (TA) | CYP2C9 inhibitor | NZ Scientific Support Ltd |
| Ketoconazole (Keto) | CYP3A4/5 inhibitor | Sigma Aldrich |
| Ritonavir (Rito) | CYP3A4/5 inhibitor | AK Scientific, USA |
| Furafylline | CYP1A2 inhibitor | Cayman Chemical |
| 2-Phenyl-2-(1-piperidinyl) propane (PPP) | CYP2B6 inhibitor | Cayman Chemical |
| Montelukast | CYP2C8 inhibitor | USP |
| Sulfaphenazole | CYP2C9 inhibitor | Sigma Aldrich |
| (S)-*N*-3-benzylnirvanol | CYP2C19 inhibitor | Toronto Research Chemicals |
| Quinidine | CYP2D9 inhibitor | Sigma Aldrich |
| Midazolam | CYP3A substrate | PM Separations NZ Ltd |
| α-Hydroxymidazolam | Reference metabolite | PM Separations NZ Ltd |

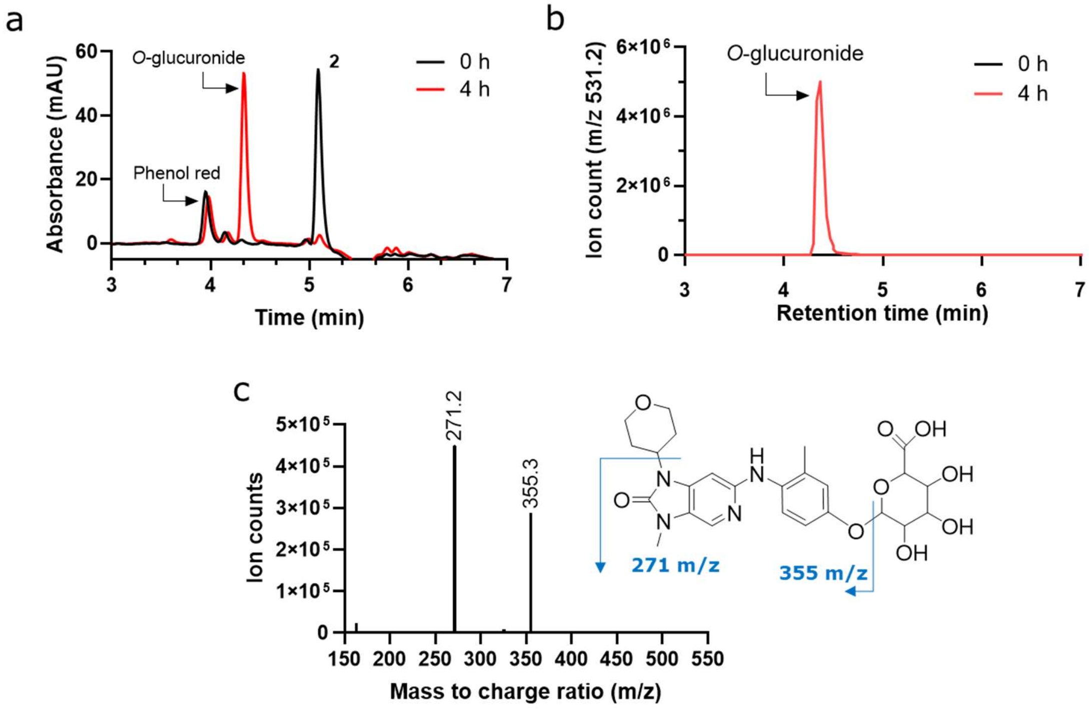


**Fig. S1.** Evidence for formation of the *O*-glucuronide metabolite BxPC-3 cultures incubated with DNA-PKi **2** for 4 h in the experiment shown in Fig. 2f. **a** Diode array absorbance chromatogram (250 nm, bandwidth 4 nm; reference 550 nm, bandwidth 4 nm) after incubation for 0 or 4 h. **b** Extracted ion chromatogram, m/z 531.2. **c** Collision-induced MS/MS spectrum for retention time 4.35 min peak in b.

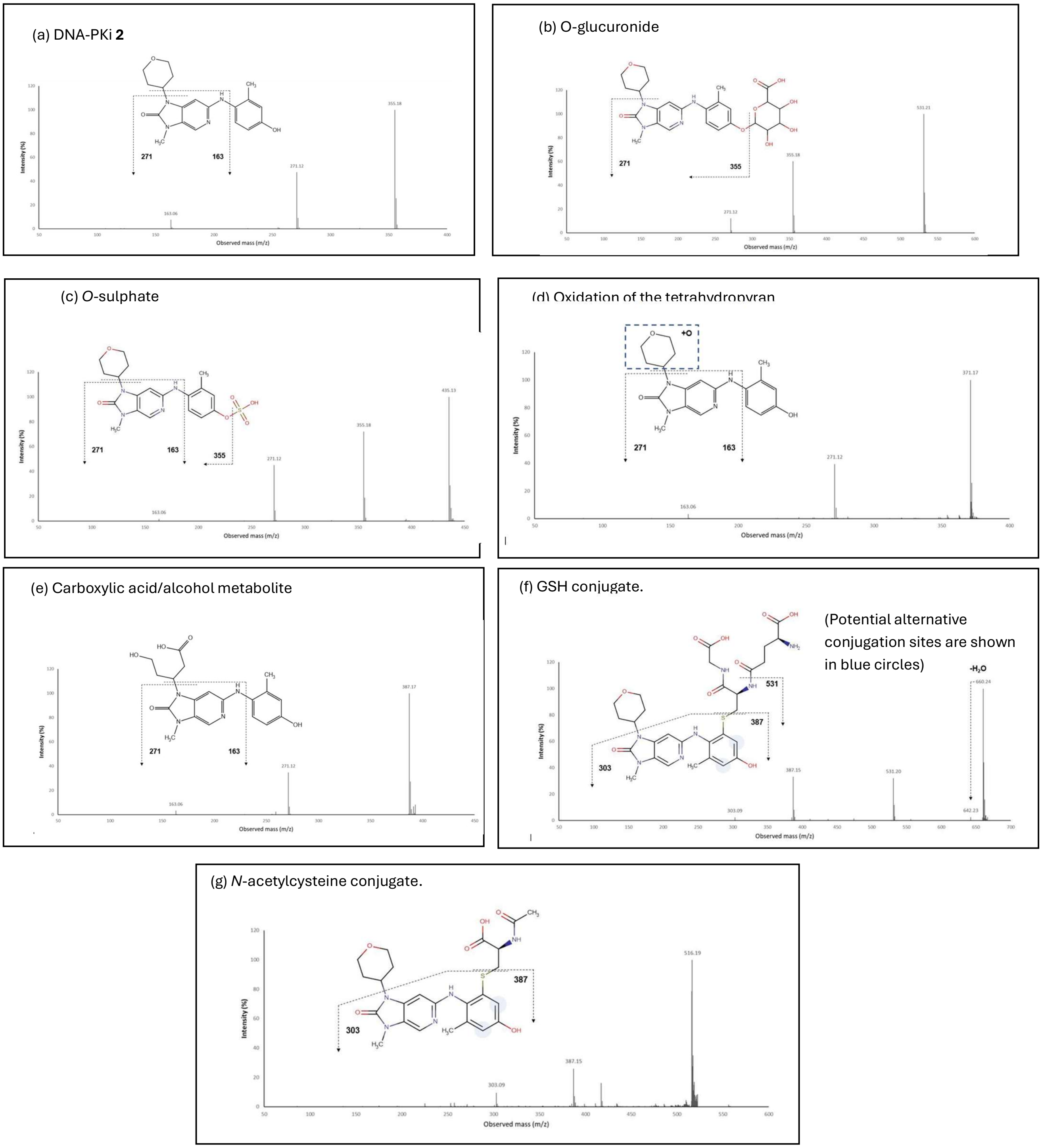


**Fig. S2.** Collision-induced dissociation of metabolites of DNA-PKi **2** incubated at 2 μM with hepatocytes. (a) Parent DNA-PKi **2** in rat hepatocytes, incubated for 2 min. Retention time (Rt) 3.71 min. (b) Human hepatocytes incubated with **2** for 180 min. Species at Rt 3.28 min, identified as the *O*-glucuronide. (c) Dog hepatocytes incubated with **2** for 240 min. Species at Rt 3.88 min, identified at the *O*-sulphate. (d) Rat hepatocytes incubated with **2** for 120 min. Species at Rt = 3.24 min, identified as resulting from hydroxylation of the tetrahydropyran ring. (e) Rat hepatocytes incubated with **2** for 240 min. Species at Rt = 2.80 min, identified as resulting from further oxidation of the tetrahydropyran ring. (f) Mouse hepatocytes incubated with **2** for 240 min. Species at Rt 3.40 min, identified as a GSH conjugate. (g) Mouse hepatocytes incubated with **2** for 240 min. Species at Rt = 3.80 min, identified as an *N*-acetylcysteine conjugate.

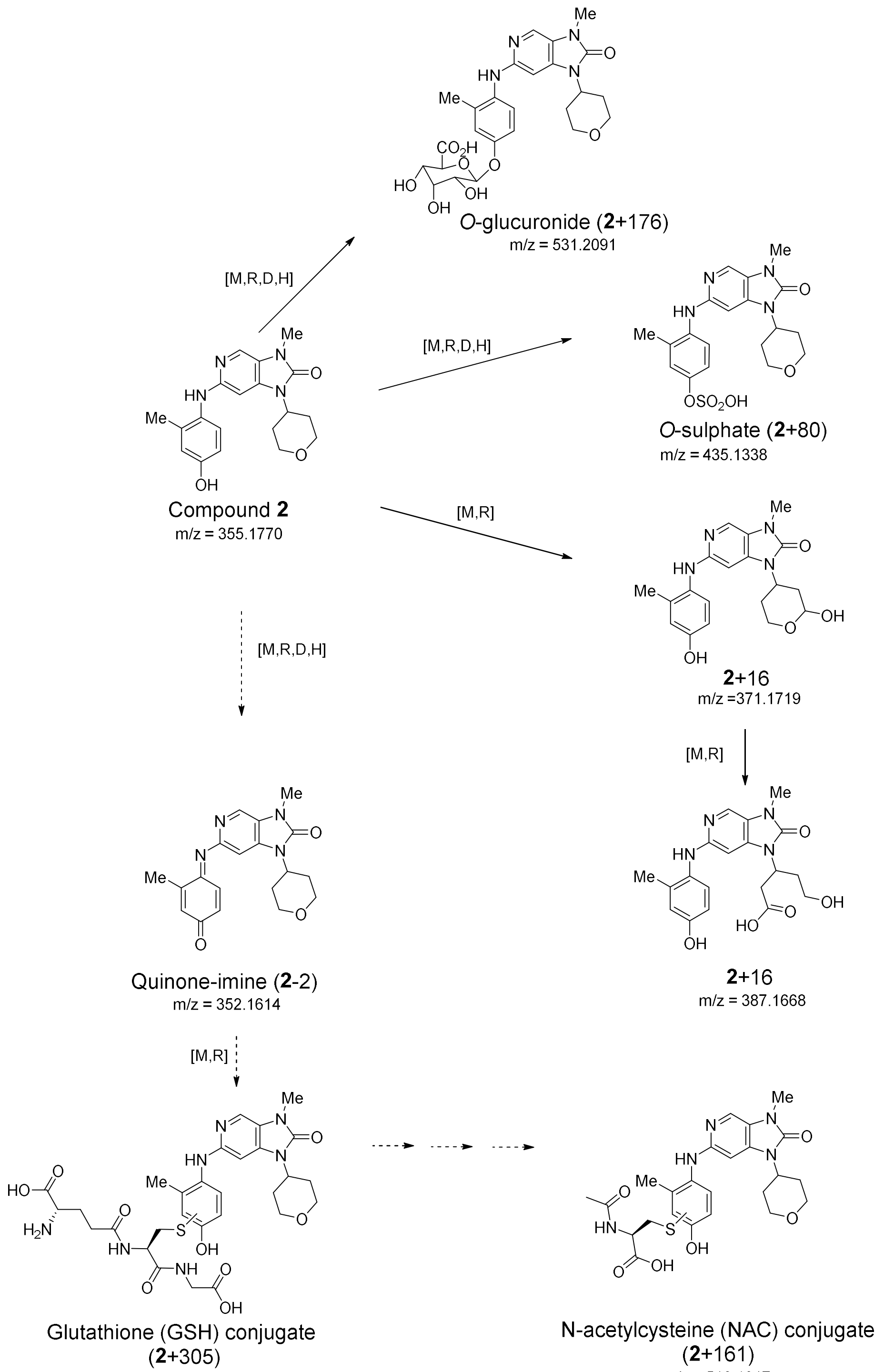


**Fig. S3.** Inferred metabolic pathway of metabolism of DNA-PKi **2**, supported by mass spectrometry identification of metabolites in mouse [M], rat [R], dog [D] and human [H] hepatocytes. High mass resolution m/z values were obtained with positive-mode electrospray ionisation. Except for the quinone-imine, identifications were supported by collision-induced dissociation MS/MS spectra (Fig. S2).

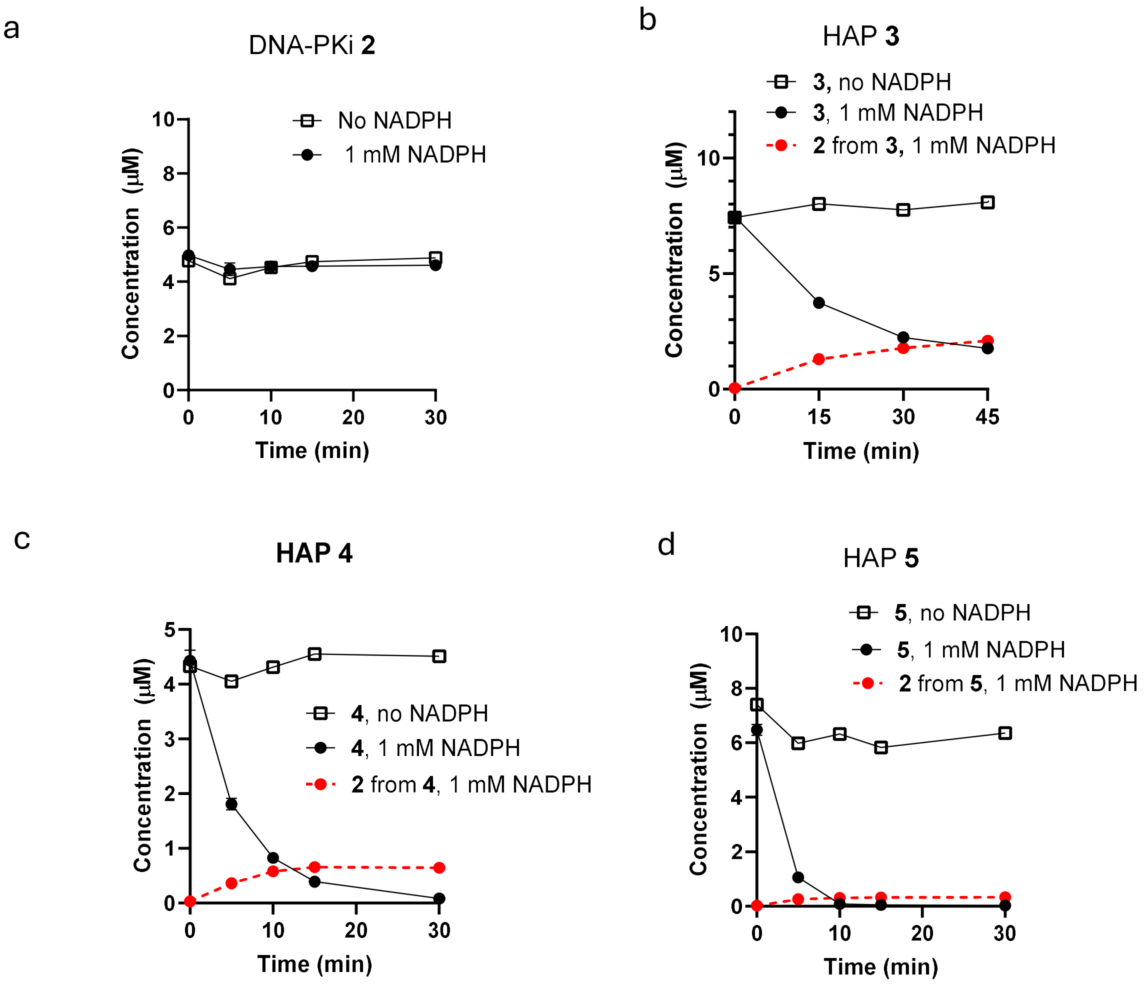


**Fig. S4.** Metabolism of DNA-PKi **2** and HAPs **3-5** in oxic mouse liver S9 (1 mg protein/mL) with 1 mM NADPH ( N=3 technical replicates) or without NADPH (N=1).

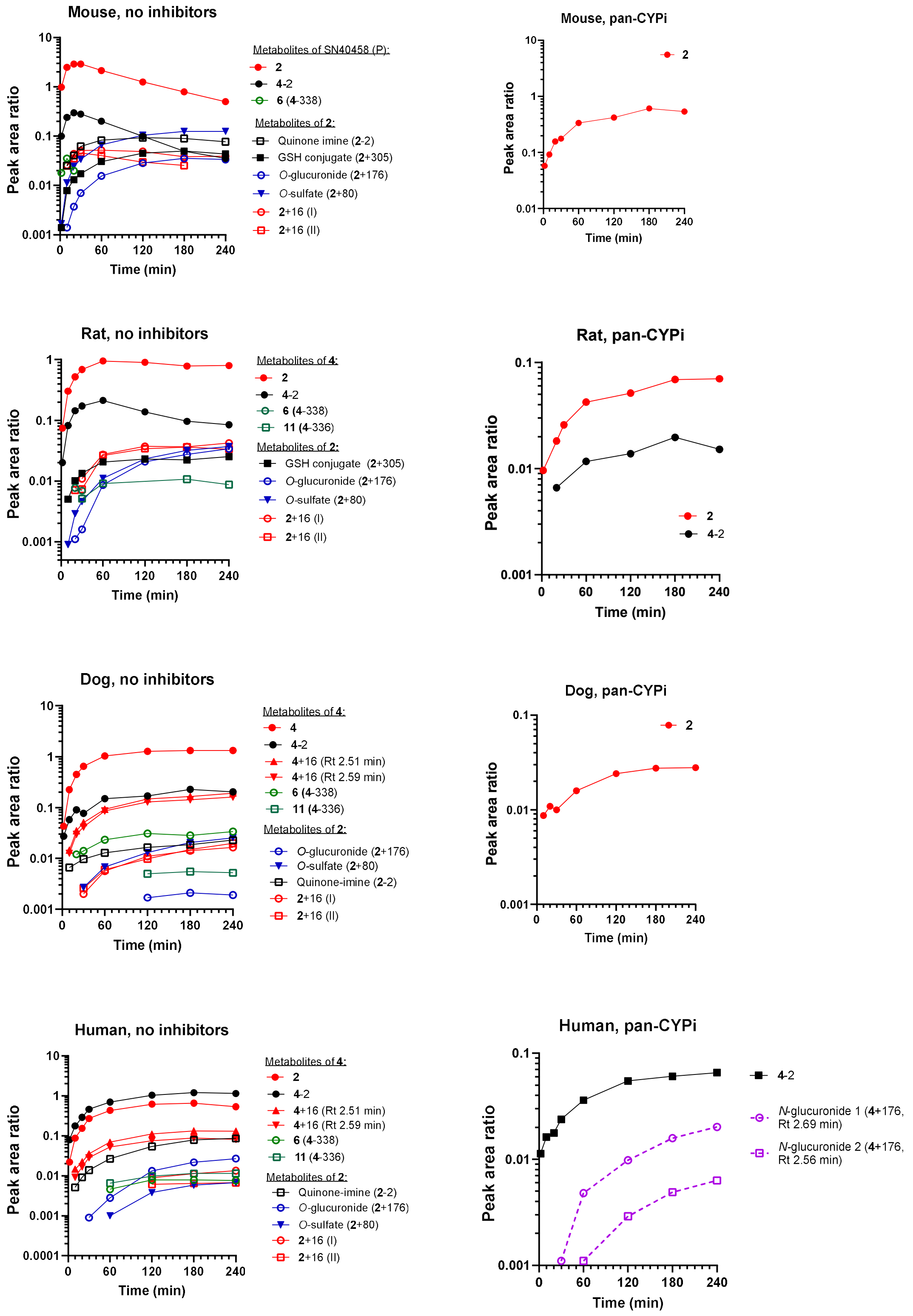


**Fig. S5:** Metabolite profile of HAP **4** (0.5 μM), with and without pan-CYPi, in oxic mouse, rat, dog and human hepatocytes ($5 \times 10^5$ cells/mL). The ordinate is the peak area ratio relative to internal standard metolazone.

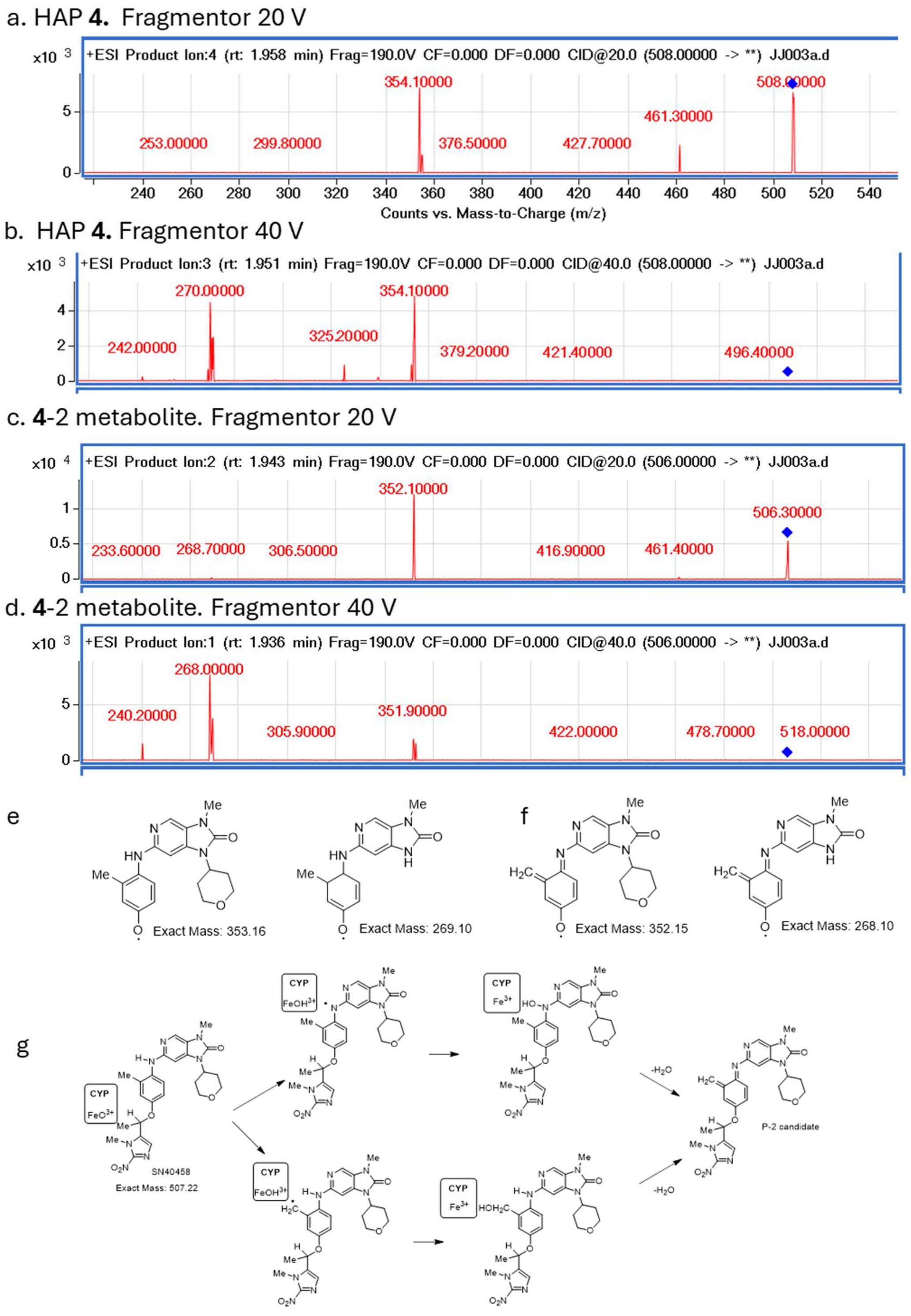


**Fig. S6.** Provisional identification of **4**-2, a -2Da metabolite of **4.** Collision-induced dissociation of HAP **4** at a fragmentor voltage of 20 V (a) and 40 V (b) generating fragments consistent with the structures in e. Collision-induced dissociation of **4**-2 under the same conditions (c,d) yielded fragments with m/z 2 Da lower, suggesting loss of 2H from the core structure as in f. The proposed CYP-mediated biotransformation pathway is shown in g.

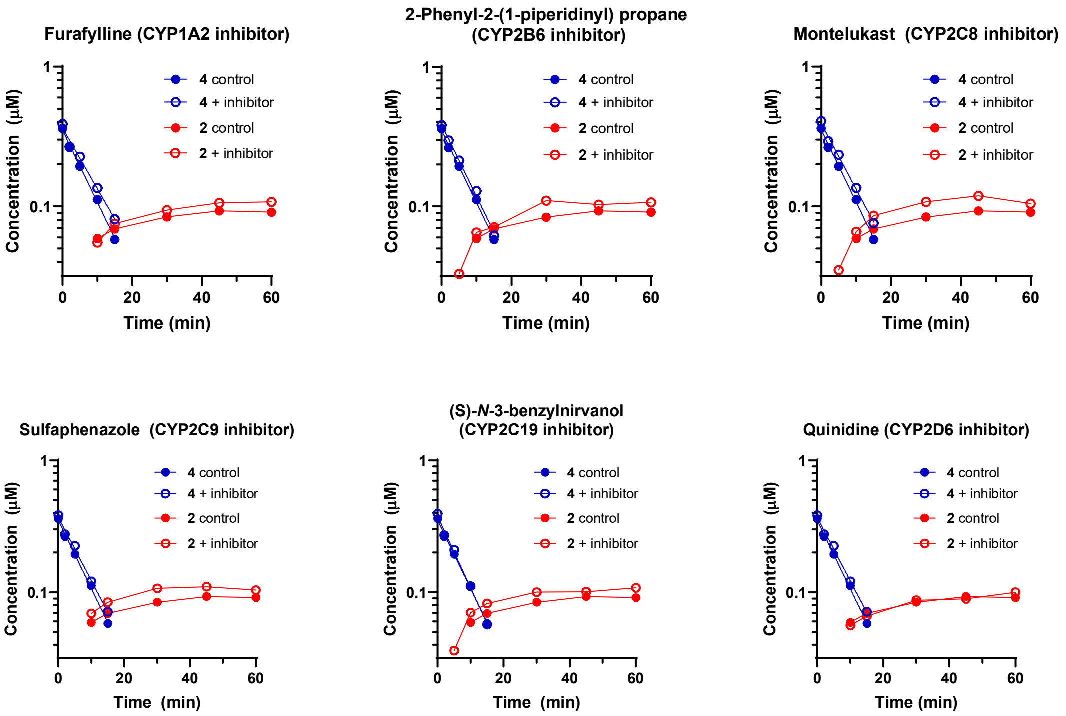


**Fig. S7.** Metabolism of **4** (0.5 µM) in oxic human liver microsomes (1 mg protein/mL1.3 mM NADPH) is not sensitive to the following isoform-specific CYP inhibitors: Furafylline, 125 µM; 2-phenyl-2-(1-piperidinyl) propane, 75 µM; montelukast, 10 µM; sulfaphenazole, 15 µM; (S)-*N*-3-benzylnirvanol, 15 µM and quinidine, 2 µM.

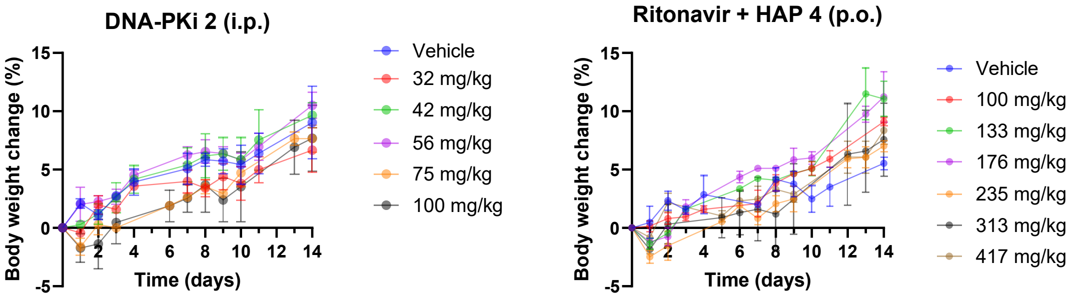


**Fig. S8.** Tolerability of single doses of HAP **4** + ritonavir (25 mg/kg 1 h prior), both orally by gavage, and **2** (i.p.) in male CD-1 mice (initial body weights 30-40 g). No clinical signs of toxicity were observed in any animals.

**Fig. S9**. Transverse sections of the ileum of CD-1 mice, stained for EdU (gold) and counterstained with H33342 (blue). Mice were dosed with EdU (50 mg/kg, i.p.) 4 days after treatment and euthanised 2 h later. Formalin-fixed paraffin embedded sections (10 µm) were cleared and stained for EdU by conjugation with AlexaFluor 647 using click chemistry. 4 animals/group, except for HAP **4** + Rito (N=3) because EdU labelling was not observed in ileum or tongue of in one mouse (presumed failed dosing with EdU).

A: Untreated controls

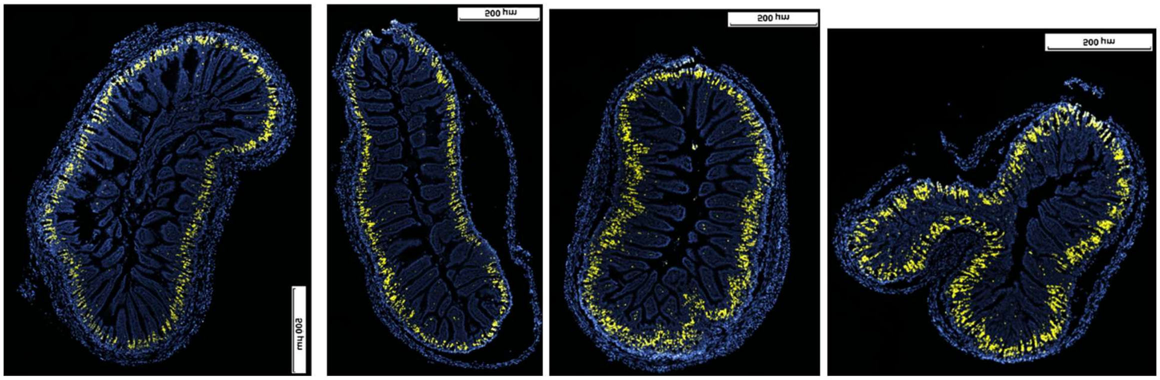

B: DNA-PKi **2**, 100 mg/kg, i.p.

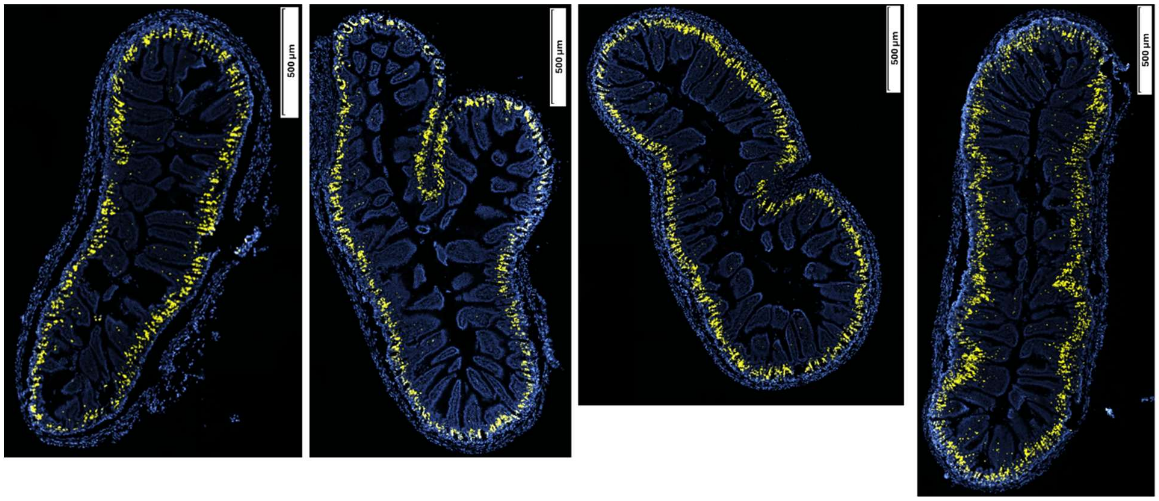


C: Rito (25 mg/kg, p.o) + HAP **4** (400 mg/kg, p.o.)

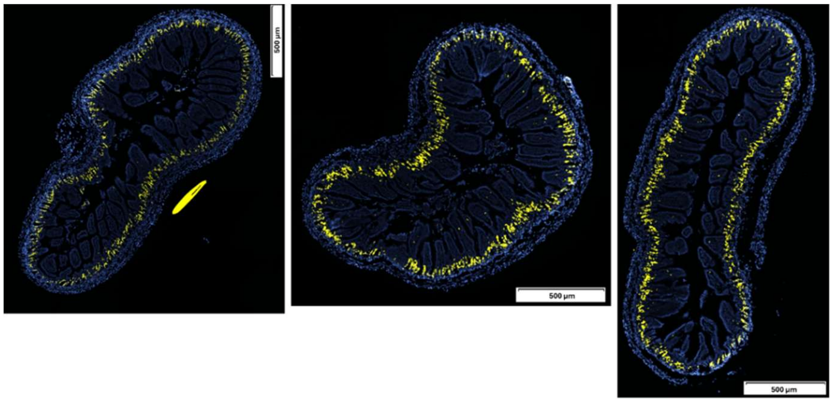

D: 10 Gy

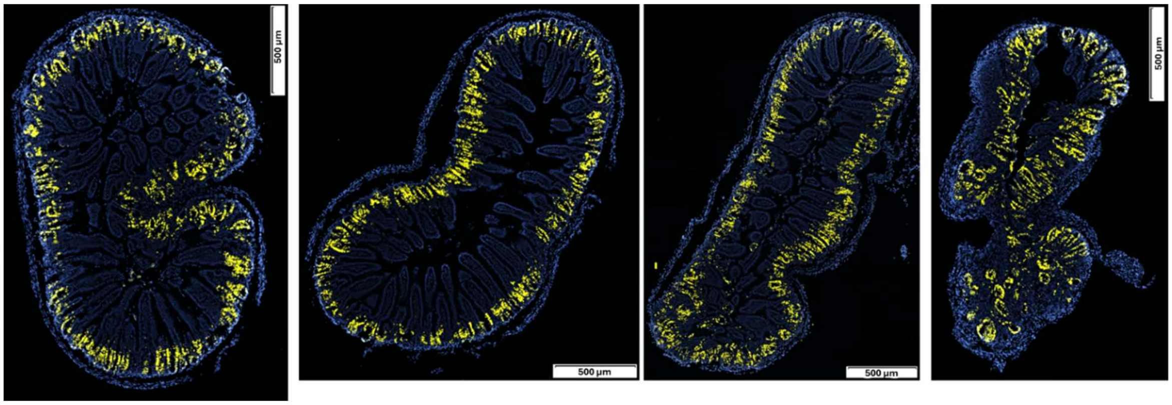


E: 10 Gy + DNA-PKi **2** (10 mg/kg, i.p.)

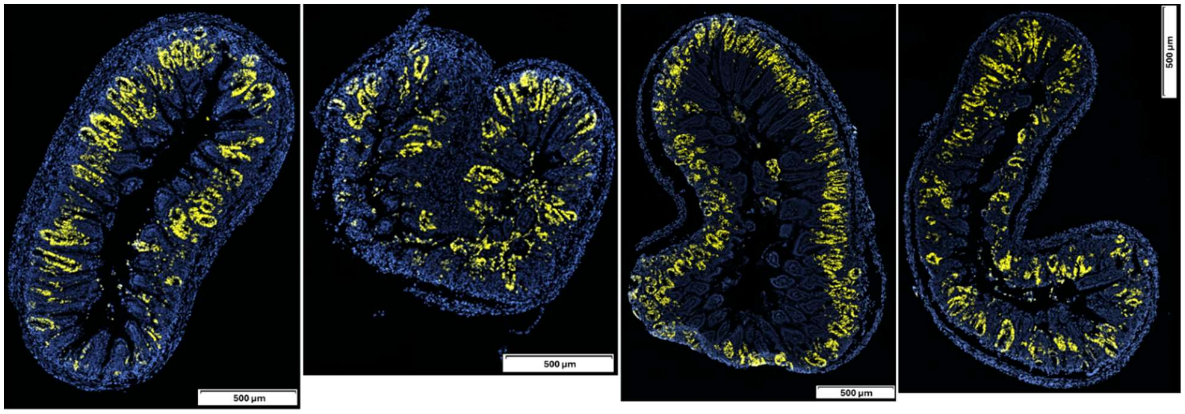


F: 10 Gy + DNA-PKi **2** (10 mg/kg, i.p.)

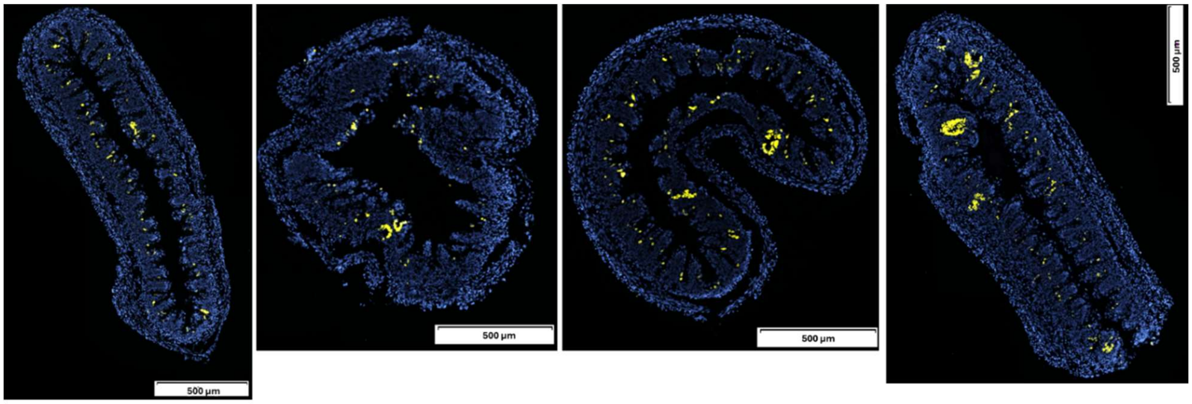

G. 10 Gy + DNA-PKi **2** (10 mg/kg, i.p.)

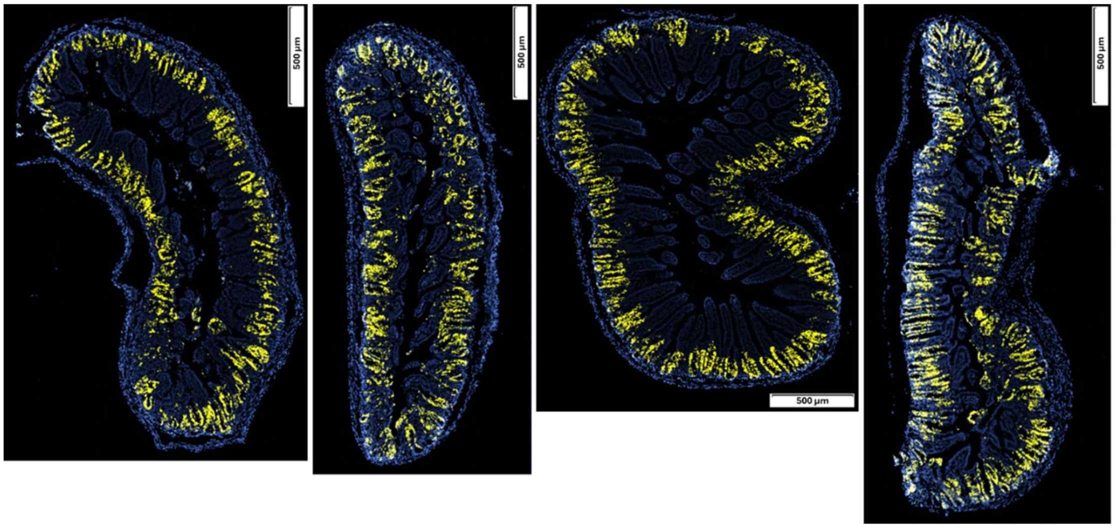

**Fig. S10**. Longitudinal sections of the tongue of CD-1 mice, stained for EdU (gold) and counterstained with H33342 (blue). Methods as in Fig. S10.

A: Untreated controls

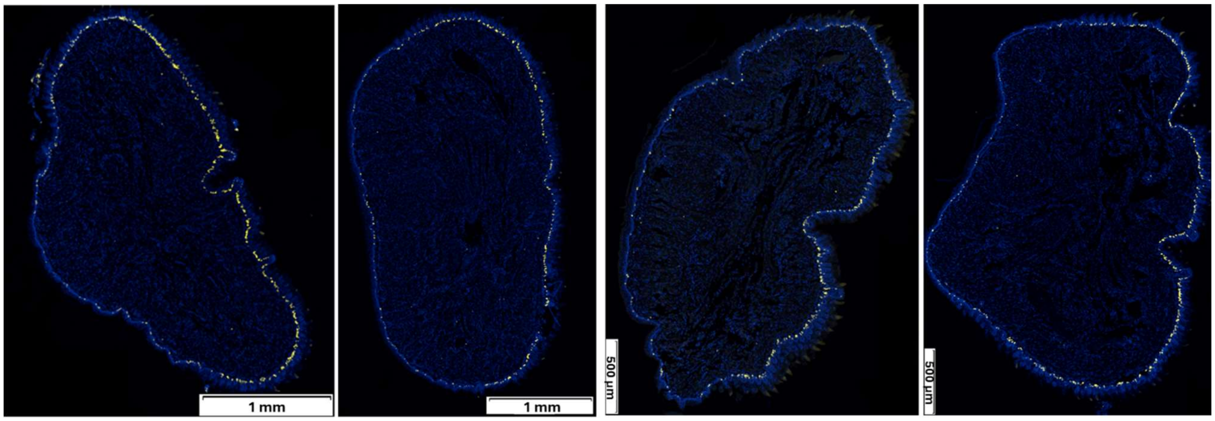


B: DNA-PKi **2** (100 mg/kg, i.p.)

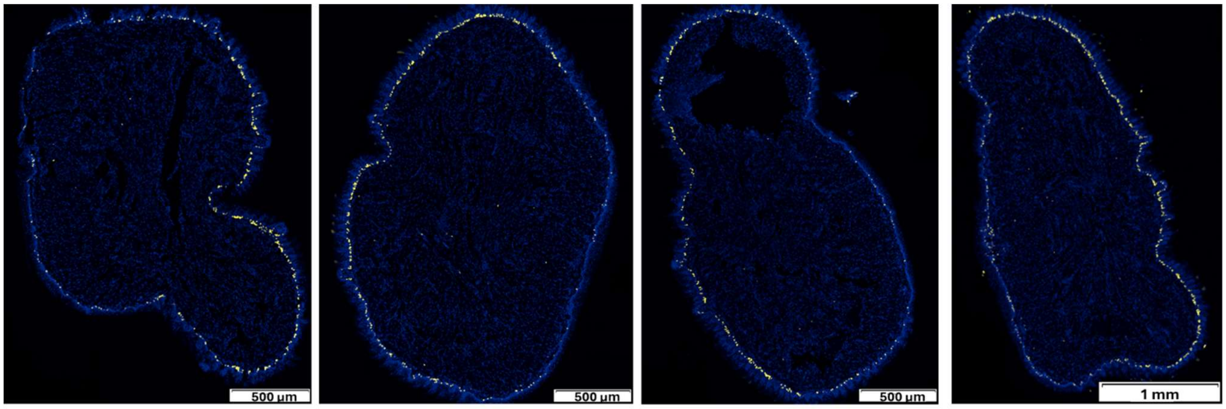


C: Rito (25 mg/kg, p.o) + HAP **4** (400 mg/kg, p.o.)

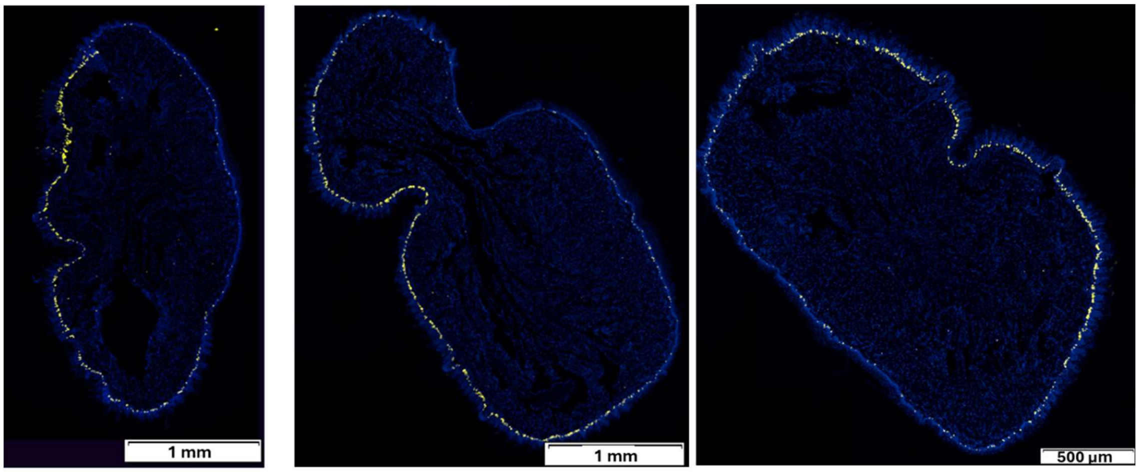

D: 10 Gy

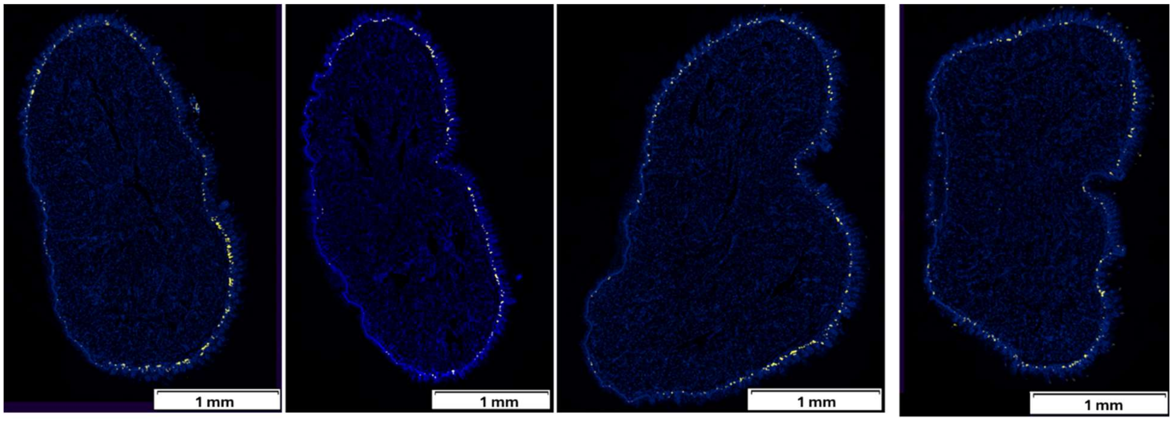


E: 10 Gy + DNA-PKi **2** (10 mg/kg, i.p.)

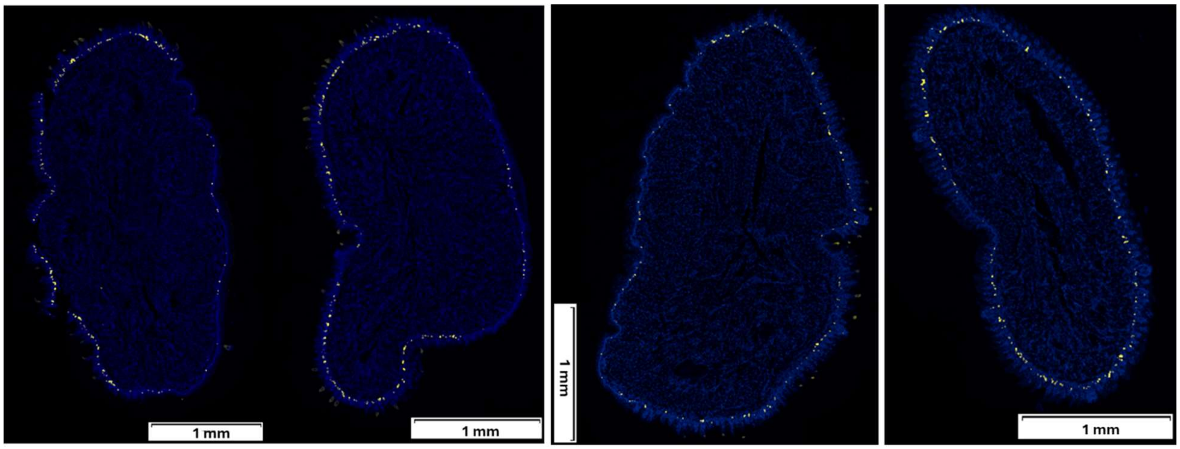


F: 10 Gy + DNA-PKi **2** (100 mg/kg, i.p.)

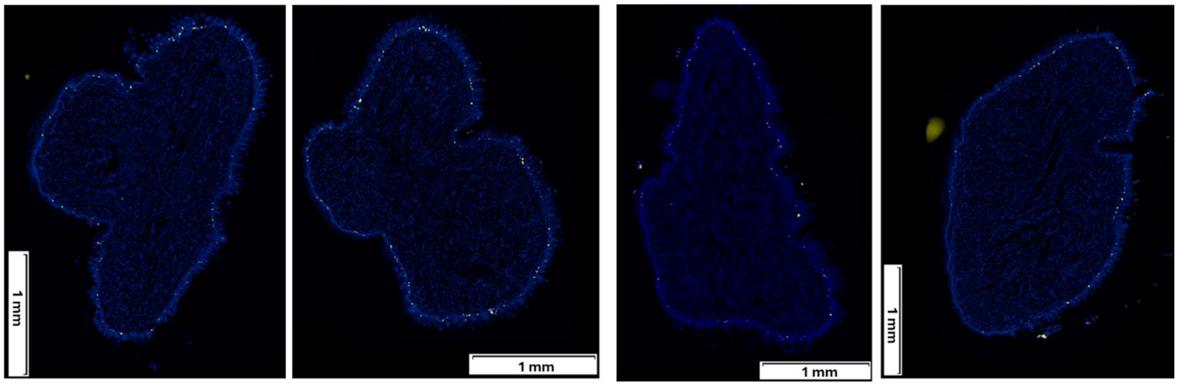

G: 10 Gy + Rito (25 mg/kg, p.o.) + HAP **4** (400 mg/kg, i.p.)

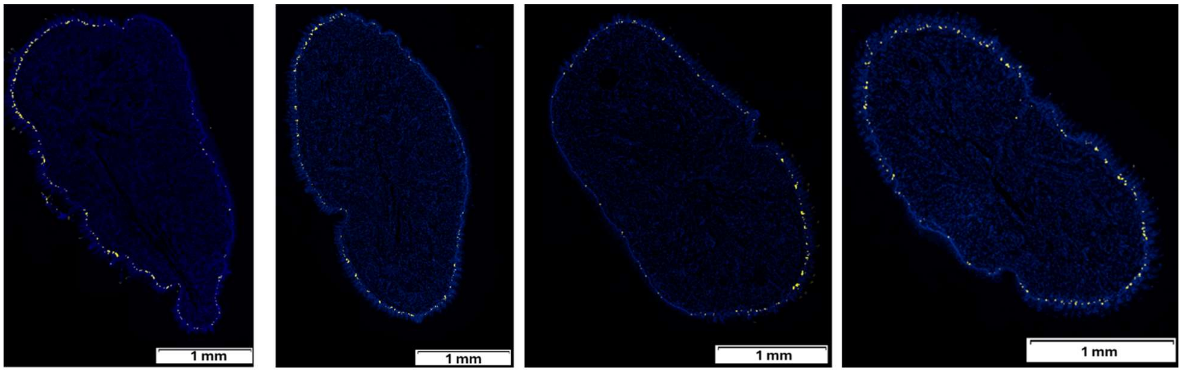

### Supplementary references


1. Hong, C. R. *et al* Identification of 6-anilino imidazo [4,5-*c*] pyridin-2-ones as selective DNA-dependent protein kinase inhibitors and their application as radiosensitizers. *J. Med. Chem.* **67**, 12366–12385 (2024).
2. Liew, L. P. *et al.* Hypoxia-activated prodrugs of PERK inhibitors. *Chem. Asian J.*, **14**, 1238–1248 (2019).
3. Dickson, B. D., Wong, W. W., Wilson, W. R., Hay, M. P. Studies towards hypoxia-activated prodrugs of PARP inhibitors. *Molecules*, **24**, 1559 (2019).
4. Narcombe, P. J., Norris, R. K. Substitution reactions of nitrothiophenes. III. Radical and ionic reactions of 4- and 5-nitro-2-thienyl-methyl and ethyl chlorides and acetates. *Aust. J. Chem.* **32**, 2647-2658 (1979).
5. Hong, C.R. et al. SCCVII tumours and normal tissues in mice by the DNA-dependent protein kinase inhibitor AZD7648. *Radiother. Oncol.* **166**, 162–170 (2022).
6. Jamieson, S.M. *et al*. Evofosfamide for the treatment of human papillomavirus-negative head and neck squamous cell carcinoma. *JCI Insight.* **8**, e169136 (2023).
7. Hong, C.R. *et al.* Bystander effects of hypoxia-activated prodrugs: Agent-based modeling using three dimensional cell cultures. *Front. Pharmacol.* **18**, 10132018 (2018).
8. Jamieson, S. M. *et al.* Evofosfamide for the treatment of human papillomavirus-negative head and neck squamous cell carcinoma. *JCI Insight* **3**, e122204 (2018).
