## Supplementary material for "Systemic CYP3A inhibition by ritonavir enables selective targeting of hypoxic tumour cells by prodrugs of DNA-PK inhibitors": NMR spectra

### $^1$H and $^{13}$C Spectra of Compounds 1-5, 7, 8, 11 and 12

#### Compound 1 $^1$H

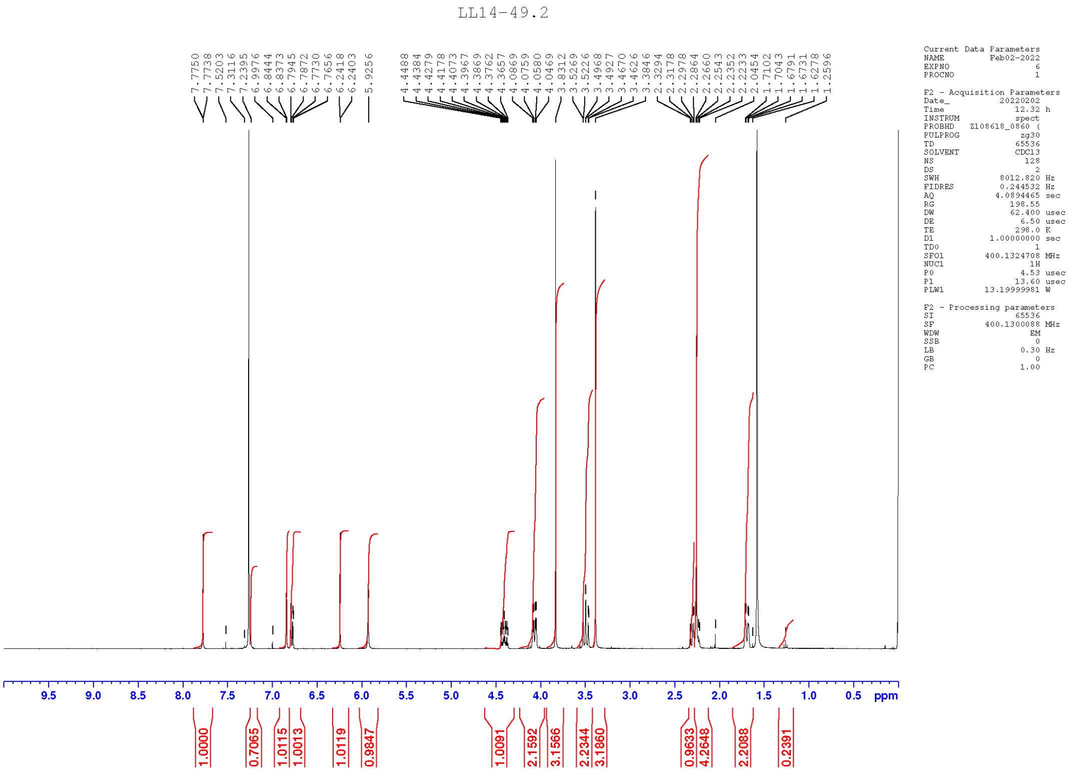


#### Compound 1 $^{13}$C

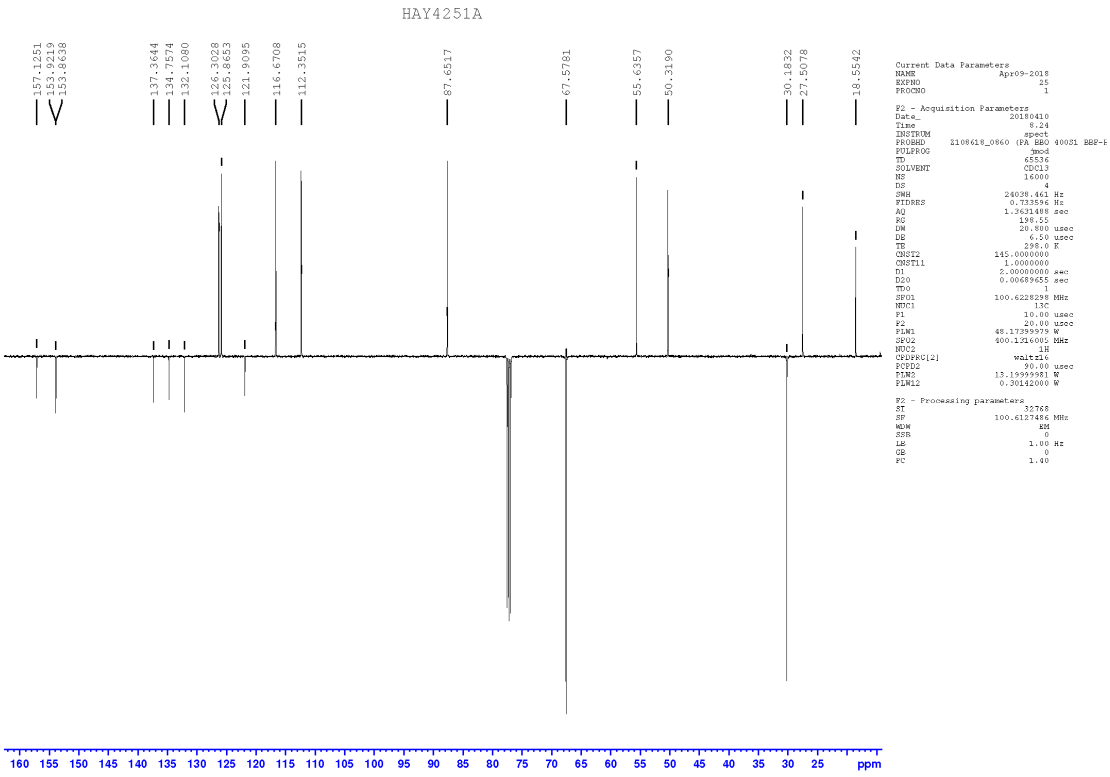

#### Compound 2 $^{1}$H

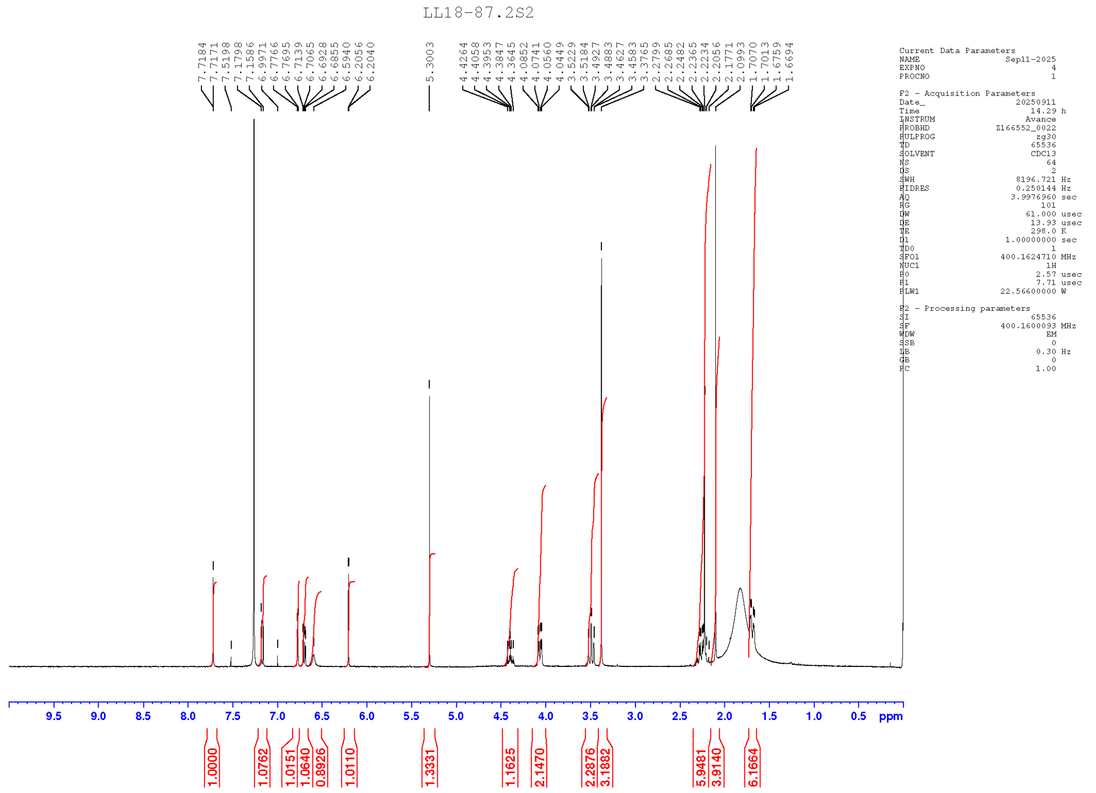

#### Compound 2 $^{13}$C

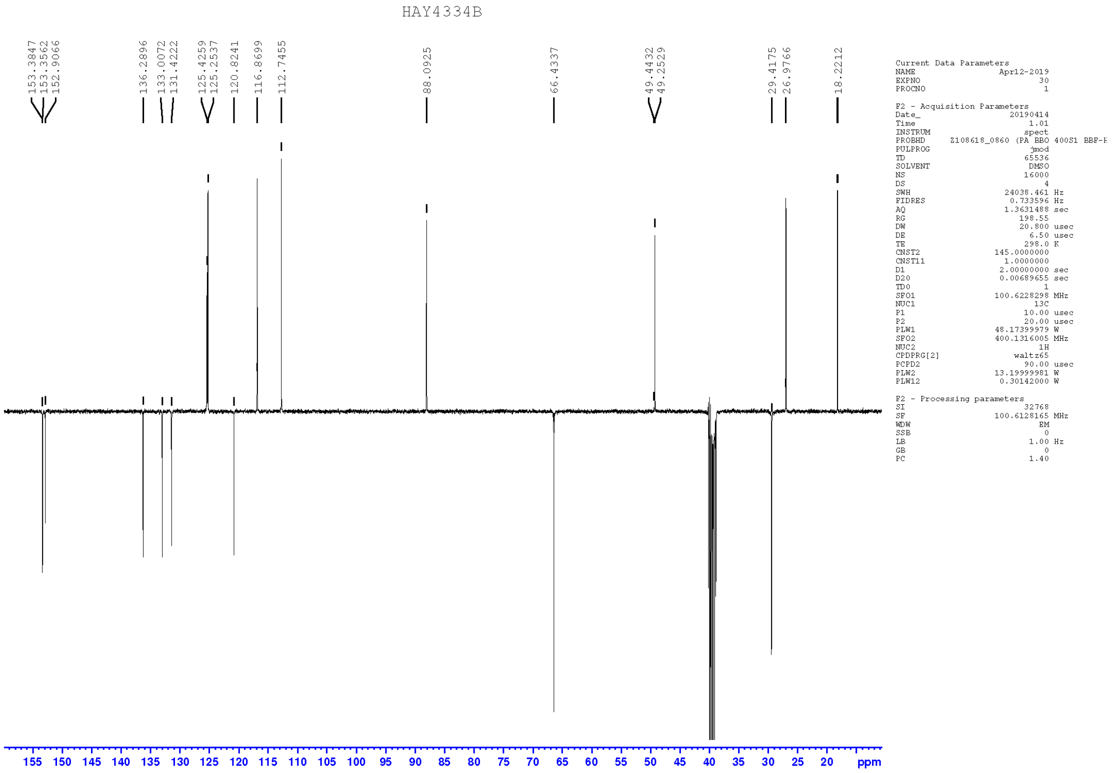

#### Compound 3 ¹H

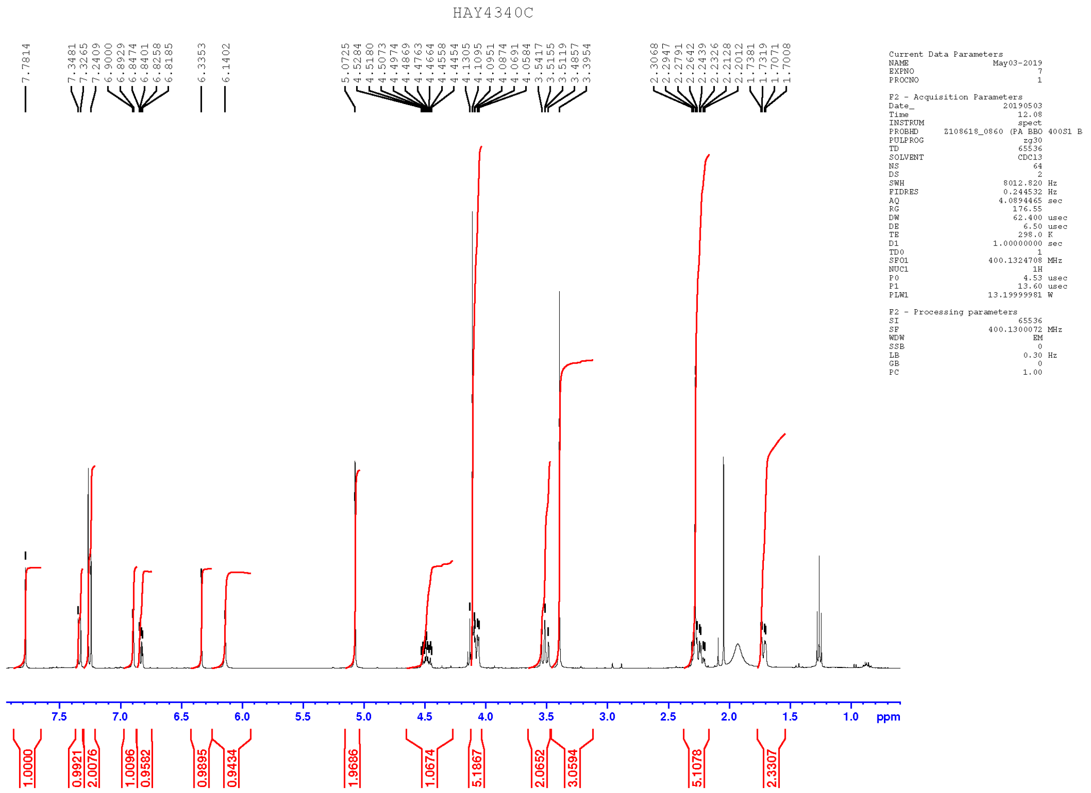


#### Compound 3 ¹³C

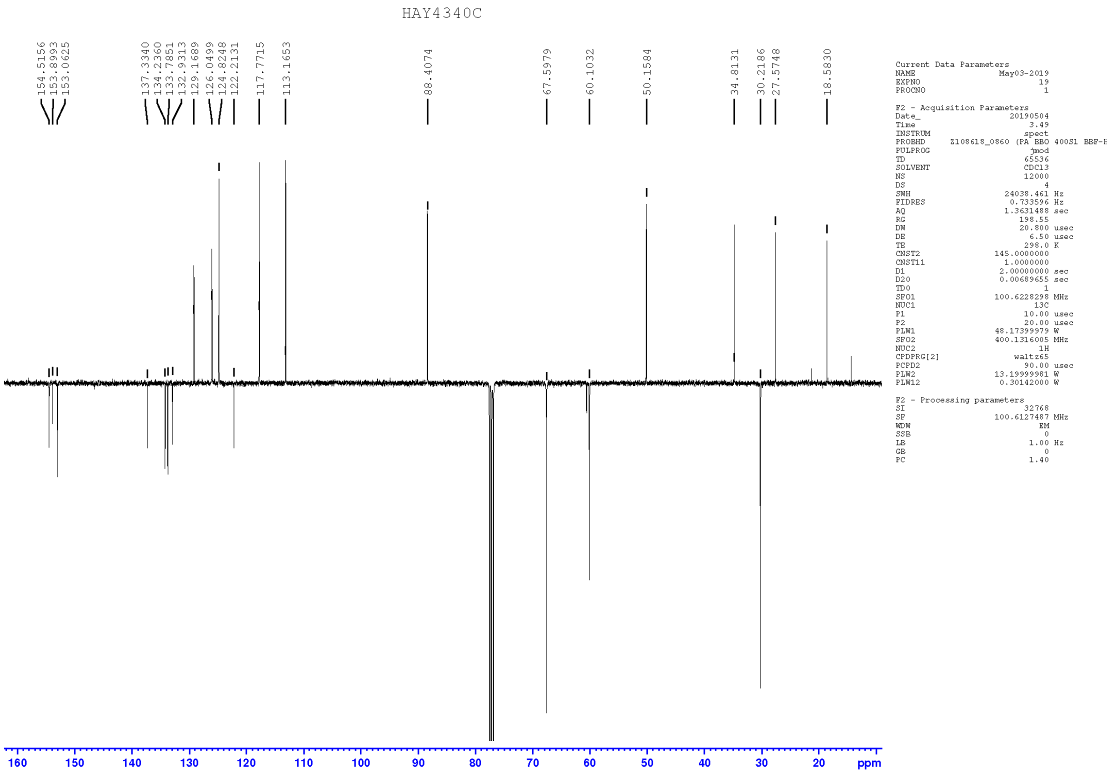

#### Compound 4 $^{1}H$

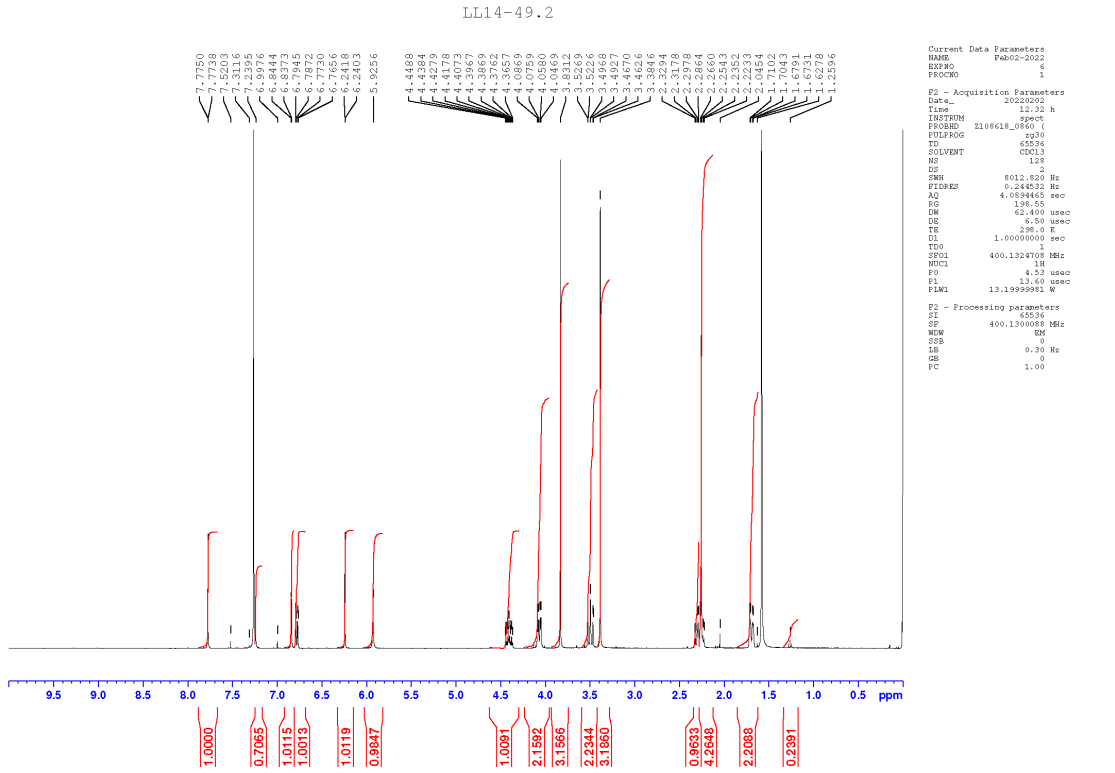

#### Compound 4 $^{13}C$

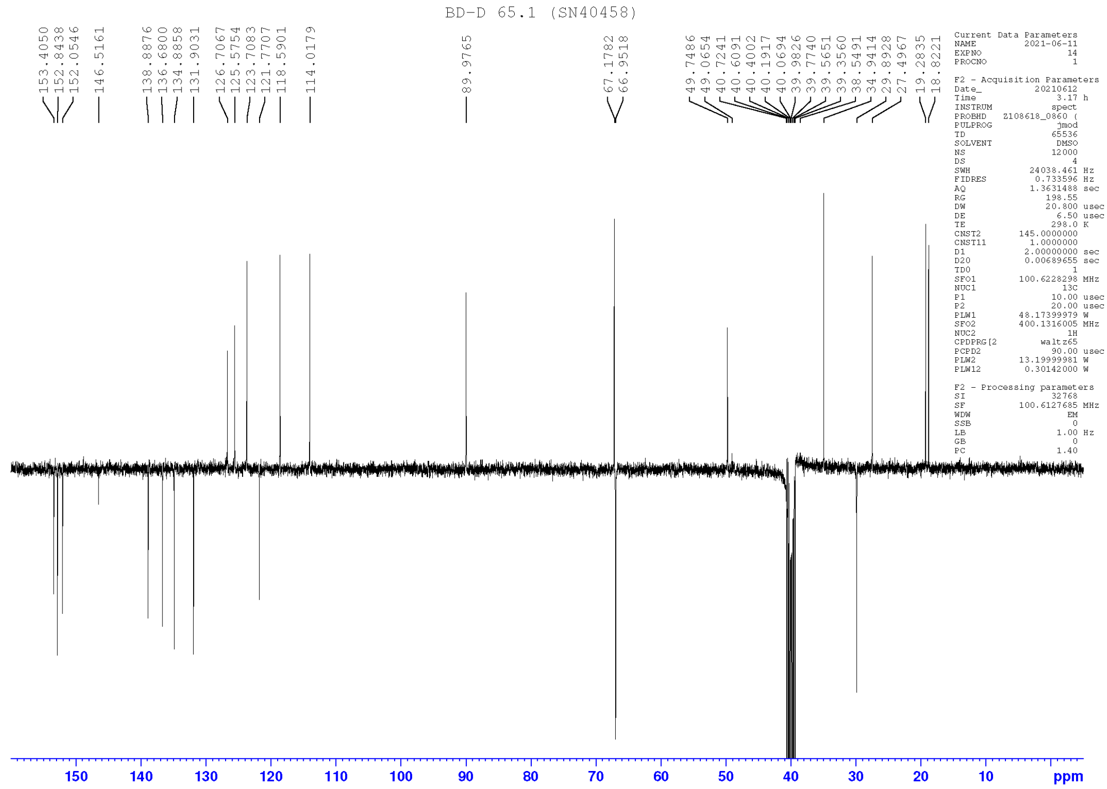

#### Compound 5 $^{1}$H

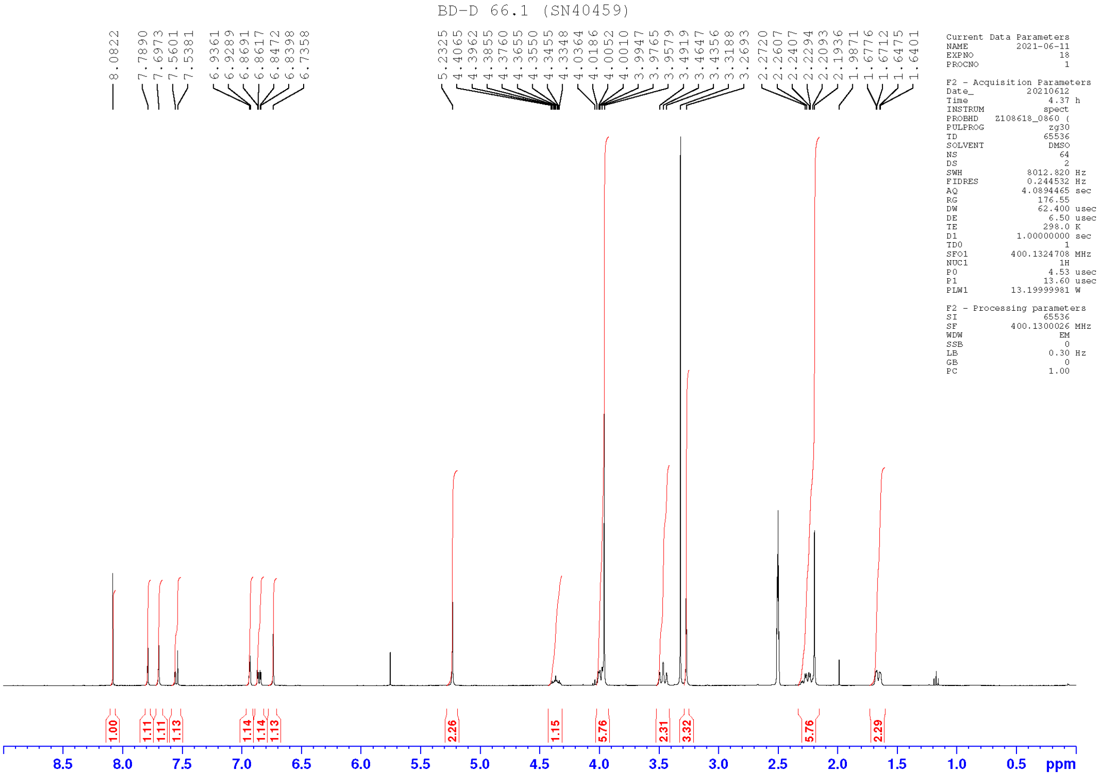


#### Compound 5 $^{13}$C

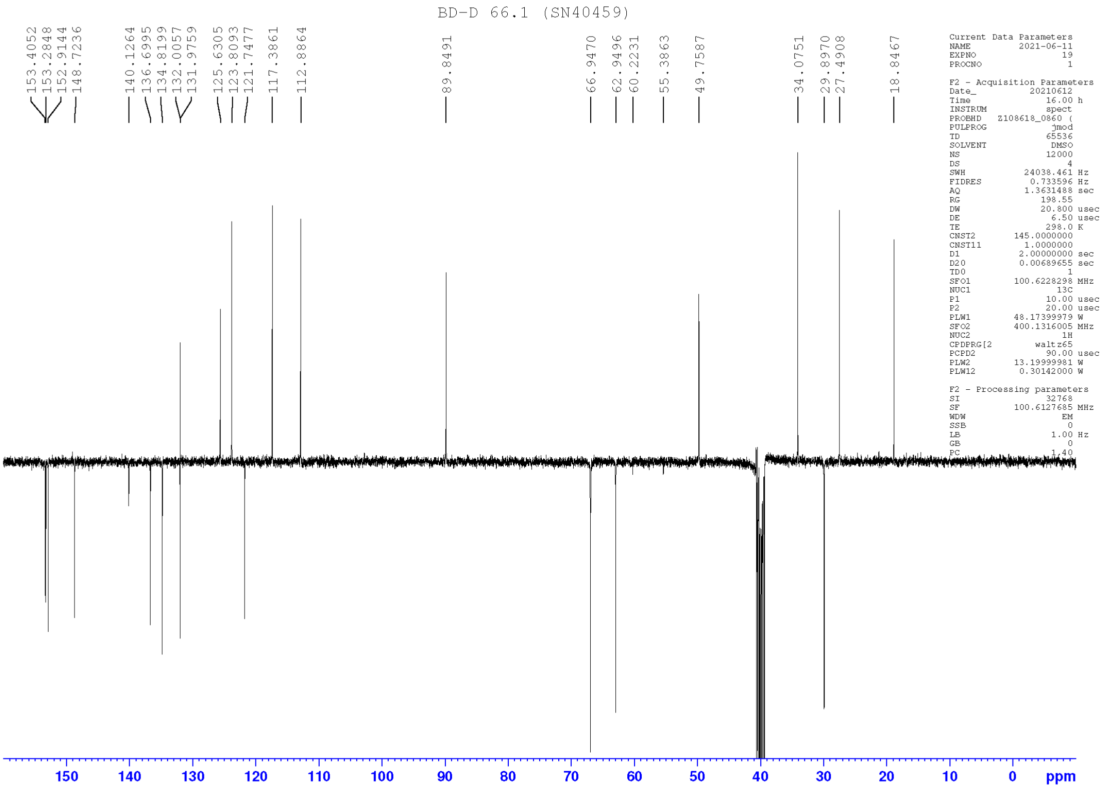

#### Compound 7 $^{1}$H

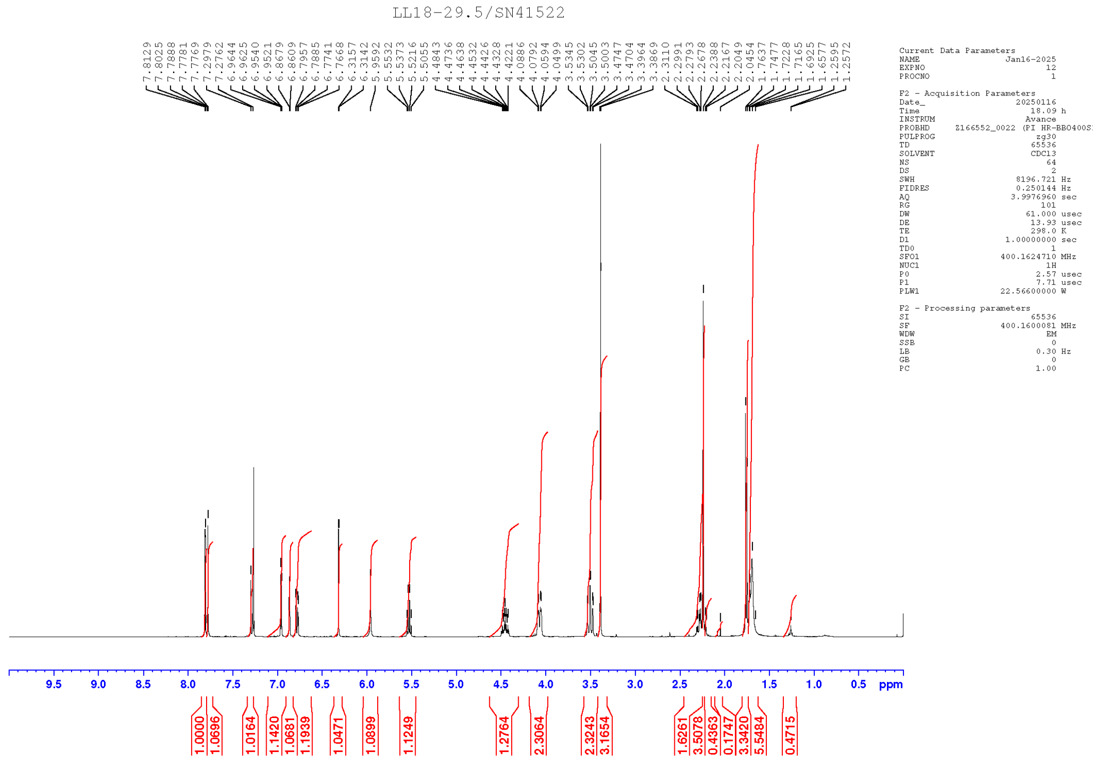


#### Compound 7 $^{13}$C

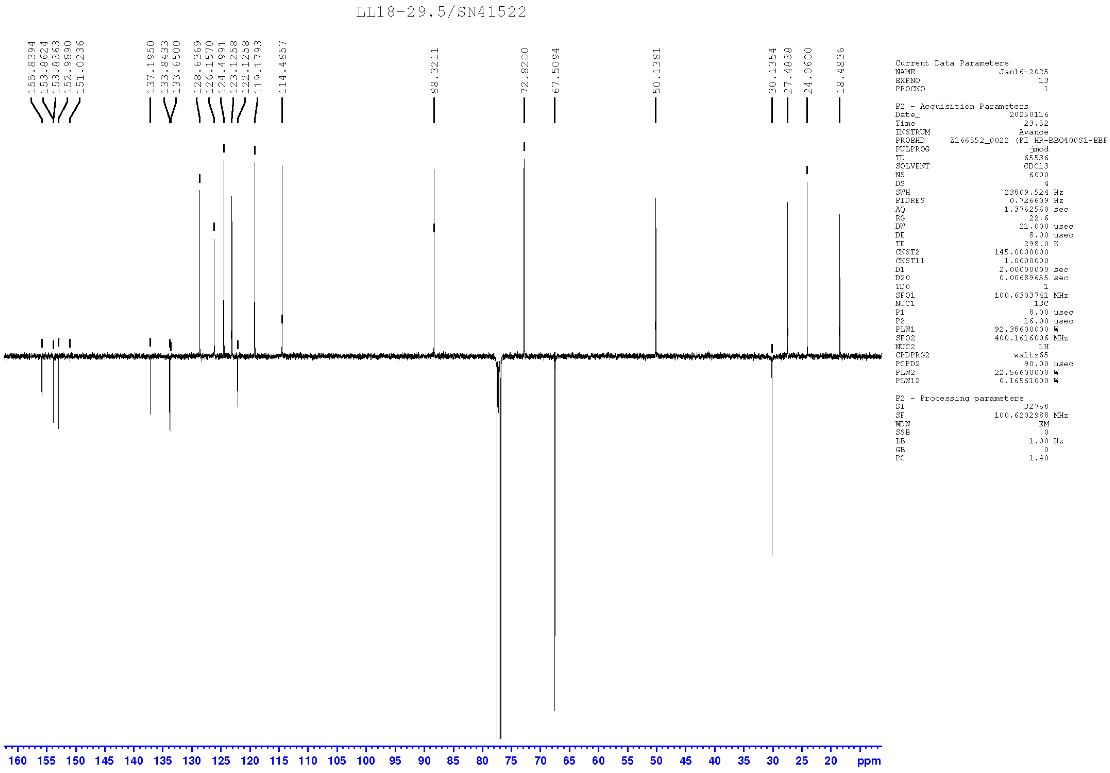

#### Compound 8 $^{1}H$

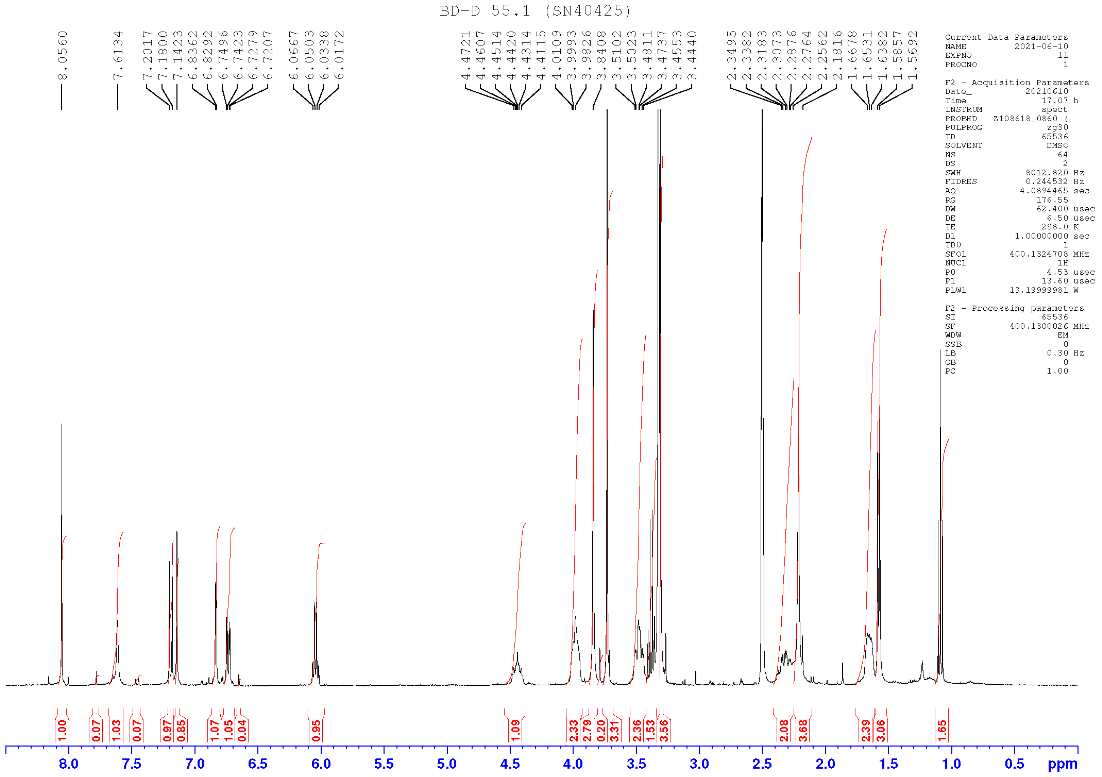

#### Compound 8 $^{13}C$

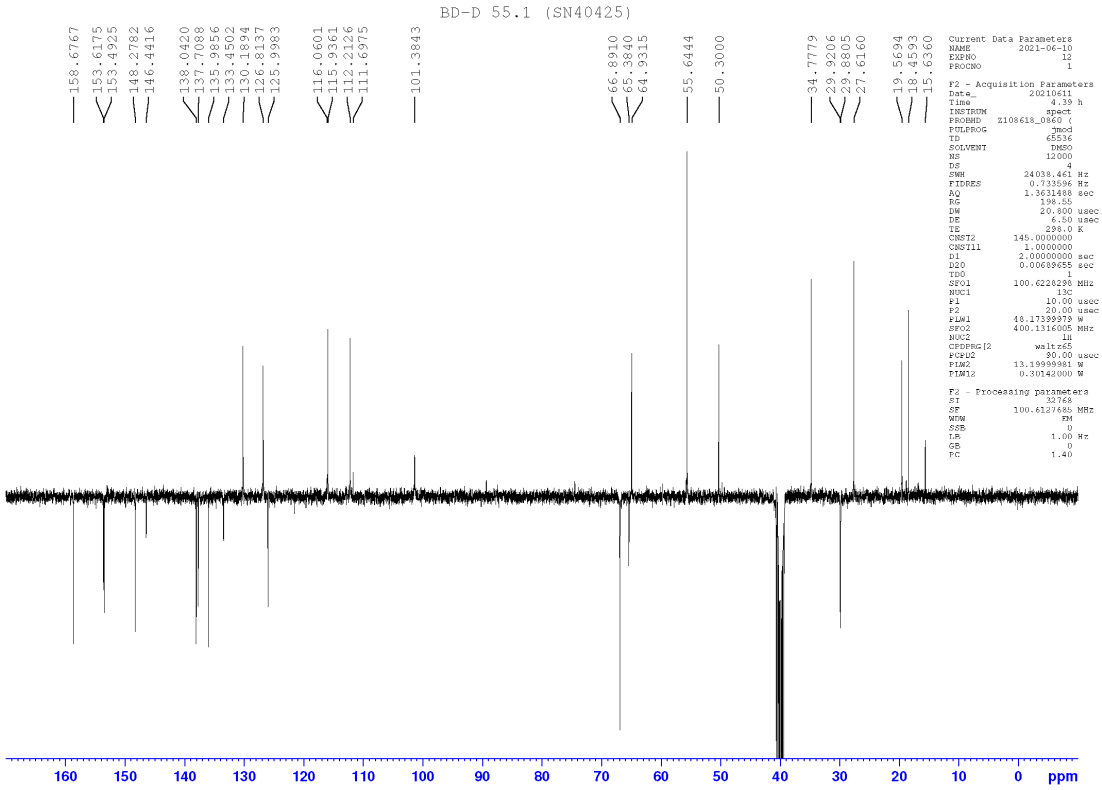

#### Compound 11 $^{1}H$

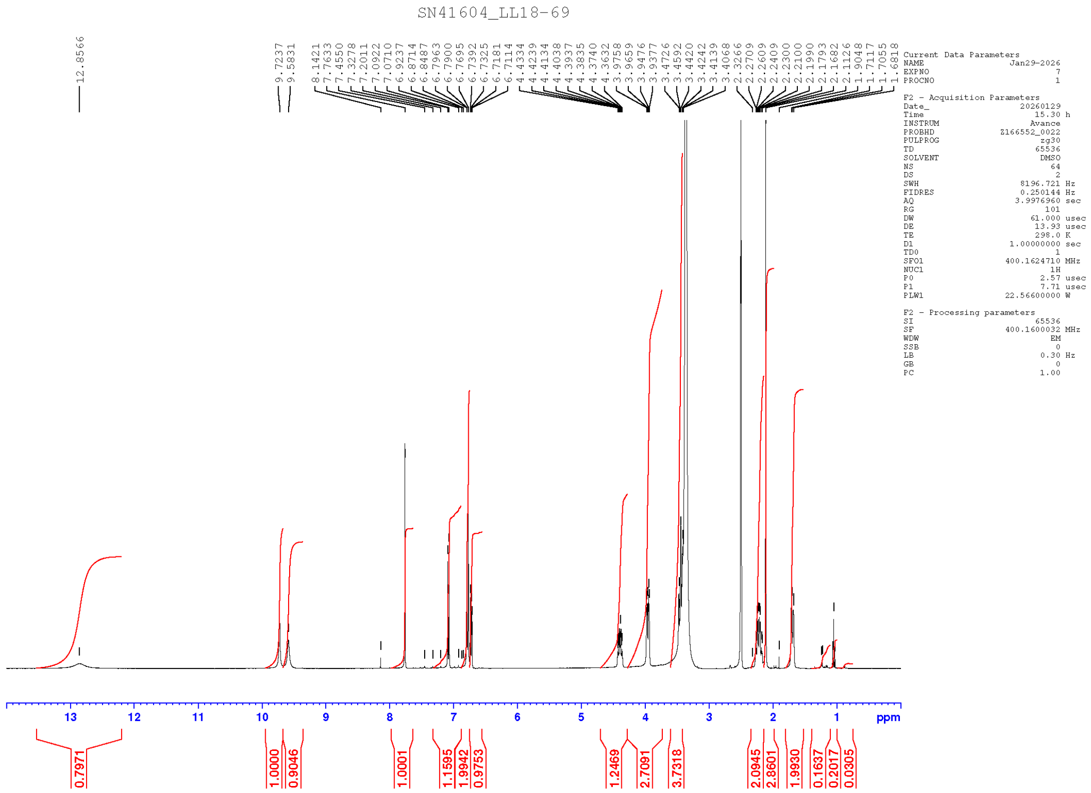


#### Compound 11 $^{13}C$

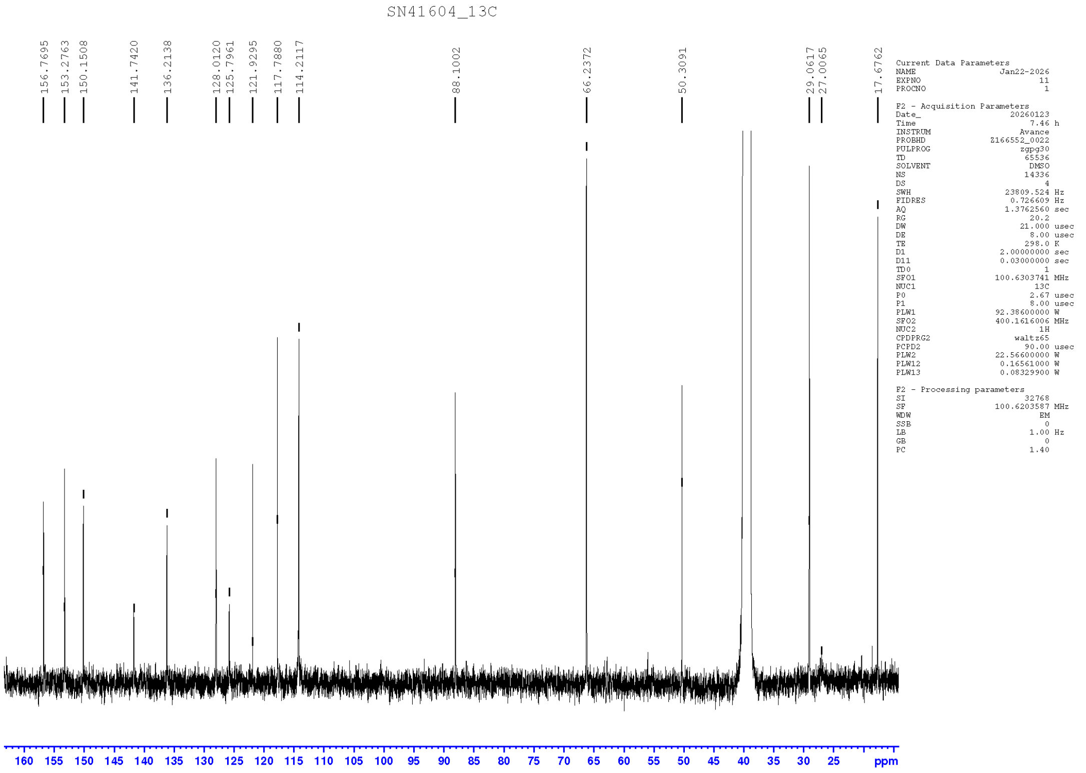

#### Compound 12 $^{1}$H

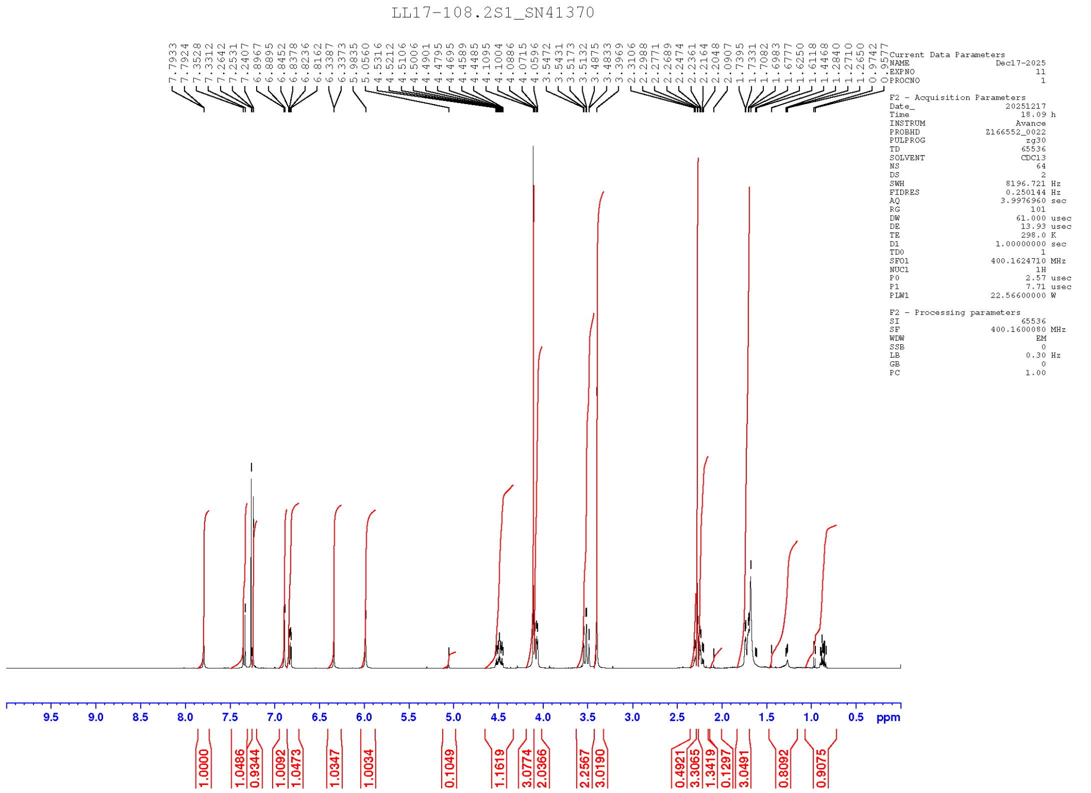

#### Compound 12 $^{13}$C

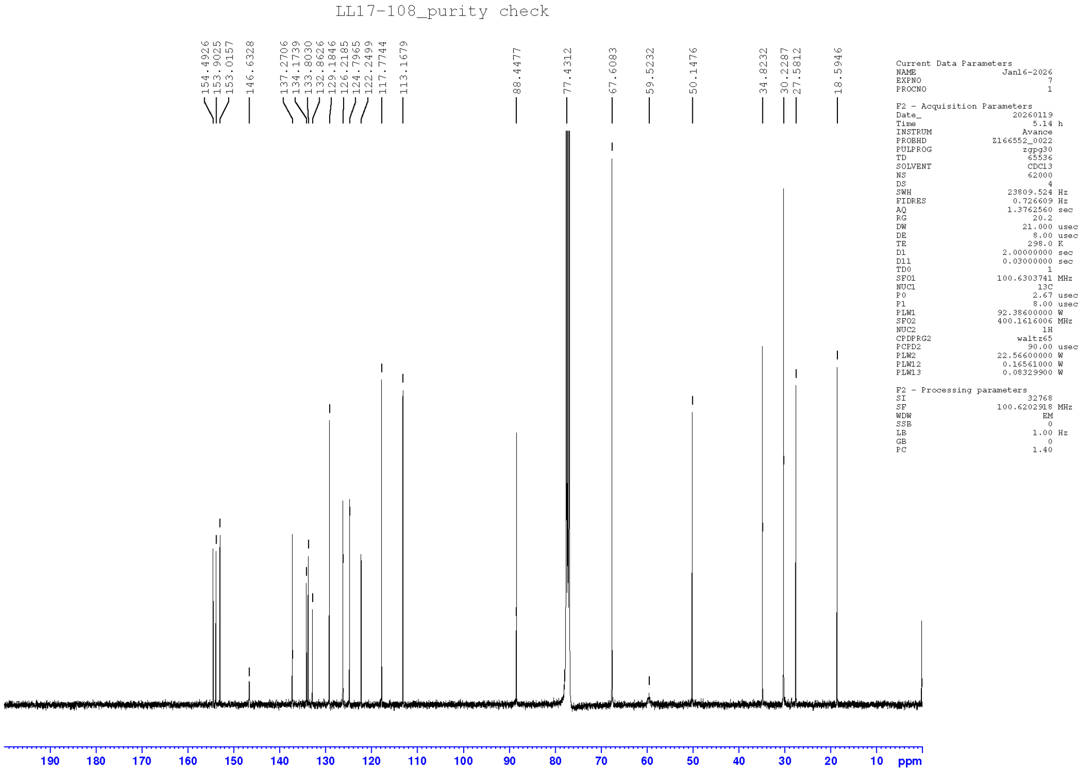

### HPLC Spectra of Compounds 1-5, 7, 8, 11 and 12

#### Compound 1

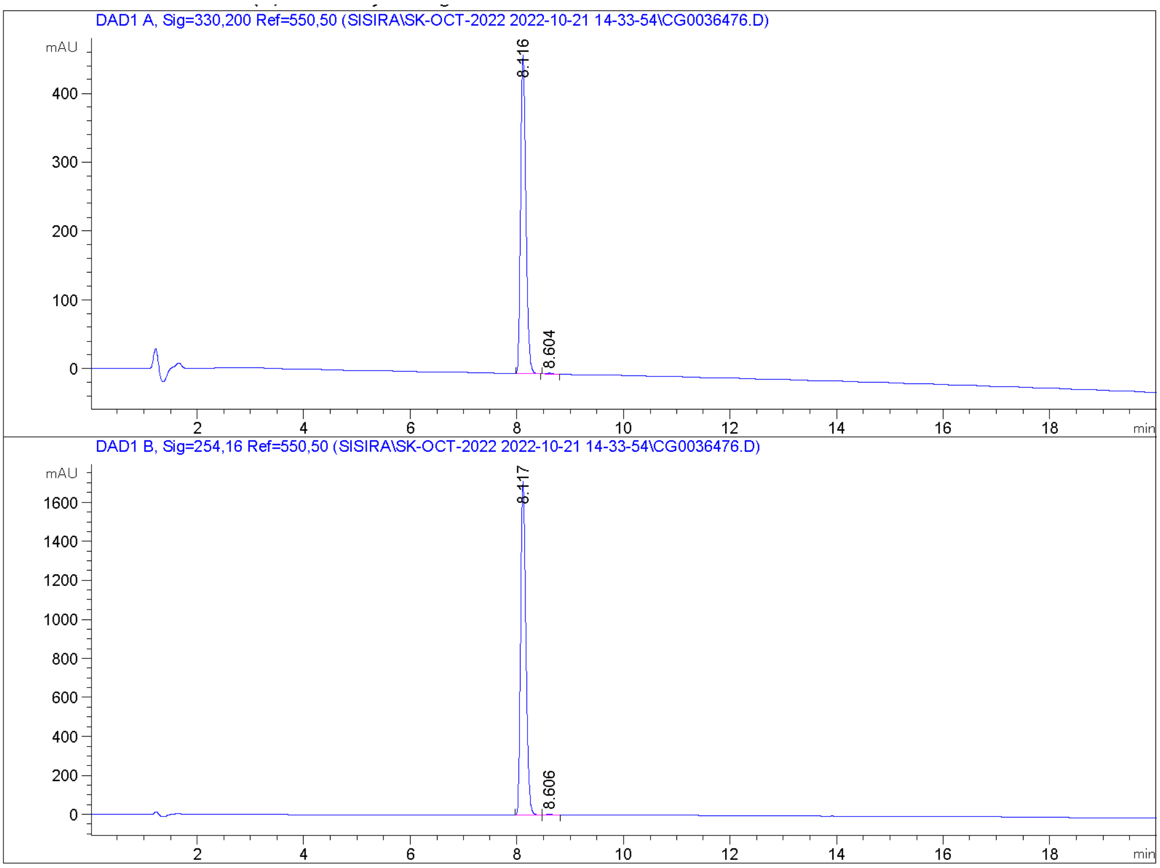


Signal 1: DAD1 A, Sig=330,200 Ref=550,50

| Peak # | RetTime [min] | Type | Width [min] | Area [mAU*s] | Height [mAU] | Area % |
|---|---|---|---|---|---|---|
| 1 | 8.116 | BB | 0.1084 | 3193.94995 | 463.59213 | 99.6238 |
| 2 | 8.604 | BB | 0.1148 | 12.06129 | 1.54937 | 0.3762 |

Totals : 3206.01124 465.14151

**Compound 2**

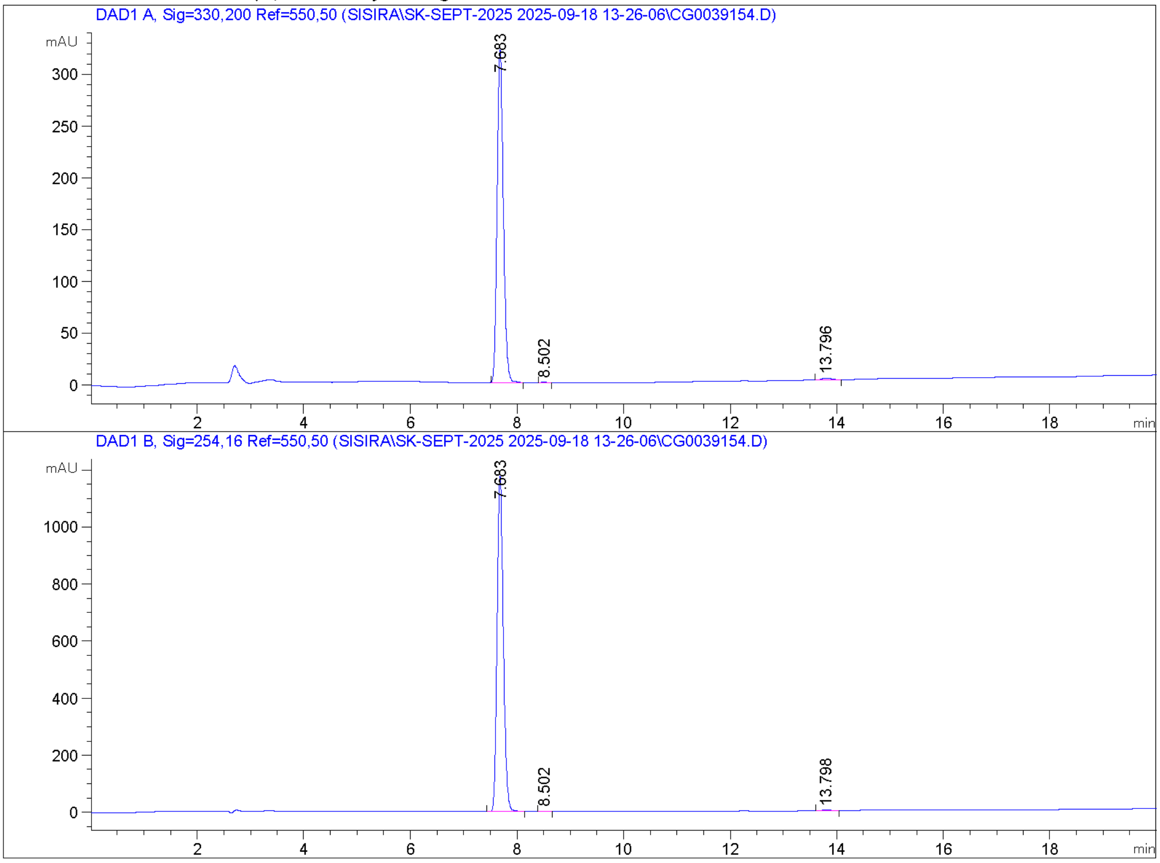


Signal 1: DAD1 A, Sig=330,200 Ref=550,50

| Peak # | RetTime [min] | Type | Width [min] | Area [mAU*s] | Height [mAU] | Area % |
|---|---|---|---|---|---|---|
| 1 | 7.683 | BB | 0.1267 | 2625.34424 | 323.56531 | 99.0544 |
| 2 | 8.502 | MM | 0.1134 | 3.75273 | 5.51655e-1 | 0.1416 |
| 3 | 13.796 | BB | 0.1829 | 21.30984 | 1.73940 | 0.8040 |

Totals : 2650.40681 325.85636

#### Compound 3

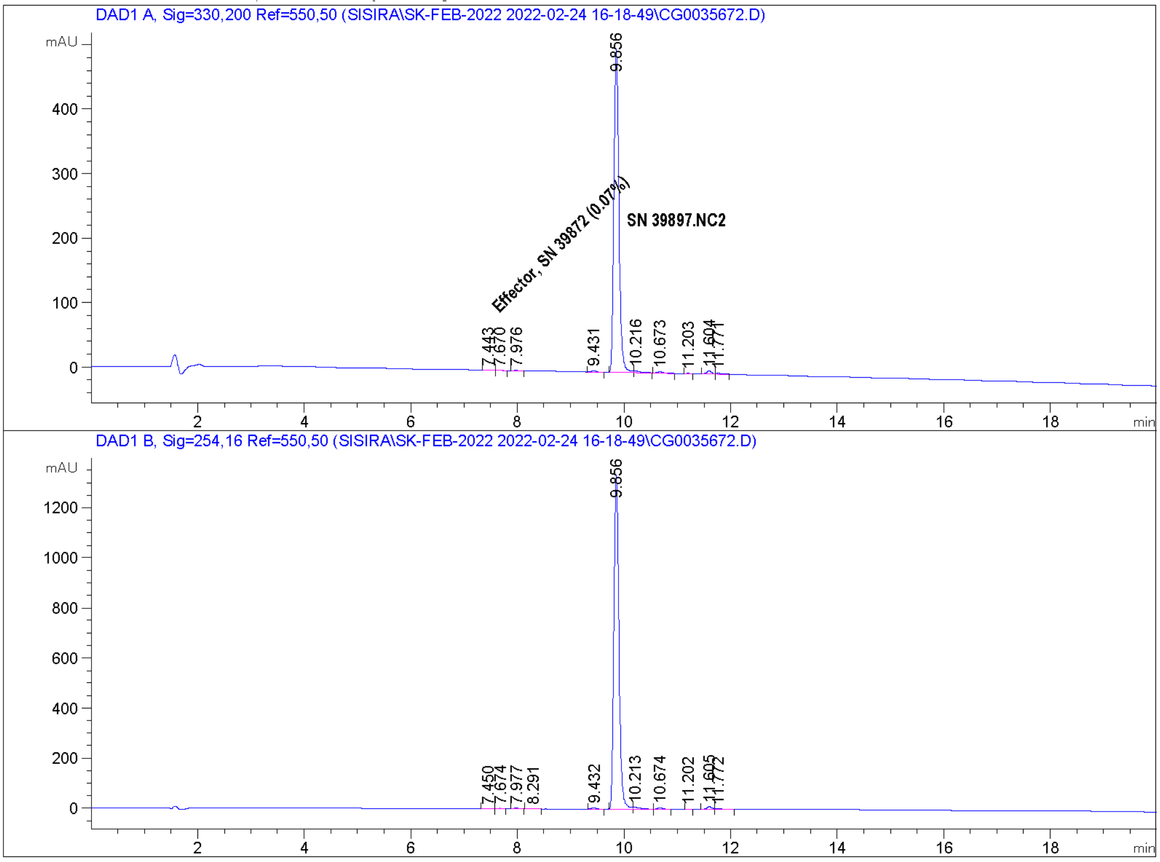


Signal 1: DAD1 A, Sig=330,200 Ref=550,50

| Peak # | RetTime [min] | Type | Width [min] | Area [mAU*s] | Height [mAU] | Area % |
|---|---|---|---|---|---|---|
| 1 | 7.443 | MF | 0.1108 | 2.98949 | 4.49480e-1 | 0.0885 |
| 2 | 7.670 | FM | 0.0890 | 2.30042 | 4.30653e-1 | 0.0681 |
| 3 | 7.976 | MM | 0.0887 | 3.43339 | 6.45138e-1 | 0.1017 |
| 4 | 9.431 | BB | 0.0989 | 10.85664 | 1.78714 | 0.3215 |
| 5 | 9.856 | MF | 0.1086 | 3277.57983 | 503.05859 | 97.0564 |
| 6 | 10.216 | FM | 0.1306 | 17.63085 | 2.25061 | 0.5221 |
| 7 | 10.673 | BB | 0.1021 | 14.33208 | 2.14191 | 0.4244 |
| 8 | 11.203 | MM | 0.0826 | 1.93203 | 3.89989e-1 | 0.0572 |
| 9 | 11.604 | BV | 0.1124 | 34.40152 | 4.75498 | 1.0187 |
| 10 | 11.771 | VB | 0.1067 | 11.52854 | 1.55086 | 0.3414 |

Totals : 3376.98480 517.45935

#### Compound 4

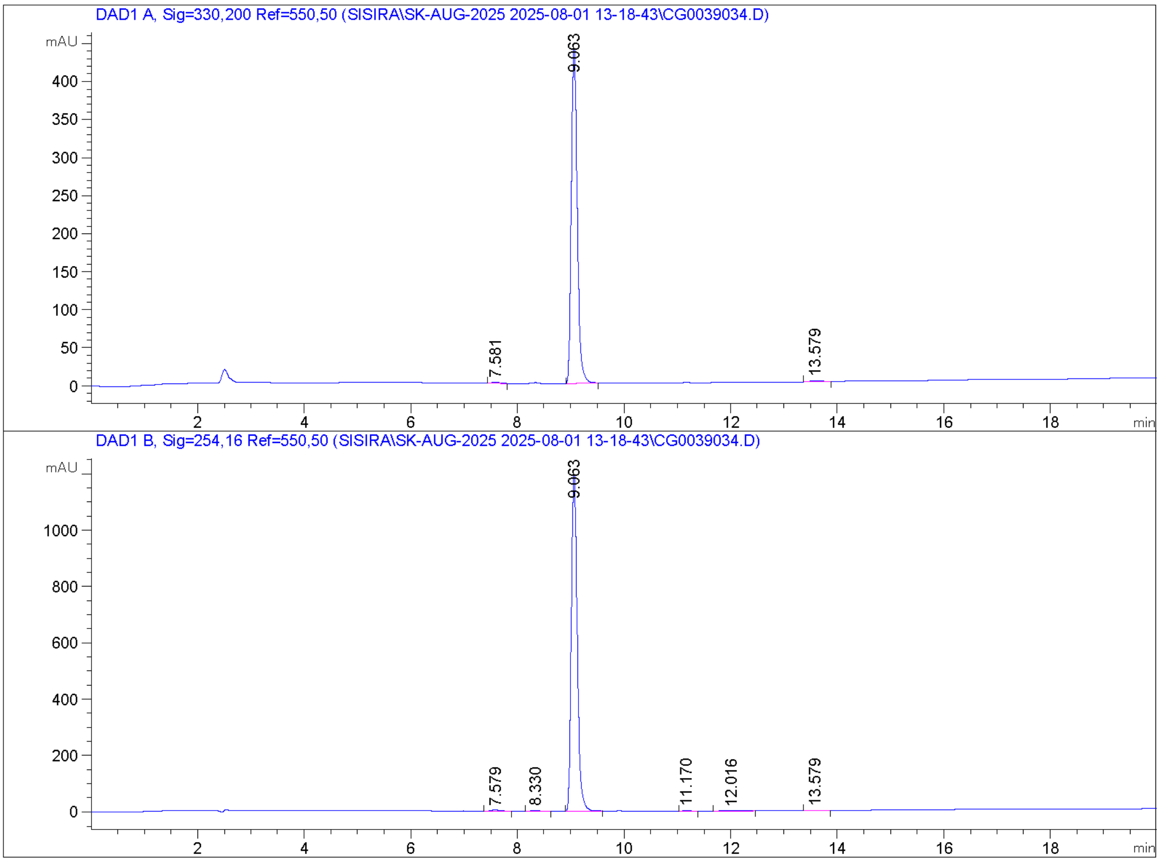


Signal 1: DAD1 A, Sig=330,200 Ref=550,50

| Peak # | RetTime [min] | Type | Width [min] | Area [mAU*s] | Height [mAU] | Area % |
|---|---|---|---|---|---|---|
| 1 | 7.581 | BB | 0.1410 | 14.24465 | 1.52531 | 0.4167 |
| 2 | 9.063 | BB | 0.1214 | 3386.42041 | 441.84253 | 99.0571 |
| 3 | 13.579 | BB | 0.1804 | 17.98889 | 1.49521 | 0.5262 |

Totals : 3418.65395 444.86304

#### Compound 5

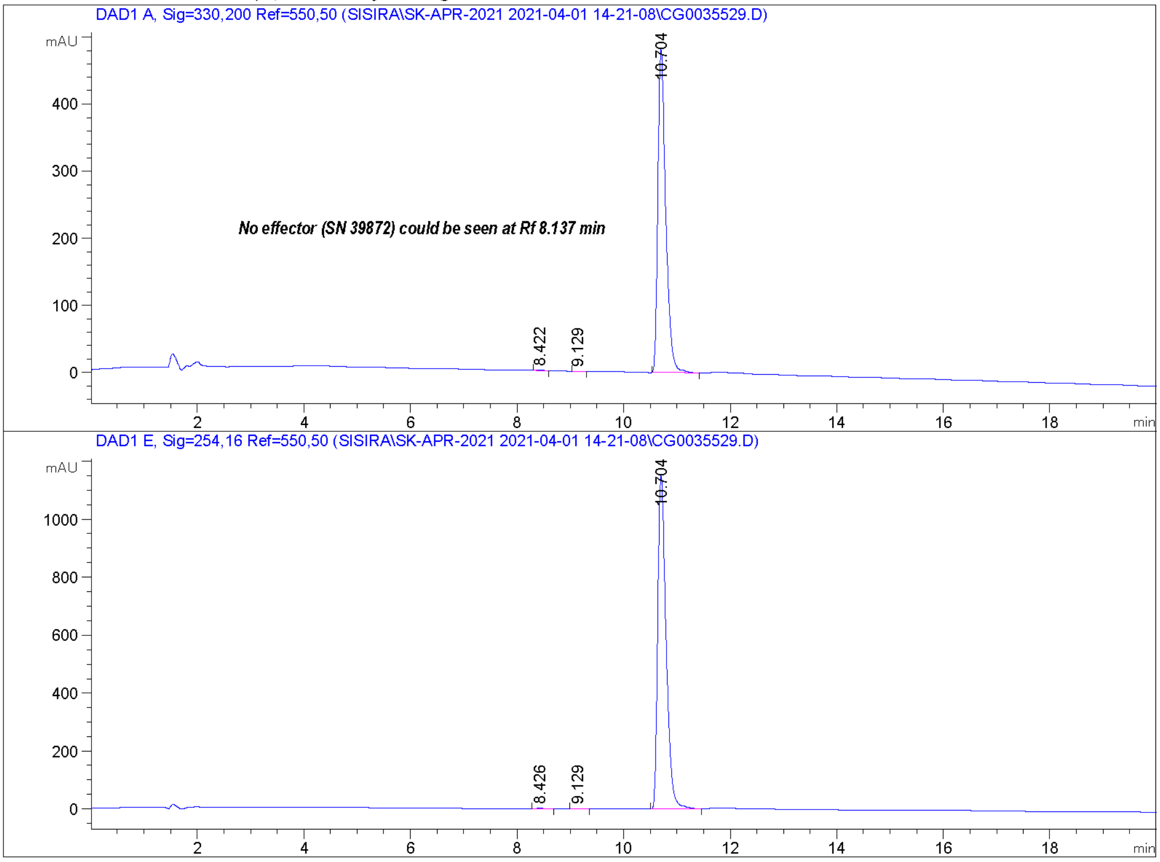


Signal 1: DAD1 A, Sig=330,200 Ref=550,50

| Peak # | RetTime [min] | Type | Width [min] | Area [mAU*s] | Height [mAU] | Area % |
|---|---|---|---|---|---|---|
| 1 | 8.422 | MM | 0.1257 | 4.29073 | 5.69004e-1 | 0.0876 |
| 2 | 9.129 | MM | 0.1285 | 1.94863 | 2.52815e-1 | 0.0398 |
| 3 | 10.704 | MM | 0.1688 | 4894.20557 | 483.16489 | 99.8727 |

Totals : 4900.44492 483.98671

#### Compound 7

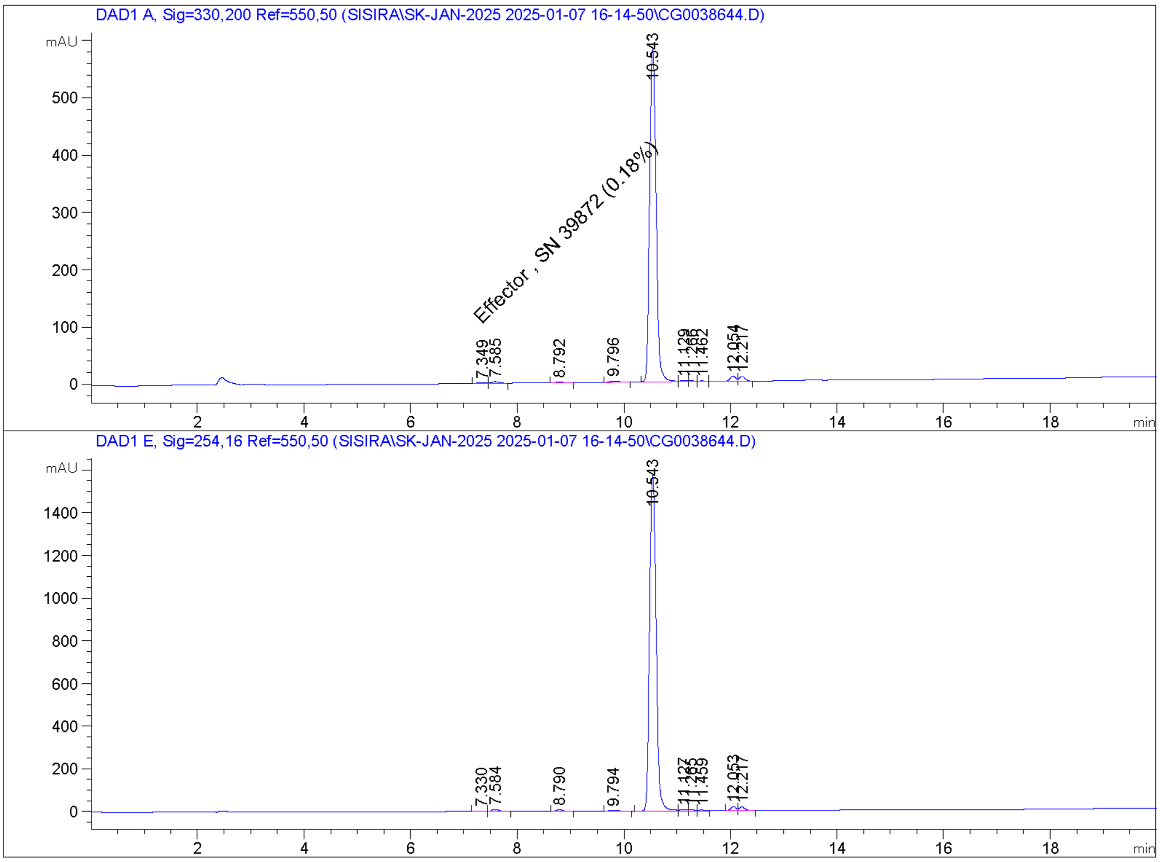


Signal 1: DAD1 A, Sig=330,200 Ref=550,50

| Peak # | RetTime [min] | Type | Width [min] | Area [mAU*s] | Height [mAU] | Area % |
|---|---|---|---|---|---|---|
| 1 | 7.349 | MF | 0.1852 | 9.45488 | 8.50895e-1 | 0.1841 |
| 2 | 7.585 | FM | 0.1611 | 27.02843 | 2.79566 | 0.5264 |
| 3 | 8.792 | BB | 0.1506 | 18.04240 | 1.90375 | 0.3514 |
| 4 | 9.796 | BB | 0.1740 | 23.99941 | 1.97178 | 0.4674 |
| 5 | 10.543 | BV | 0.1340 | 4884.36865 | 581.68329 | 95.1223 |
| 6 | 11.129 | VV | 0.1222 | 18.61271 | 2.21188 | 0.3625 |
| 7 | 11.266 | VV | 0.1091 | 13.88770 | 1.81868 | 0.2705 |
| 8 | 11.462 | VV | 0.1312 | 10.52611 | 1.19028 | 0.2050 |
| 9 | 12.054 | VV | 0.1189 | 70.27857 | 9.02123 | 1.3687 |
| 10 | 12.217 | VB | 0.1126 | 58.63079 | 8.08092 | 1.1418 |

Totals : 5134.82965 611.52836

#### Compound 8

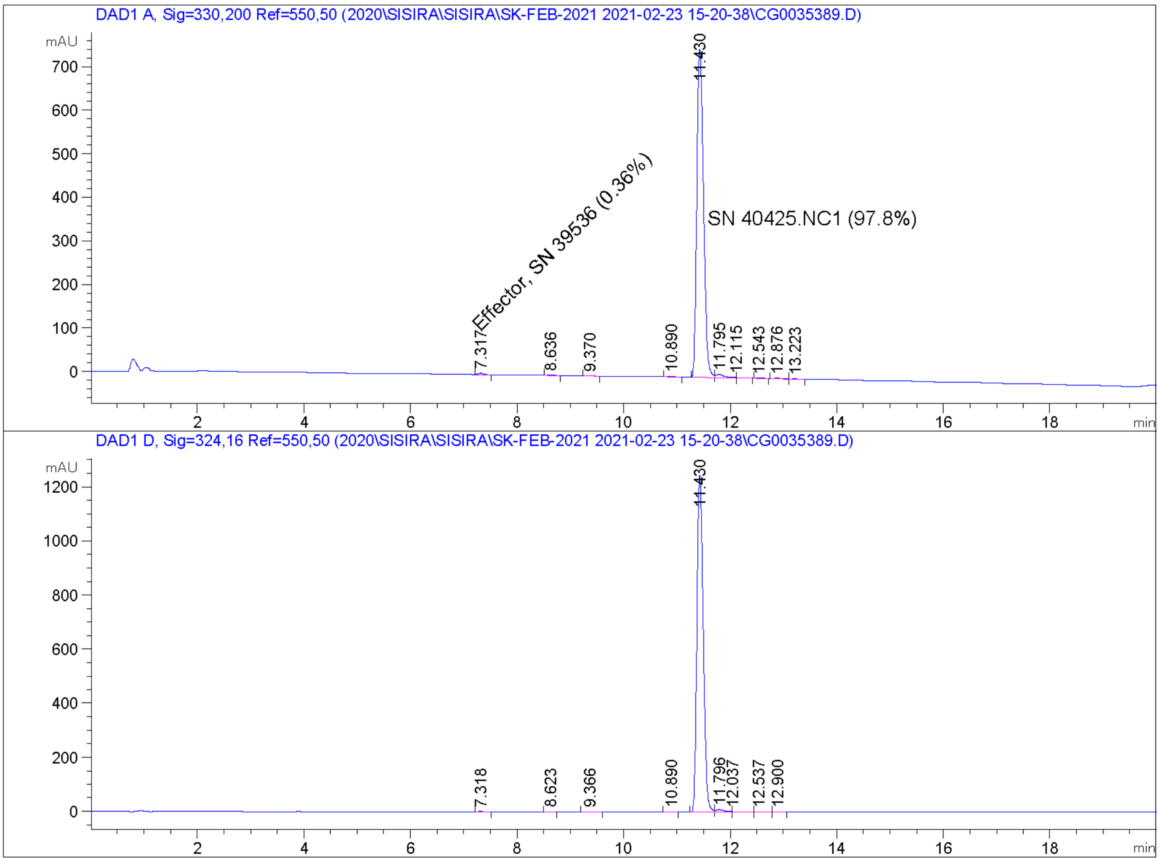


Signal 1: DAD1 A, Sig=330,200 Ref=550,50

| Peak # | RetTime [min] | Type | Width [min] | Area [mAU*s] | Height [mAU] | Area % |
|---|---|---|---|---|---|---|
| 1 | 7.317 | BB | 0.1070 | 24.46666 | 3.61369 | 0.3637 |
| 2 | 8.636 | MM | 0.1372 | 3.25501 | 3.95378e-1 | 0.0484 |
| 3 | 9.370 | MM | 0.1194 | 5.94459 | 8.29740e-1 | 0.0884 |
| 4 | 10.890 | BB | 0.1130 | 9.52399 | 1.30701 | 0.1416 |
| 5 | 11.430 | BV | 0.1373 | 6576.83594 | 757.48364 | 97.7533 |
| 6 | 11.795 | MF | 0.1809 | 84.38053 | 7.77438 | 1.2542 |
| 7 | 12.115 | FM | 0.1390 | 8.86875 | 1.06326 | 0.1318 |
| 8 | 12.543 | MM | 0.1163 | 2.28139 | 3.27031e-1 | 0.0339 |
| 9 | 12.876 | MF | 0.1349 | 8.30752 | 1.02652 | 0.1235 |
| 10 | 13.223 | FM | 0.1326 | 4.13277 | 5.19426e-1 | 0.0614 |

Totals : 6727.99716 774.34008

#### Compound 11

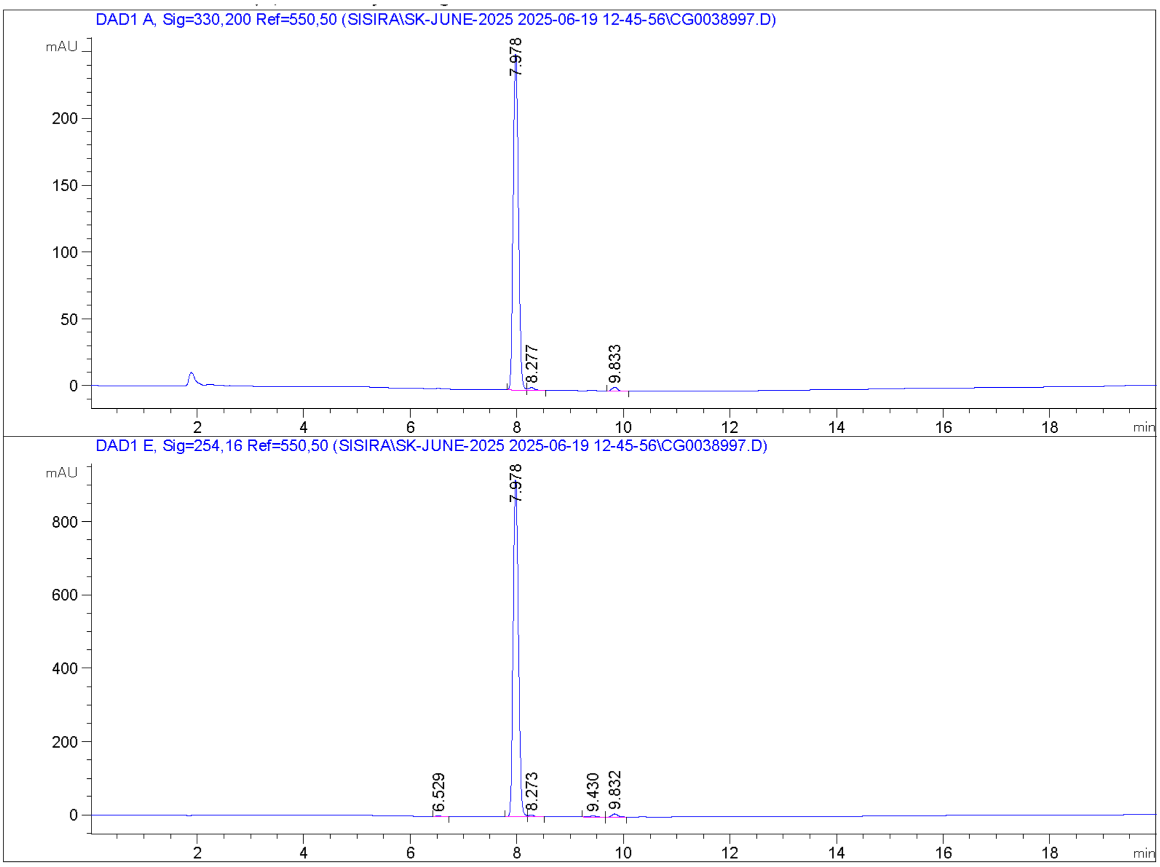


Signal 1: DAD1 A, Sig=330,200 Ref=550,50

| Peak # | RetTime [min] | Type | Width [min] | Area [mAU*s] | Height [mAU] | Area % |
|---|---|---|---|---|---|---|
| 1 | 7.978 | BV | 0.1058 | 1689.92749 | 253.57448 | 97.9001 |
| 2 | 8.277 | VB | 0.1108 | 15.21731 | 2.04423 | 0.8816 |
| 3 | 9.833 | BB | 0.1191 | 21.03091 | 2.81544 | 1.2184 |

Totals : 1726.17571 258.43414

#### Compound 12

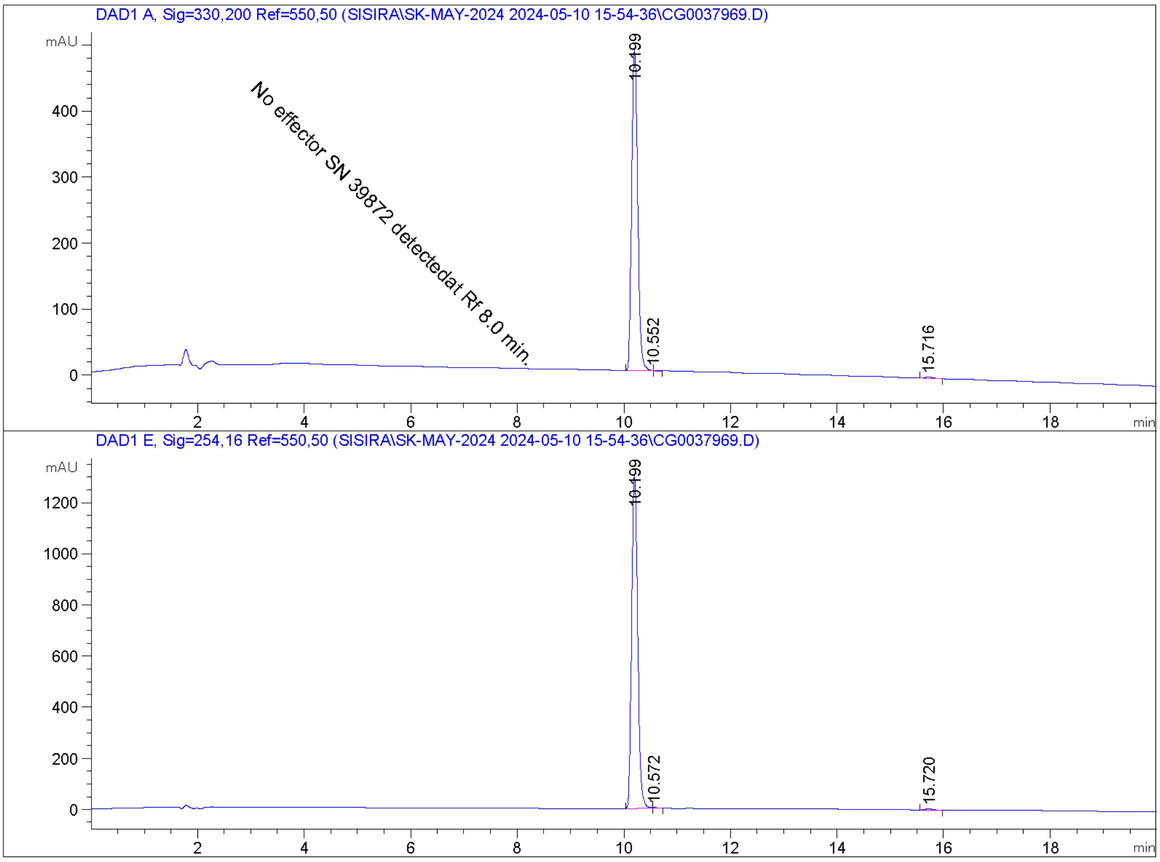


Signal 1: DAD1 A, Sig=330,200 Ref=550,50

| Peak # | RetTime [min] | Type | Width [min] | Area [mAU*s] | Height [mAU] | Area % |
|---|---|---|---|---|---|---|
| 1 | 10.199 | MF | 0.1281 | 3765.58008 | 489.95670 | 99.2744 |
| 2 | 10.552 | FM | 0.1157 | 7.20179 | 1.03752 | 0.1899 |
| 3 | 15.716 | BB | 0.1497 | 20.32187 | 2.08511 | 0.5358 |

Totals : 3793.10374 493.07933
